# RNF5 Regulation of RBBP4 Defines Acute Myeloid Leukemia Growth and Susceptibility to Histone Deacetylase Inhibitors

**DOI:** 10.1101/2020.10.25.349241

**Authors:** Ali Khateb, Anagha Deshpande, Yongmei Feng, Joo Sang Lee, Ikrame Lazar, Bertrand Fabre, Yan Li, Darren Finlay, Yu Fujita, Tongwu Zhang, Jun Yin, Ian Pass, Ido Livneh, Carol Burian, James R. Mason, Ronit Almog, Nurit Horesh, Yishai Ofran, Kevin Brown, Kristiina Vuori, Michael Jackson, Eytan Ruppin, Aniruddha J. Deshpande, Ze’ev A. Ronai

**Affiliations:** Technion Integrated Cancer Center, Faculty of Medicine, Technion Israel Institute of Technology, Haifa 31096, Israel; Cancer Center, Sanford Burnham Prebys Medical Discovery Institute, La Jolla, CA 92037, USA; Cancer Data Science Lab (CDSL), National Cancer Institute, National Institute of Health, Bethesda, MD 20892, USA; Laboratory of Translational Genomics, Division of Cancer Epidemiology and Genetics, National Cancer Institute, Bethesda, MD 20892, USA; Scripps MD Anderson Cancer Center, La Jolla, CA 92121, USA; Rambam Health Care Campus, Epidemiology Department and Biobank, Haifa 31096, Israel; Rambam Health Care Campus, Hematology and Bone marrow Transplantation Department, Haifa 31096, Israel

**Keywords:** AML, HDAC, RBBP4, RNF5, ubiquitin ligase

## Abstract

Acute myeloid leukemia (AML) remains incurable, largely due to its resistance to conventional treatments. Here, we found that increased expression and abundance of the ubiquitin ligase RNF5 contributes to AML development and survival. High RNF5 expression in AML patients correlated with poor prognosis. RNF5 inhibition decreased AML cell growth in culture and *in vivo*, and blocked development of MLL-AF9–driven leukemogenesis in mice, prolonging their survival. RNF5 inhibition led to transcriptional changes that overlapped with those seen upon HDAC1 inhibition. RNF5 induced the formation of K29 ubiquitin chains on the histone-binding protein RBBP4, promoting its recruitment and subsequent epigenetic regulation of genes involved in AML development and maintenance. Correspondingly, RNF5 or RBBP4 knockdown enhanced the sensitivity of AML cells to histone deacetylase (HDAC) inhibitors. Notably, low expression of *RNF5* and *HDAC* coincided with a favorable prognosis. Our studies identified ERAD-independent role for RNF5, demonstrating that its control of RBBP4 constitutes an epigenetic pathway that drives AML while highlighting RNF5/RBBP4 as markers to stratify patients for treatment with HDAC inhibitors.

## INTRODUCTION

AML is a heterogeneous hematological cancer characterized by the accumulation of somatic mutations in immature myeloid progenitor cells. The mutations alter the self-renewal, proliferation, and differentiation capabilities of the progenitor cells ^1, 2^. The prognosis of AML patients is strongly influenced by the type of chromosomal or genetic alterations and by changes in gene expression ^1, 3^. Although numerous mutations and chromosomal aberrations that drive AML development have been identified ^1, 4^, the molecular components and epigenetic modulators that contribute to the etiology and pathophysiology of AML are not well defined. Approximately one third of AML patients fail to achieve complete remission in response to chemotherapy, and 40 – 70% of those who do enter remission relapse within 5 years. Thus, there is an urgent need to better understand the molecular mechanisms underlying AML development and progression to facilitate the development of more effective therapies.

RING finger protein 5 (RNF5) is an ER-associated E3 ubiquitin ligase and a component of the UBC6e-p97 complex, which has been implicated in ER-associated degradation (ERAD) ^5, 6^, a pathway involved in maintaining protein homeostasis. RNF5 recognizes misfolded proteins and promotes their ubiquitination and proteasome-dependent degradation ^5, 6^. *RNF5* expression is increased in several cancers, including breast cancer, hepatocellular carcinoma, and AML ^7^. RNF5 regulates glutamine metabolism through degradation of misfolded glutamine carrier proteins, a function that is important in the response of cancer cells to ER stress-inducing chemotherapies such as paclitaxel ^8^. RNF5 promoted the degradation of the protease ATG4B, which limits basal amounts of autophagy ^9^. RNF5 also limits intestinal inflammation by its control of S100A8 protein stability ^10^. Given the important pathophysiological roles of RNF5, the observation that it is upregulated in AML cells and patient samples prompted us to investigate the possible contribution of RNF5 to the development and progression of this disease.

Histone modification by acetylation contributes to the dynamic regulation of chromatin structure and affects gene expression programs. Histone acetylation status is associated with the transcriptional regulation of leukemic fusion proteins, such as AML1-ETO, PML-RARα, and MLL-CBP ^11, 12^. Correspondingly, histone deacetylases (HDACs) are implicated in the etiology and progression of leukemia ^13^, and HDAC inhibitors affect the growth, differentiation, and apoptosis of leukemia cells ^14^. The retinoblastoma binding protein 4 (RBBP4) is a component of multi-protein complexes involved in nucleosome assembly and histone modifications, which influences gene transcription and regulates cell cycle and proliferation ^15^. Such complexes include nucleosome remodeling and deacetylase (NuRD) complex, polycomb repressor complex 2 (PRC2), and switch independent 3A (SIN3A) ^15, 16^. Overexpression of *RBBP4* and *HDAC1* correlates with clinicopathologic characteristics and prognosis in breast cancer ^17^, and *RBBP4* expression correlates with hepatic metastasis and poor prognosis in colon cancer patients ^18^. RBBP4 was also implicated in the regulation of DNA repair genes and its suppression in glioblastoma enhanced the sensitivity to temozolomide chemotherapy ^19^. However, the function of RBBP4 in AML has not been studied.

Here, we identified a central role for the RNF5-RBBP4 axis in AML maintenance and responsiveness to HDAC inhibitors. Our data suggest that targeting RNF5 and HDAC pathways represents a new therapeutic modality to inhibit AML and that expression of *RNF5* could serve as a prognostic marker and means to stratify patients for treatment with HDAC inhibitors.

## RESULTS

### Increased expression of RNF5 in AML patients correlates with poor prognosis

Analysis of RNA-seq datasets for various cancer cells from the Cancer Cell Line Encyclopedia database identified higher amounts of *RNF5* transcripts in AML, chronic myeloid leukemia (CML), and T-cell acute lymphoblastic leukemia (T-ALL), compared with other tumor types (Extended Data Fig. 1A). Higher amounts of RNF5 protein were confirmed in AML and CML, compared with other tumor types (Extended Data Fig. 1B). To assess the clinical relevance of RNF5 in AML, we analyzed the amount of RNF5 in peripheral blood mononuclear cells (PBMCs) from independent cohorts of AML patients. Similar to the AML cell lines, the average of RNF5 abundance was significantly higher in PBMCs from AML patients compared with PMBCs from healthy subjects (Fig. 1A,B). Stratification of the 50 patients into two groups based on high (N = 8, 15%) and low (n = 42, 85%) revealed that high level of RNF5 abundance coincided with poor overall survival (*P* = 0.05, Fig. 1C). An independent analysis of AML patients (n = 154) from The Cancer Genome Atlas (TCGA) dataset confirmed a significant correlation between high *RNF5* expression (10%) and poor survival (*P* = 0.009, Fig. 1D). Notably, AML patients with or without *FLT3* or *NPM1* mutations did not exhibit any difference in *RNF5* expression (Extended Data Fig. 1C,D), suggesting that the importance of RNF5 in AML depends on select oncogenic driver(s) and the activation of related signaling pathways.

**Fig. 1:**
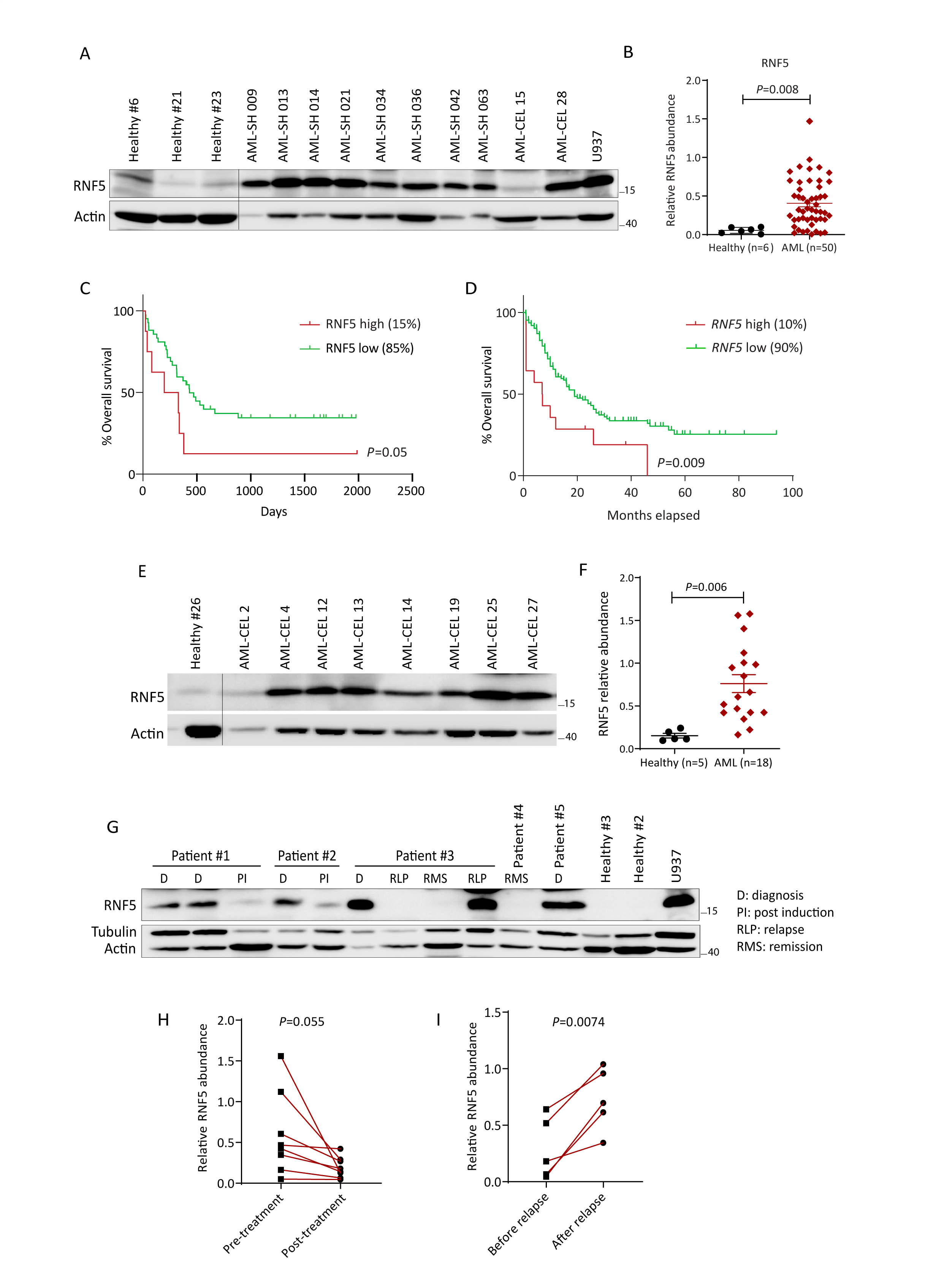
RNF5 expression in AML patient samples. **(A)** Representative western blot analysis of RNF5 in peripheral blood mononuclear cells (PBMCs) from 3 healthy control subjects and 10 AML patients (Scripps Health cohort). **(B)** Relative RNF5 abundance in AML samples (n = 50) compared with healthy control subjects (n = 6) from the Scripps Health Center. Data were quantified relative to RNF5 abundance in U937 cells that was used as a reference. Quantified data are presented as the mean ± SEM. *P* = 0.008 by unpaired two-tailed *t*-test. **(C)** Kaplan–Meier survival curve of AML patients stratified by top high (n = 8) versus low (n = 42) RNF5 abundance (Scripps Health cohort). *P* = 0.05 by Mantel–Cox log-rank test. **(D)** Kaplan–Meier survival curve of AML patients stratified by top high (n = 14) versus low (n = 150) *RNF5* transcript levels (TCGA dataset). *P* = 0.009 by Mantel– Cox log-rank test. **(E-G)** Abundance of RNF5 and control proteins in PBMCs from healthy control subjects and AML patients from the Rambam Health Campus Center cohort. Quantified data are presented as the mean ± SEM. *P* = 0.006 by unpaired two-tailed *t*-test. **(H, I)** Relative abundance of RNF5 protein in AML patient samples collected before and after induction treatment (H) or before and after relapse (I). Lines connect values for the same patient. \*\**P* = 0.0066, \**P* = 0.054, \*\**P* = 0.0074 by unpaired (B and F) or paired (H and I) two-tailed *t*-test.

Assessment of an independent AML patient cohort (Rambam Health Campus Center, Haifa, Israel) corroborated the higher level of RNF5 in AML patients (n = 18), compared with healthy donors (n = 5) (Fig. 1E,F). Because this cohort included multiple samples from the same patient, that were obtained prior to and following therapy, we could monitor changes in RNF5 abundance in AML patients that were subjected to therapy and at remission and relapse stages. Notably, RNF5 abundance markedly decreased following chemotherapy and during remission (n = 8) (Fig. 1G,H and Extended Data Fig. 1E). Conversely, the amount of RNF5 was similar to that observed at diagnosis in patients that either relapsed or were refractory to treatment (n = 5) (Fig. 1I, J). These results suggested that the amount of RNF5 may serve as a prognostic marker for AML.

### RNF5 is required for AML cell proliferation and survival

Discovering high amounts of RNF5 in AML, CML, and T-ALL cell lines, coupled with the link to AML progression, led us to explore the impact of RNF5 knockdown (RNF5-KD) on leukemia cell growth. Interestingly, RNF5-KD using RNF5-targeting short hairpin RNAs (shRNF5) reduced the viability and attenuated the growth of the AML cell lines MOLM-13 and U937 (Fig. 2A and Extended Data Fig. 2A) but not of CML cells (K-562) or T-ALL cells (Jurkat) (Extended Data Fig. 2B,C). RNF5-KD in MOLM-13 and U937 AML cells also caused the accumulation of cells in the G1 phase of the cell cycle (Fig. 2B), which was accompanied by an increase in the cell cycle regulatory proteins p27 and p21 (Fig. 2C). Moreover, RNF5-AML cells reduced colony formation in soft agar (Fig. 2D) and increased the abundance of proteins associated with apoptosis reflected in the level of cleaved forms of caspase-3 (Fig. 2E) and poly ADP ribose polymerase (PARP) (Extended Data Fig. 2D). The effects of RNF5-KD on U937 and MOLM-13 cells were corroborated in two additional AML cell lines, HL-60 and THP-1 (Extended Data Fig. 2E–H). Importantly, re-expression of RNF5 restored cell proliferation and reduced the apoptosis-associated proteins in RNF5-KD cells (Fig. 2F,G), substantiating the role of RNF5 in the control of cell cycle and death programs in AML cells.

**Fig. 2:**
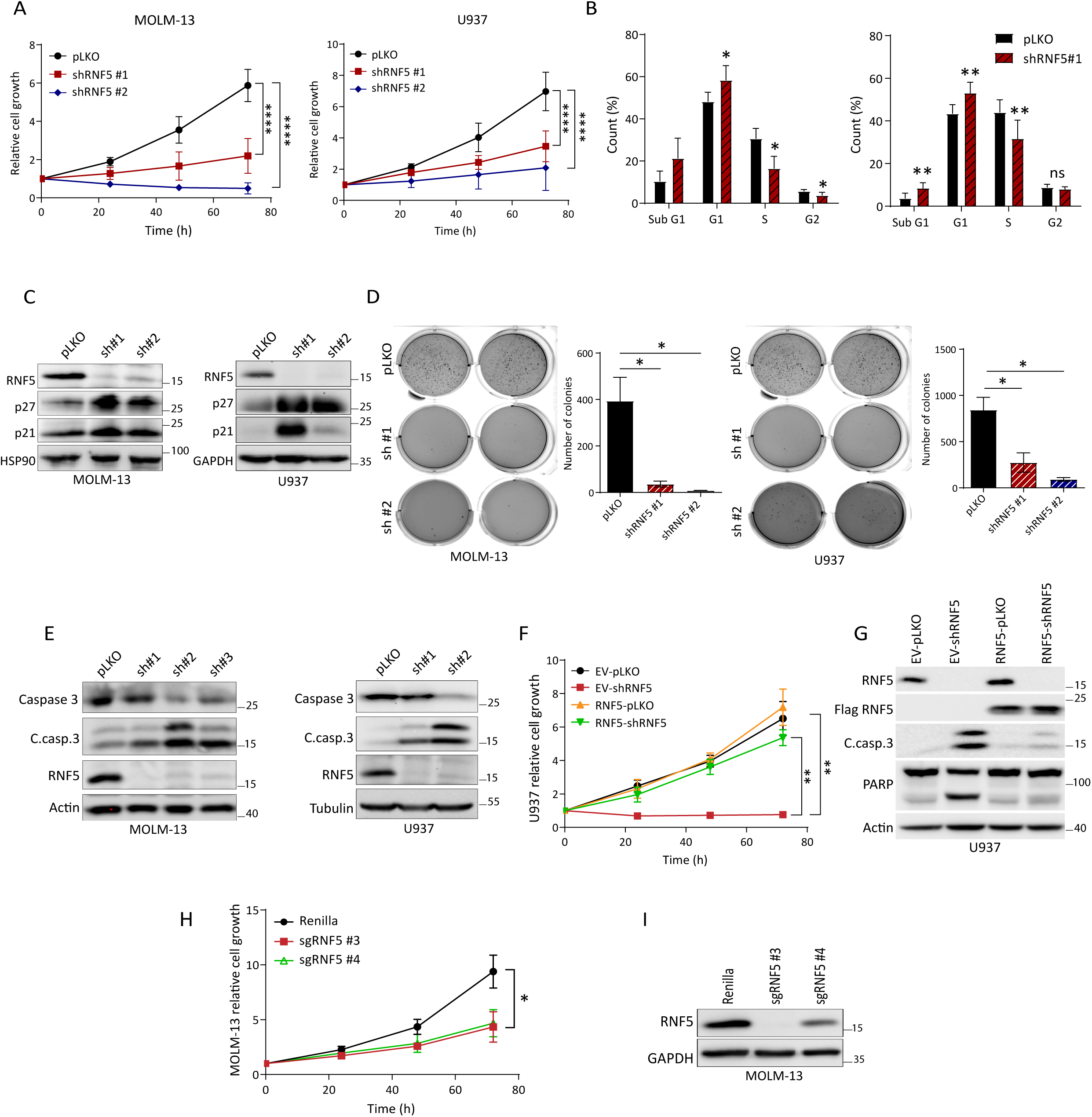
RNF5 is required for AML cell proliferation and survival. (A) Growth assay of MOLM-13 and U937 cells after transduction with empty vector (pLKO) or two different shRNF5 constructs. (B) Cell cycle analysis of MOLM-13 and U937 cell lines 5 days after transduction. (C) Western blot analysis of cell cycle regulatory proteins in MOLM-13 and U937 AML cells 5 days after transduction. (D) Representative images (left) and quantification (right) of MOLM-13 and U937 colonies in soft agar assessed after 15 days in culture. (E) Western blot analysis of apoptosis-related proteins in MOLM-13 and U937 AML cells 5 days after transduction (C.casp.3 = cleaved caspase-3). (F) Growth assay of U937 cells expressing doxycycline-inducible Flag-tagged RNF5 or empty vector (EV). Cells were induced with doxycycline (1 µg/ml) for 48 h and then transduced with empty vector (pLKO) or shRNF5 for 5 days. (G) Western blot analysis of the indicated proteins in U937 infected as described in (F). (H) Growth assay of MOLM-13 cells stably expressing Cas9 and transduced with control *Renilla*-targeting sgRNA or two RNF5-targeting sgRNAs. CRIPSR was performed based on CRIPSR knockout cell pool. (I) Western blot analysis of the indicated proteins in MOLM-13 cells as described in (H). Quantified data are presented as the mean ± SD of three independent experiments. Western blot data are representative of three experiments. \**P* < 0.05, \*\**P* < 0.01, and \*\*\*\**P* < 0.0001 by two-way ANOVA followed by Tukey’s multiple comparison test (A, F, and H) or two-tailed *t*-test (B, and D).

To verify the role of RNF5 in AML cells, we used the CRISPR-Cas9 editing technology to deplete RNF5. Indeed, impaired growth of MOLM-13 AML cells stably expressing Cas9 and transduced with RNF5-targeting guide RNAs (sgRNAs) was seen, compared with control cells transduced with *Renilla* luciferase-targeting sgRNAs (Fig. 2H,I). These data substantiated our observations with shRNA-mediated knockdown and supported a role for RNF5 in AML cell proliferation.

Despite detecting high amounts of the ubiquitin ligase RNF5 in many AML, CML, and T-ALL cell lines, our data suggested that RNF5 was specifically important for the survival and growth of AML cells. A cell- or tissue-specific role for RNF5 is supported by studies in breast cancer and melanoma, in which RNF5-KD or overexpression elicits distinct outcomes: RNF5 overexpression promotes the growth of melanoma by affecting the tumor microenvironment, whereas RNF5 overexpression inhibits breast cancer growth through tumor-intrinsic effects ^7, 20^.

### ER stress-induced apoptosis in AML cells is enhanced upon RNF5 inhibition and attenuated by RNF5 overexpression

Because RNF5 is part of ERAD and the ER stress response, we assessed whether modulation of RNF5 abundance alters the ER stress response in AML cells. We exposed MOLM-13 cells to thapsigargin or tunicamycin to inhibit the ER Ca^2+^-ATPase (SERCA) or protein glycosylation, respectively ^21^, and induce ER stress. Both agents caused a greater increase in the abundance of apoptotic markers in RNF5-KD MOLM-13 cells than in control cells (Fig. 3A,B), and thapsigargin also decreased viability to a greater extent in RNF5-KD MOLM-13 cells (Fig. 3C). Tunicamycin also reduced viability of RNF5-KD HL-60 cells (Extended Data Fig. 3A). Consistent with a role in ER stress, RNF5-KD resulted in increased transcripts for key UPR components, including CHOP, ATF3, and sXBP1, in thapsigargin-treated MOLM-13 (Fig. 3D) and HL-60 cells (Extended Data Fig. 3B).

**Fig. 3:**
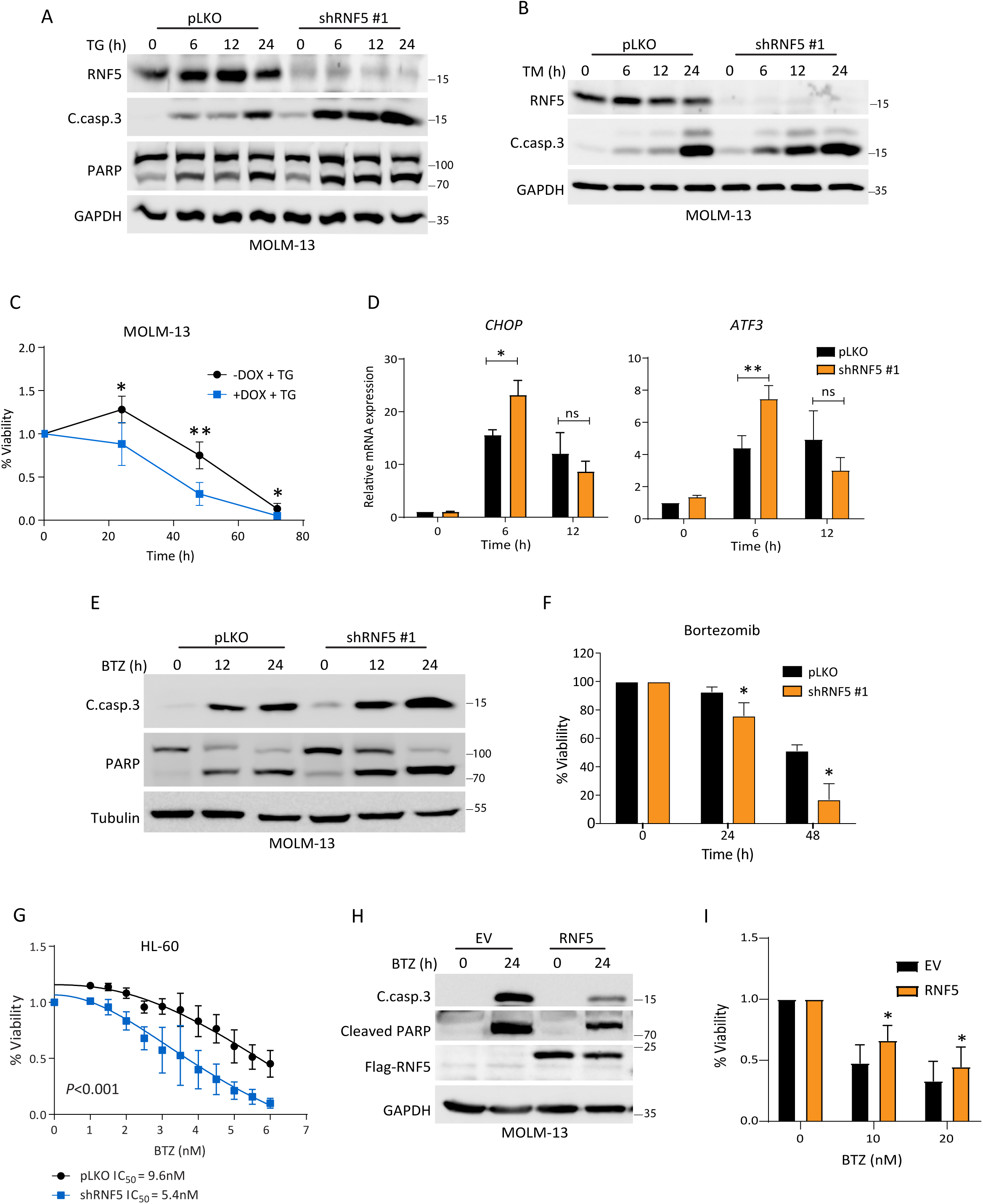
RNF5 inhibition sensitizes AML cells to ER stress-induced apoptosis. **(A, B)** Western blot analysis of the indicated proteins in MOLM-13 cells expressing empty vector (pLKO) or shRNF5 #1 and treated with thapsigargin (TG, 1 µM) (A) or tunicamycin (TM, 2 µg/ml) (B) for the indicated times. **(C)** Luminescence viability assay of MOLM-13 cells expressing inducible shRNF5 treated with or without doxycycline (DOX, 1 µg/ml) for 3 days before treatment with TG (100 nM) for the indicated times. **(D)** RT-qPCR analysis of *CHOP* and *ATF3* mRNA in MOLM-13 cells expressing empty vector (pLKO) or shRNF5 #1 and treated with TG (100 nM) for the indicated times. **(E)** Western blot analysis of cleaved caspase-3 (C. casp.3) and PARP in MOLM-13 cells expressing empty vector (pLKO) or shRNF5 #1 and treated with bortezomib (BTZ, 5 nM) for the indicated times. **(F)** Fluorescence viability assay of HL-60 cells expressing empty vector (pLKO) or shRNF5 and treated with BTZ (5 nM) for the indicated times. Cell viability was determined by flow cytometry of cells stained with annexin V conjugated to fluorescein isothiocyanate and propidium iodide. **(G)** Luminescence viability assay of HL-60 cells expressing empty vector (pLKO) or shRNF5 and treated with the indicated concentrations of BTZ for 48 h. **(H)** Western blot analysis of MOLM-13 cells expressing inducible empty vector (EV) or an RNF5 overexpression vector. Cells were first induced with DOX (1 µg/ml) for 24 h and then treated with BTZ (5 nM) for the indicated times. **(I)** Luminescence viability assay of MOLM-13 cells expressing DOX-inducible Flag-tagged RNF5 or empty vector (EV). Cells were treated with DOX (1 µg/ml) for 2 days and then treated with the indicated concentrations of BTZ for 12 h. Data are presented as the mean ± SD of three independent experiments. \**P* < 0.05 and \*\**P* < 0.01 by two-tailed *t*-test (C, D, and F) or by two-way ANOVA (G). ns: not significant.

Given the close link between ER stress and proteasomal degradation, we assessed potential synergy between knockdown of RNF5 and inhibition of the proteasomes. Indeed, RNF5-KD MOLM-13 cells treated with the proteasome inhibitor bortezomib (BTZ) exhibited increased amounts of apoptotic markers (Fig. 3E) and decreased viability (Fig. 3F), compared with control cells. Using annexin V and propidium iodide staining, which monitor degree of programmed cell death, we showed that RNF5-KD enhanced apoptosis of HL-60 cells exposed to BTZ (Fig. 3G), reducing the inhibitory concentration at which 50% of cells died (IC_50_) from 9.6 nM in control cells to 5.4 nM in RNF5-KD cells. Overexpression of Flag-tagged RNF5 partially protected AML cells from tunicamycin or BTZ-induced activation of apoptosis (Fig. 3H,I, Extended Data Fig. 3C,D), confirming the role of RNF5 in protecting AML cells from proteotoxic stress.

### RNF5 loss delays leukemia establishment and progression

Our findings with cultured AML cells prompted us to examine the role of RNF5 in leukemia growth *in vivo*. We used the human AML xenograft model in which the luciferase-expressing U937 cells (U937-pGFL) were transduced with doxycycline-inducible shRNF5 before the cells were injected intravenously into NOD/SCID mice (Extended Data Fig. 4). Following leukemia establishment, confirmed by bioluminescence, mice were fed a doxycycline-containing diet and monitored for disease progression and overall survival. RNF5-KD markedly decreased leukemia burden resulting in prolonged survival, compared with control mice (Fig. 4A,B). These data indicated that RNF5 is also required for proliferation of AML cells *in vivo*.

**Fig. 4:**
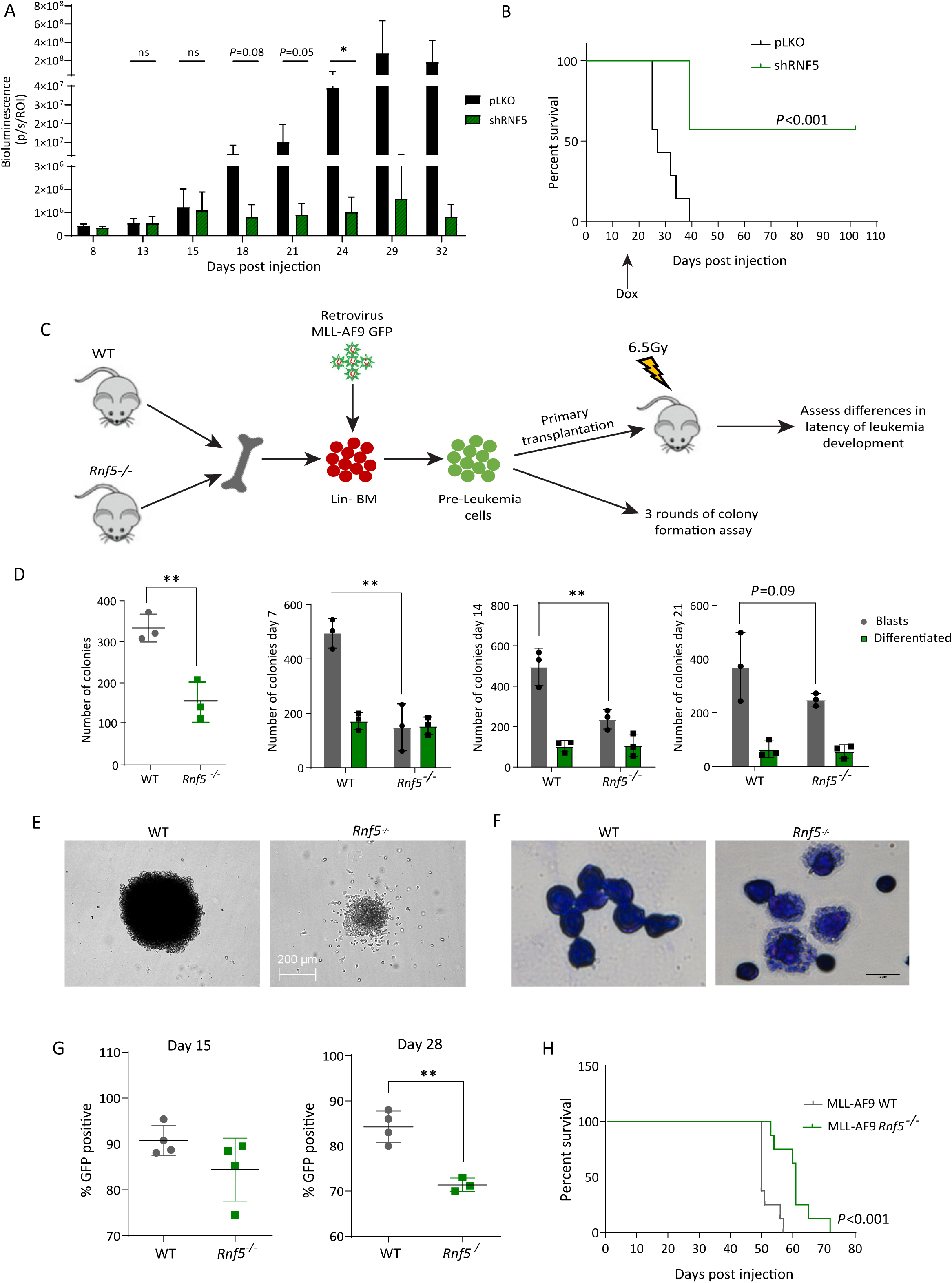
RNF5 suppression impairs leukemia establishment and progression *in vivo*. (A) Graph depicting growth in mice of U937-pGFL expressing empty vector (pLKO) or inducible shRNF5. Bioluminescence was quantified as measure of disease burden. Data are presented as the mean ± SD of 7 mice per group. \**P* < 0.05 by unpaired two-tailed *t*-test. (B) Kaplan–Meier survival curves of mice injected with U937-pGFL cells expressing empty vector (pLKO) (n = 7 mice/group) or inducible shRNF5 (n = 7 mice/group). *P* < 0.001 by Mantel–Cox log-rank test. (C) Schematic representation of the experiment. Lin^−^Sca1^+^c-Kit^+^ (LSK) cells were purified from the bone marrow of WT or *Rnf5^−/–^* mice, transduced *in vitro* with a GFP-tagged MLL-AF9 fusion gene, and then either analyzed in colony-forming assays *in vitro* or intravenously injected into sub lethally irradiated WT C57BL/6 mice. (D) Quantification of total colonies (left) or blast-like and differentiated colonies (right) of GFP-MLL-AF9–transformed WT or *Rnf5^−/–^*cells after 7 days, 14, or 21 days in culture. Data are presented as the mean ± SD of three independent experiments. \*\**P* < 0.01 by paired two-tailed *t*-test. (E) Representative pictures of colonies of GFP-MLL-AF9–transformed WT or *Rnf5^−/–^*cells after 7 days in culture. Scale bar 200 µm. (F) Wright–Giemsa-staining of GFP-MLL-AF9–transformed WT or *Rnf5^−/–^* cells after 7 days in culture. (G) Flow cytometric quantification of GFP+ cells in the peripheral blood of mice intravenously injected with GFP-MLL-AF9–transformed WT (n = 4) or *Rnf5*^−/–^ (n = 4) cells at days 15 and 28 post-injection. Data are presented as the mean ± SD. \*\**P* < 0.01 by unpaired two-tailed *t*-test. (H) Kaplan–Meier survival curves of mice injected with GFP-MLL-AF9–transformed WT or *Rnf5^−/–^* cells. Data are from two independent experiments (n = 4 mice/group per experiment). *P* < 0.001 by Mantel–Cox log-rank test.

We also assessed whether RNF5 has a role in AML initiation using the MLL-AF9 model ^22^ for *in vitro* and *in vivo* studies. Hematopoietic stem and progenitor (Lin-depleted) cells (HSPCs) were purified from the bone marrow of wild-type (WT) or *Rnf5^−/−^* C57/BL6 mice and retrovirally transduced with a bicistronic construct harboring MLL-AF9 and a green fluorescent protein (GFP) marker. We then assessed the colony-forming ability of these pre-leukemic cells *in vitro*. Compared with WT GFP-MLL-AF9 cells, *Rnf5^−/–^* GFP-MLL-AF9 cells exhibited markedly reduced colony-forming capacity in methylcellulose after 7, 14, and 21 days in culture, with a striking reduction in the number of blast-like colonies (Fig. 4C). The reduced size and number of cells in these colonies was consistent with terminal differentiation of *Rnf5^−/−^* cells, reflected by a greater cytoplasm/nuclei ratio and more vacuolated cytoplasm (Fig. 4D,E).

To assess leukemogenesis *in vivo*, sublethally irradiated WT C57/BL6 recipient mice were injected with GFP-MLL-AF9–transduced *Rnf5*^WT^ or *Rnf5*^−/–^ cells, and cell engraftment was monitored by flow cytometry of GFP-positive (GFP+) cells in peripheral blood (Fig. 4F). Analysis on days 15 and 28 post-injection identified fewer GFP+ cells in mice injected with GFP-MLL-AF9 *Rnf5*^−/–^ cells, compared with mice injected with GFP-MLL-AF9 *Rnf5*^WT^ cells, indicating a delay in leukemia development (Fig. 4G). Moreover, mice harboring GFP-MLL-AF9 *Rnf5*^−/–^ cells exhibited prolonged survival compared with mice injected with GFP-MLL-AF9 *Rnf5*^WT^ cells (Fig. 4H). Collectively, these data showed that RNF5 loss promotes terminal differentiation and decreases the colony-forming ability of MLL-AF9–transformed pre-leukemic cells *in vitro* and delays leukemia progression *in vivo*.

### RNF5 affects gene transcription in AML cells

To assess possible pathways affected by RNF5 in AML cells, we first monitored transcriptional changes in MOLM-13, U937, and HL-60 AML cell lines expressing RNF5-WT or subjected to RNF5-KD. RNA sequencing (RNA-seq) analysis identified a differentially expressed gene set in RNF5 KD AML, compared with control (RNF5 WT) cells (Fig. 5A, Extended Data Fig. 5A and Supplemental Table 1). Ingenuity Pathway Analysis identified selective enrichment of genes implicated in myeloid cell function, such as NF-κB signaling, IL-8 signaling, reactive oxygen species, and several pathways related to cell migration such as Rho GTPases and Tec kinase signaling (Fig. 5B). Among the genes upregulated by RNF5-KD in all three AML cell lines were *CDKN1A* and *CDKN2D* encoding cell cycle inhibitors, *LIMK1* encoding a kinase involved in regulation of the actin cytoskeleton, *ANXA1* encoding a calcium-binding protein involved in metabolism, EGFR and cell death programs, and *NCF1* encoding a subunit of NADPH oxidase (Fig. 5C and Extended Data Fig. 5B). These regulatory pathways and the transcriptional changes are consistent with the phenotypic changes of reduced proliferation, increased apoptosis, and increased differentiation that we observed in RNF5-KD AML cell lines. Interestingly, analysis of the Library of Integrated Network-Based Cellular Signatures (LINCS) drug screening database identified a notable overlap between the transcriptomic changes induced by the HDAC1 inhibitor mocetinostat in different cancer cells, to those seen in shRNF5 treated MOLM-13 and HL-60 cells (Fig. 5D). Five out of top ten transcriptional changes identified in LINCS following HDAC inhibition overlapped with those seen following shRNF5 in MOLM-13 cells (Fig. 5D and Extended Data Fig. 5C). Among the commonly affected pathways were activation of GP6 and Rho GTPase signaling, and repression of the nucleotide excision repair (NER) pathway (Fig. 5E). These observations point to the possibility that HDAC may underlie RNF5 role in AML.

**Fig. 5:**
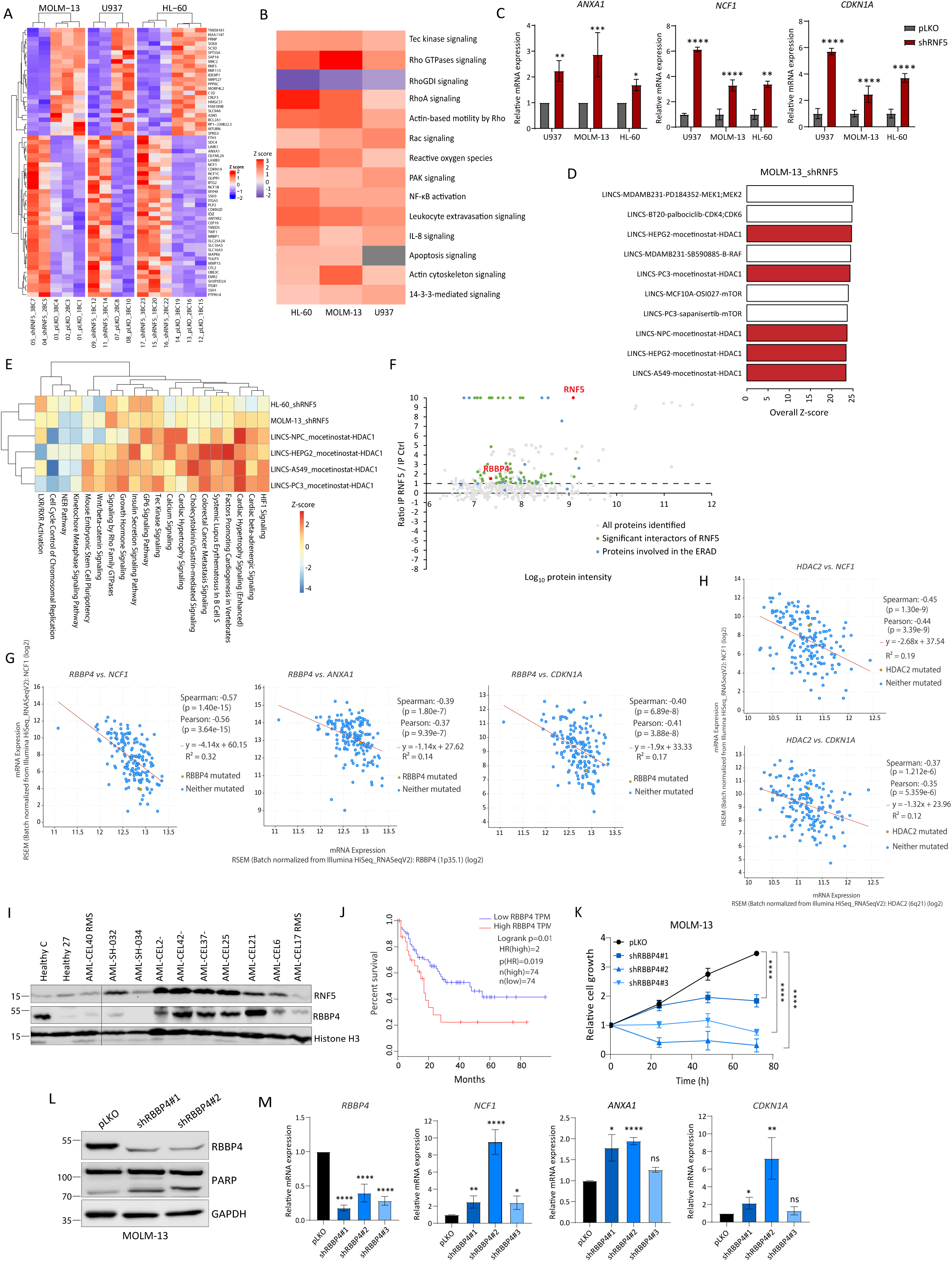
Transcriptional and survival outcome overlap for RNF5 and RBBP4 in AML. (A) Heatmap of RNA-seq data performed on control (pLKO) or RNF5-KD (shRNF5) AML cell lines. Genes with Benjamini-Hochberg (BH) corrected p value< 0.05 and Log2 transformed fold change >=0.4 or <=-0.4 were selected as significantly differentially expressed genes. (B) Top canonical pathways identified by Ingenuity Pathway Analysis comparing the differentially expressed genes in AML cell lines (HL-60, U937, and MOLM-13) upon RNF5-KD. (C) RT-qPCR validation of indicated genes identified as deregulated by RNF5-KD. Data are presented as the mean ± SD of three independent experiments. (D) Top ten drug screening results from the Library of Integrated Network-Based Cellular Signatures (LINCS) matched by shRNF5 in MOLM-13 cell line. Values are overall z-scores from IPA Analysis Match database. HDAC1 inhibitor results are shown in red. (E) shRNF5 in MOLM-13 and HL-60 cell lines induced similar IPA Canonical Pathway changes compared with NPC, HEPG2, A549 and PC3 cancer cell lines treated with the HDAC1 inhibitor mocetinostat. Z-score was calculated by IPA, with positive z-score predicting activation of the pathway, and negative z-score predicting inhibition of the pathway. (F) Log_2_-transformed ratio of proteins in anti-Flag immunoprecipitates of RNF5-overexpressing versus control cells determined by MS intensity. Green indicates proteins significantly enriched in RNF5-overexpressing cells; blue indicates proteins in the ERAD pathway enriched in RNF5-overexpressing cells. RNF5 and RBBP4 are indicated in red. (G, H) Co-expression of *RBBP4* (G) or *HDAC2* (H) and the indicated RNF5 target genes in AML analyzed in cBioPortal using the TCGA database. (I) Western blot analysis of RBBP4 in PBMCs from healthy control subjects and AML patients from Scripps Health and Rambam Health Campus Center cohorts. RMS stands for remission. (J) Overall survival rate of AML patients expressing high (30%) or low (70%) *RBBP4* transcripts. Survival analysis was performed by the bioinformatics database GEPIA using TCGA tumor samples. TPM: Transcripts Per Million. HR: hazard ratio. (K) Growth assay of MOLM-13 cells after transduction with empty vector (pLKO) or the indicated shRBBP4 constructs. (L) Western blot analysis of the indicated proteins in MOLM-13 cells expressing empty vector (pLKO) or two different shRBBP4 constructs. (M) RT-qPCR analysis of indicated genes identified as deregulated by RNF5-KD in MOLM-13 cells expressing the indicated constructs. Data are presented as the mean ± SD of three independent experiments. \**P* < 0.05, \*\**P* < 0.01, \*\*\**P* < 0.001, and \*\*\*\**P* < 0.0001 by two-tailed *t*-test (C and M) or by two-way ANOVA followed by Tukey’s multiple comparison test (K).

### RNF5 interacts with and ubiquitinates retinoblastoma binding protein 4

We hypothesized that RNF5 elicited transcriptional changes through intermediate regulatory component(s). Therefore, we performed liquid chromatography–tandem mass spectrometry (LC-MS/MS) to identify RNF5-interacting proteins, which may also include RNF5 substrates that mediate the transcriptional and phenotypic changes seen upon RNF5-KD. We compared the proteins isolated from MOLM-13 cells expressing inducible Flag-tagged RNF5 or empty vector by immunoprecipitation with an antibody recognizing Flag-RNF5 (Extended Data Fig. 5D,E). Among the 65 RNF5-interacting proteins that we identified were previously reported substrates, such as 26S proteasome components, VCP and S100A8 ^5, 10^, as well as proteins implicated in AML development, such as DHX15 ^23^ and gelsolin ^24, 25^ (Supplemental Table 2). Among the more abundant RNF5-bound proteins were components of the ERAD, translation initiation, proteolysis, and mRNA catabolic processes (Fig. 5F, Extended Data Fig. 5D,E and Supplemental Table 2).

Although none of the interacting proteins were transcription factors, epigenetic modification upon altered RNF5 expression may underlie changes observed in gene expression. Thus, we assessed whether RNF5-interacting proteins include epigenetic regulators, identifying the histone binding protein RBBP4 (Fig. 5F and Supplemental Table 2). Analysis of transcriptome data from TCGA revealed that expression of *RBBP4* was inversely correlated with the expression of genes that were increased in RNF5-KD cells (Fig. 5G and Extended Data Fig. 5F), which implies that RNF5 positively controls the transcriptional regulatory function of RBBP4. RBBP4 is a component of chromatin assembly, remodeling, and nucleosome modification complexes, including PRC2 ^26^ and the corepressor NuRD complex which contains HDAC1 and HDAC2 ^27^. Indeed, such correlation between *RBBP4* and RNF5-upregulated genes was also seen for *HDAC1*, *HDAC2* and *EZH2* (Fig. 5H and Extended Data Fig. 5G-I). Increased *RBBP4* expression is correlated with the malignant phenotypes in human tumors including AML ^28^. Analysis of tumor data in TCGA revealed high expression of *RBBP4* in AML compared with other tumor types (Extended Data Fig. 5J). Assessment of AML patient cohort confirmed higher RBBP4 expression in AML patients, compared with healthy donors (Fig. 5I). Stratifying AML patients by *RBBP4* expression showed that high expression correlated with poor overall survival (Fig. 5J).

If RNF5 is a positive regulator of RBBP4, we expected that RBBP4-KD would result in similar phenotypic changes to AML cells subjected to RNF5-KD. Indeed, knock down of RBBP4 in MOLM-13 and U937 cells with shRNAs impaired their growth (Fig. 5K and Extended Data Fig. 5K), exhibited PARP cleavage indicating apoptosis (Fig. 5L and Extended Data Fig. 5L) and showed induction of genes induced by RNF5-KD (Fig. 5M). These observations further substantiate a possible link between RNF5 and RBBP4.

RNF5 is a transmembrane protein primarily associated with the ER with the ubiquitin ligase domain in the cytosol ^6, 29^. We assessed a possible interaction between RNF5 and RBBP4 in HEK293T and MOLM-13 cell lines by coimmunoprecipitation of ectopically expressed WT, catalytically inactive RING mutant (RNF5 RM), or C-terminal transmembrane domain deletion mutant (RNF5 ΔCT) (Figure 6A). Endogenous RBBP4 coimmunoprecipitated with all RNF5 constructs, suggesting that both the RING and transmembrane domain are dispensable for this interaction (Fig. 6B,C). Next, we assessed potential effects of RNF5 on RBBP4 ubiquitination. Co-expression of HA-tagged ubiquitin, Myc-tagged RBBP4, and Flag-tagged RNF5 constructs, revealed that RBBP4 was ubiquitinated by WT RNF5, but not by RNF5 RM nor by RNF5 ΔCT (Fig. 6D), indicating that ubiquitin ligase activity (RING domain-dependent) and membrane association are required for RNF5-mediated increase in RBBP4 ubiquitination. Correspondingly, RNF5-KD in HEK293T or MOLM-13 cells decreased RBBP4 ubiquitination (Extended Data Fig, 6A,B).

**Fig. 6:**
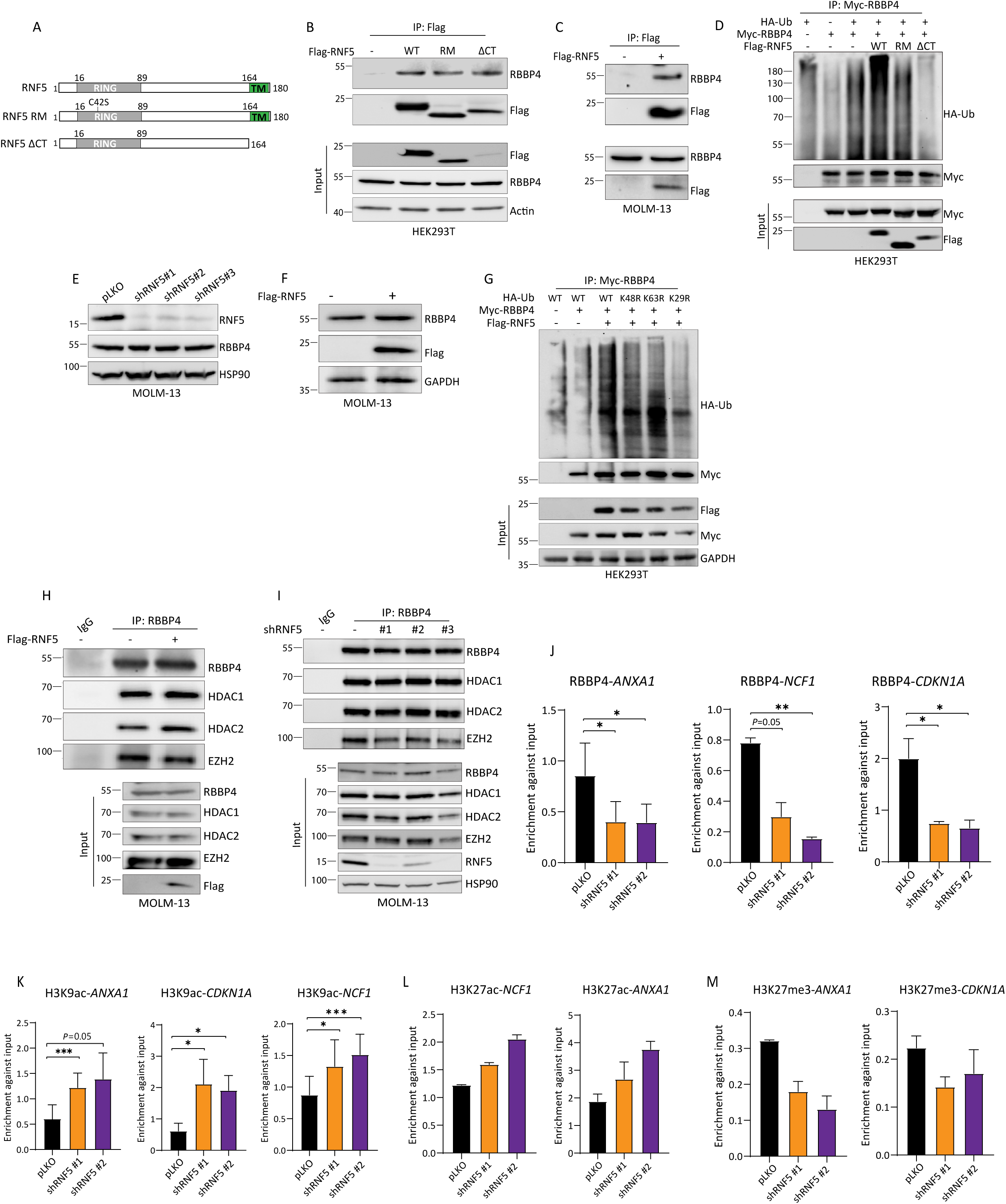
RBBP4 is RNF5 substrate, mediates epigenetic regulation of RNF5 target genes. (A) A schematic presentation of full-length RNF5 and its mutants. (B) Immunoprecipitation (IP) and Western blot analysis of HEK293T cells transfected with Flag-tagged full-length RNF5 (WT), a mutant with a catalytically inactive RING domain (RM), or a C-terminal transmembrane domain deletion mutant (ΔCT) and then treated with MG132 (10 µm 4 h) before lysis. (C) Immunoprecipitation and Western blot analysis of ectopically expressed RNF5 and endogenous RBBP4 in MOLM-13 cells expressing doxycycline-inducible Flag-tagged RNF5. Cells were incubated with or without doxycycline (1 µg/ml) for 2 days and with MG132 (10 µm 4 h) before lysis. (D) Western blot analysis of anti-Myc immunoprecipitates and lysates of HEK293T cells co-expressing Myc-RBBP4, hemagglutinin-tagged ubiquitin (HA-Ub), and the indicated Flag-tagged RNF5 constructs. Cells were treated with MG132 (10 µm 4 h) before lysis. (E) Western blot analysis of the indicated proteins in MOLM-13 cells expressing empty vector (pLKO) or the indicated shRNF5 construct. (F) Western blot analysis of the indicated proteins in MOLM-13 cells expressing empty vector or doxycycline-inducible Flag-tagged RNF5. (G) Western blot analysis of anti-Myc immunoprecipitates and lysates of HEK293T cells co-expressing Myc-RBBP4, Flag-tagged RNF5, and different HA-tagged ubiquitin constructs. WT, wild-type ubiquitin; K48R, K63R, K29R are lysine-to-arginine ubiquitin mutants at the indicated positions. Cells were treated with MG132 (10 µm 4 h) before lysis. (H) Immunoprecipitation and Western blot analysis of the interaction of RBBP4 with HDAC1, HDAC2, or EZH2 in MOLM-13 cells expressing doxycycline-inducible Flag-tagged RNF5. (I) Immunoprecipitation and Western blot analysis of the interaction of RBBP4 with HDAC1, HDAC2, or EZH2 in MOLM-13 cells expressing empty vector or the indicated shRNF5 construct. Cells were treated with MG132 (10 µm 4 h) before lysis. (J-M) ChIP and qPCR indicating the enrichment of RBBP4, H3K9ac, H3K27ac, or H3K27me3 (normalized to input) at the indicated gene promoters in MOLM-13 cells expressing empty vector or two shRNF5 constructs. Data in L and M are presented as the mean ± SD of two independent experiments. \**P* < 0.05 and \*\*\**P* < 0.001 by two-tailed *t*-test (J and K).

Notably, neither RNF5 overexpression nor RNF5-KD altered the abundance of RBBP4, suggesting that ubiquitination of RBBP4 by RNF5 is not through the formation of proteasome-targeting K48 ubiquitin chains and does not affect RBBP4 stability (Fig. 6E,F and Extended Data Fig. 6C). To assess the type of RBBP4 ubiquitination induced by RNF5, we used mutant HA-ubiquitin constructs, which disable ubiquitin chain formation on specific lysines (K48, K63, or K29), and monitored changes in Myc-tagged RBBP4 ubiquitination in cells expressing Flag-tagged RNF5. Only ubiquitin with the K29R mutation impaired RNF5-induced RBBP4 ubiquitination (Fig. 6G), suggesting that RNF5 induces K29-topology polyubiquitination of RBBP4.

### RNF5 promotes recruitment of RBBP4 to gene promoters

Because RBBP4 stability was not altered by RNF5, we evaluated whether RNF5 affects RBBP4 localization or interactions with other proteins. Subcellular fractionation and immunofluorescent analyses of nuclear and chromatin bound RBBP4 did not identify changes in RBBP4 localization following RNF5 inhibition (Extended Data Fig. 6E,F).

RBBP4 is a component of PRC2 and complexes containing HDACs ^30, 31^, thus, we assessed whether RNF5 affects the formation of these complexes or their recruitment to promoters of target genes. Neither overexpression nor KD of RNF5 affected the interaction of RBBP4 with HDAC1, HDAC2, or EZH2 (Fig. 6H,I and Extended Data Fig. 6G), suggesting that ubiquitination of RBBP4 by RNF5 is not required for the assembly of RBBP4-containing complexes. Chromatin immunoprecipitation (ChIP) and quantitative PCR (qPCR) assays were next used to evaluate the recruitment of RBBP4 to the promoters of genes that were regulated by either RNF5 or RBBP4. RNF5-KD reduced the recruitment of RPPB4 to the *ANXA1*, *NCF1*, and *CDKN1A* promotors (Fig. 6J). Examination of histone modifications in the promoters of these genes identified that RNF5-KD increased H3K9 and H3K27 acetylation (Fig. 6K and L) and reduced H3K27 methylation (Fig. 6M), changes that are indicative of increased gene expression ^32, 33^. These changes are consistent with their increased expression upon RNF5-KD (Fig. 5C). These results suggested that RNF5 control of gene expression in AML cells is mediated by RBBP4.

### RNF5 inhibition sensitizes AML cells to HDAC inhibitors

To provide independent support for the importance of the RNF5-RBBP4 regulatory axis in promoting AML cell growth, we screened for synergistic interactions between RNF5 and epigenetic modulators. We assessed the effect of 261 epigenetic inhibitors at two concentrations (Supplemental Table 3) on the growth of AML cells that stably express inducible shRNF5 (Fig. 7A,B). Of the epigenetic inhibitors, 49 reduced viability of shRNF5-expressing cells compared with that of control AML cells (Fig. 7B). Among the inhibitors were several hypomethylation agents, including several histone methyltransferases (such as G9a), histone demethylases (such as Jumonji histone demethylases) and HDAC inhibitors (for example, TMP269, pimelic diphenylamide 106, and *N*-acetyldinaline [CI-994]). Because RBBP4 is a key component of the HDAC complex, and given that RNF5 KD induces transcriptional changes that resemble those seen upon HDAC1 inhibition (Fig. 5D,E), we further assessed possible synergy between RNF5 inhibition and HDAC inhibitors. CI-994, which is in clinical trials for several cancers (https://www.drugbank.ca/drugs/DB12291), was thus selected for additional validation. Indeed, U937 and HL-60 cells that were subjected to RNF5-KD exhibited a lower IC_50_ for cell viability by CI-994, compared with control cells (Fig. 7C and Extended Data Fig. 7A). These observations suggested that RNF5-KD sensitizes AML cells to HDAC inhibition.

**Fig. 7:**
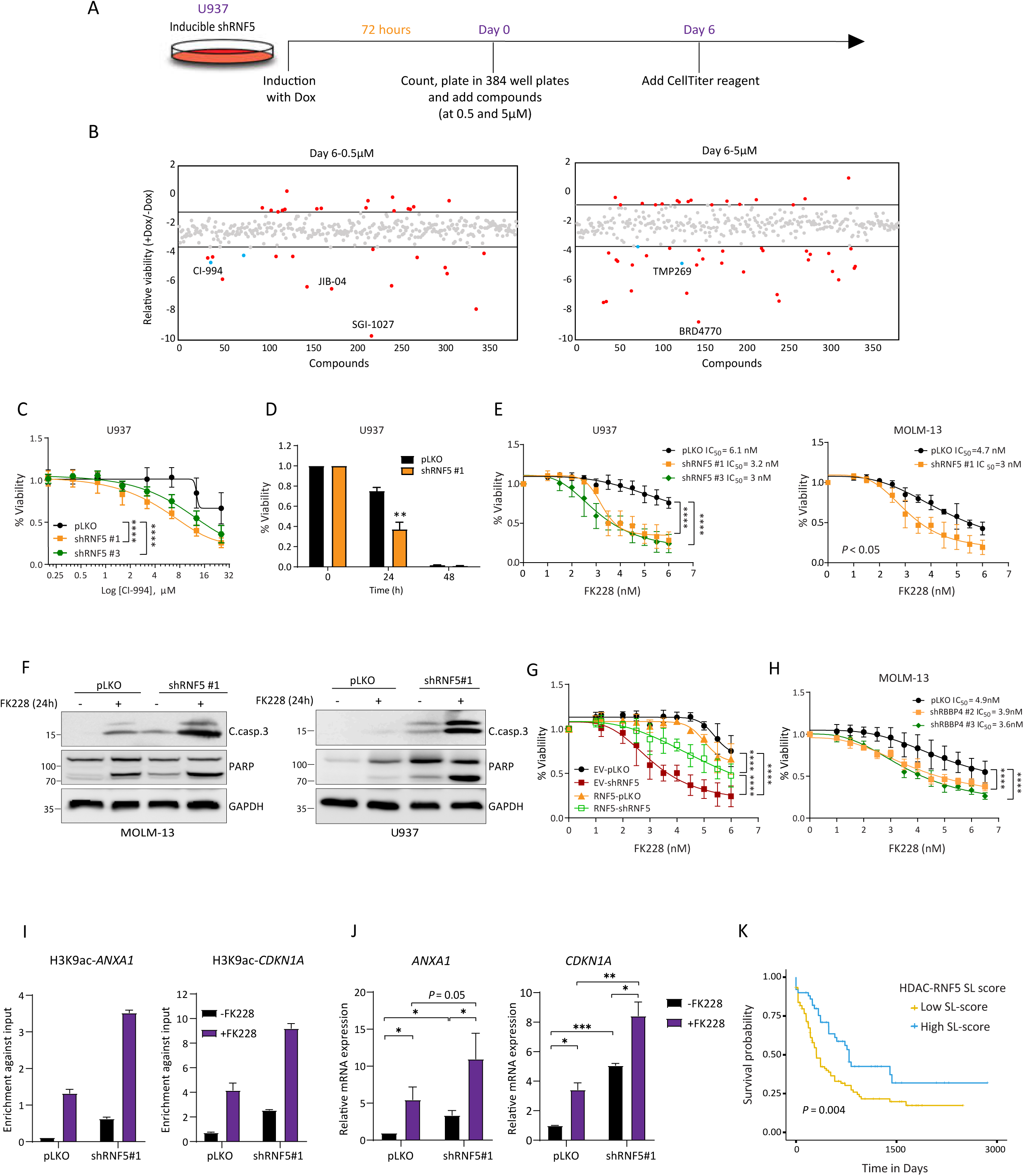
RNF5 inhibition sensitizes AML cells to HDAC inhibitors. **(A)** Experimental design of the epigenetic screen. U937 cells expressing inducible shRNF5 were treated with doxycycline (Dox) for 72 h, incubated with library compounds at two concentrations for 6 days, and analyzed for viability using the luminescence assay. (**B**) Log_2_-transformed ratios of the relative viability of doxycycline-induced (+Dox) versus uninduced (-Dox) U937 cells treated with compounds (0.5 μM or 5 μM) for 6 days. Grey dots represent all compounds tested, red dots represent candidate compounds that affected the viability of RNF5-KD more than of uninduced cells, and blue dots represent select HDAC inhibitors as potential candidates. **(C)** Viability of U937 cells expressing pLKO or shRNF5 after treatment for 24 h with the indicated concentrations of CI-994. **(D)** Viability of U937 cells after treatment for 24 h or 48 h with 3.5 nM FK228. (**E**) Viability of U937 cells or MOLM-13 cells expressing the indicated constructs after treatment for 24 h with the indicated FK228 concentrations. (**F**) Detection of apoptotic markers by Western blot analysis of MOLM-13 and U937 cells expressing pLKO or shRNF5#1. Cells were incubated with or without FK228 (4 nM 24 h) before lysis. **(G)** Viability of U937 cells expressing pLKO or shRNF5 and co-infected with control or RNF5 overexpression vectors. Cells were treated for 24 h with the indicated FK228 concentrations before analysis. EV-pLKO, control cells; EV-shRNF5, cells expressing empty vector and shRNF5; RNF5-pLKO, cells overexpressing RNF5 and empty pLKO vector; RNF5-shRNF5, cells overexpressing RNF5 and shRNF5. (**H**) Viability of MOLM-13 cells expressing the indicated constructs after treatment for 24 h with the indicated FK228 concentrations. (**I**) ChIP and qPCR indicating the enrichment of H3K9ac (normalized to input) at the indicated gene promoters in MOLM-13 cells expressing empty vector or shRNF5 constructs. Data are presented as the mean ± SD of two independent experiments. (**J**) RT-qPCR analysis of *ANXA1* and *CDKN1A* mRNA in MOLM-13 cells expressing empty vector (pLKO) or shRNF5 #1 and treated with 4nM FK228 for 15 h. (**K**) Kaplan-Meier plot showing the AML patients whose tumor exhibit lower HDAC and RNF5 transcription show better prognosis than the rest of patients (yellow lines) with hazard ratio of 0.42 (N^hi^=76, N^lo^=79). The patients were segregated into two groups based on median synthetic lethality (SL) score, which counts the occurrence of co-inactivation of SL partners in each sample. Logrank *P*-values are denoted in the figure, and the median survival differences are 488 days. Data are presented as the mean ± SD of three independent experiments. \**P* < 0.05, \*\**P* < 0.01, \*\*\**P* < 0.001, and \*\*\*\**P* < 0.0001 by two-tailed *t*-test (D and J) or by two-way ANOVA followed by Tukey’s multiple comparison test (C, E, G and H).

We noted that the HDAC inhibitor (romidepsin, also known as FK228) did not score positively in our screen. This was due to the relatively high concentrations tested, which were lethal to both RNF5-KD and control U937 cells. FK228 is approved by the Food and Drug Administration (FDA) for treatment of peripheral T-cell lymphoma ^34^ and was investigated in preclinical studies as a potential treatment for AML ^24, 35^. Therefore, we re-assessed FK228 at non-lethal concentrations (up to 6 nM for 24 h) in multiple AML cell lines. Notably, FK228 synergized with RNF5-KD in reducing cell viability (Fig. 7D,E and Extended Data Fig. 7B-D) and inducing apoptosis (Fig. 7F). We further confirmed the synergistic effect of HDACi in MOLM-13 cells in which *RNF5* was deleted using the CRISPR/Cas9 system (Extended Data Fig. 7E). The synergistic effect of RNF5-KD and FK228 on AML cell death was lost upon RNF5 re-expression (Fig. 7G), substantiating the role of RNF5 in sensitization of AML cells for HDAC inhibition.

Of the AML cell lines tested, we determined that the AML cell line MV-4-11 has a low amount of RNF5 (Extended Data Fig. 1B). MV-4-11 cells were the most sensitive to FK228 (Extended Data Fig. 7F), and RNF5-KD did not increase their sensitivity (Extended Data Fig. 7G). These observations further support the importance of RNF5 abundance in the response of AML cells to HDAC inhibitors. Moreover, RBBP4 KD sensitized AML cells to FK228 treatment (Fig. 7H and Extended Data Fig. 7H), consistent with our findings that RNF5 is a positive regulator of RBBP4. Notably, H3K9 acetylation on the promotors of RNF5 or RBBP4 regulated genes, *ANXA1* and *CDKN1A,* was elevated following FK228 treatment and further increased upon RNF5 KD (Fig. 7I). The latter is consistent with increased *ANXA1* and *CDKN1A* expression upon treatment with FK228 alone, or in combination with RNF5 KD (Fig. 7J and Extended Data Fig. 7I). To verify the relevance of the combination of RNF5 and HDAC in patients, we analyzed data from a bioinformatic pipeline that identifies clinically relevant synthetic lethal interactions ^36^. This analysis revealed a more favorable prognosis in patients with a concomitant downregulation of *HDAC* and *RNF5* (Fig. 7K). Collectively, these findings suggested that RNF5 signaling is central to the sensitivity of AML to HDAC inhibitors.

## CONCLUSIONS

High mortality of patients with AML predominantly results from failure to achieve complete remission following chemotherapy coupled with a high relapse rate. We identified the role that the ubiquitin ligase RNF5 plays in AML, and mechanisms underlying RNF5 contribution to this form of leukemia. Our studies highlighted a function for RNF5 beyond its established functions in ERAD and proteostasis ^6, 29^, demonstrating its impact on gene expression programs governing AML development and therapy response. RNF5 effect on gene expression is mediated via its interaction with and non-canonical ubiquitination of RBBP4, which enhanced the recruitment of RBBP4 to genes implicated in AML initiation and maintenance. The clinical relevance and the importance of RNF5 for AML was supported by our studies in genetic mouse models where RNF5 depletion inhibited the progression of AML and prolonged survival of mice. Furthermore, analysis of human AML samples revealed that high *RNF5* RNA and protein abundance, commonly seen in AML patient samples, correlated with poor prognosis.

RNF5-KD affected the expression of genes beyond those involved in proteostasis, in particular genes implicated in AML development and progression. We determined that RNF5 mediated these transcriptional changes, through a K29-toplogy ubiquitination of RBBP4. Limited reports on K29-linked protein ubiquitination highlights proteasome-independent functions, including changes in Wnt signaling ^37^, antiviral innate immune response ^38^, or protein aggregation in Parkinson disease ^39^. RNF5 was shown to affect cholesterol biosynthesis through induction of K29-linked polyubiquitination of the sterol regulatory element-binding protein 2 (SREBP2) chaperone SCAP ^40^. Notably, the importance of RNF5 control of RBBP4 for altered expression of genes seen in AML was substantiated by finding an inverse correlation between gene expression associated with RNF5- vs. RBBP4-expression. The implications of RNF5 effect on RBBP4 recruitment to transcriptional regulatory complexes were illustrated by the finding that RNF5 abundance affects the sensitivity of AML cells to HDAC inhibitors. Correspondingly, the transcriptional changes induced by RNF5 overlapped with those seen following HDAC1 inhibitor treatment. Further, synthetic lethal (ISLE) analysis identified a favorable prognosis in a cohort of AML patients with low *HDAC* and *RNF5* expression, substantiating the importance of RNF5 in the regulation of histone modification that control gene expression programs. Our results also indicated that *RNF5* expression may serve as a marker for stratification of patients for HDAC or proteasome inhibitors treatment. Lastly, our data suggested that pharmacological inhibitors that phenocopy RNF5-KD potentially represent a novel therapeutic modality for AML.

## METHODS

### Animal studies

All animal experiments were approved by the Sanford Burnham Prebys Medical Discovery Institute’s Institutional Animal Care and Use Committee (approval AUF 16-028). Animal care followed institutional guidelines. *Rnf5^−/−^* mice were generated on a C57BL/6 background as described ^41^. C57BL/6 WT mice were obtained by crossing *Rnf5^+/–^* mice. Female mice were maintained under controlled temperature (22.5°C) and illumination (12 h dark/light cycle) conditions and were used in experiments at 6 – 10 weeks of age.

The xenograft model was established using U937 expressing the p-GreenFire1 Lenti-Reporter Vector (pGFL). NOD/SCID (NOD.CB17-Prkdcscid/J) mice were obtained from SBP Animal Facility. Mice were irradiated (2.5 Gy), and U937-pGFL cells (2 × 10^4^ per mouse) expressing inducible shRNF5 or empty vector were then injected intravenously. Leukemia burden was serially assessed using noninvasive bioluminescence imaging by injecting mice intraperitoneally (i.p.) with 150 mg/kg D-Luciferin (PerkinElmer, 122799) in phosphate-buffered saline (PBS, pH 7.4), anesthetizing them with 2 – 3% isoflurane, and imaging them on an IVIS Spectrum (PerkinElmer). Upon disease onset (day 15), as measured by bioluminescent imaging, mice were fed with rodent chow containing 200 mg/kg doxycycline (Dox diet, Bio-Serv) to induce RNF5-KD. Mice were sacrificed upon signs of morbidity resulting from leukemic engraftment (> 10% weight loss, lethargy, and ruffled fur).

### Cell culture

Human HEK293T cells were obtained from the American Type Culture Collection (ATCC). U937 and K562 cells were kindly provided by Prof. Yuval Shaked; Kasumi-1 cells were from Prof. Tsila Zuckerman; and MV4-11, GRANTA, THP-1, and MEC-1 cells were from Dr. Netanel Horowitz. MOLM-13, U937, THP-1, Kasumi, Jurkat, and RPMI-8226 cells were cultured in RPMI medium; HL-60, MV-4-11, K-562, MEC-1, HAP-1, and KG-1α cells were cultured in IMDM; and GRANTA, A375 and HEK293T cells were cultured in DMEM. All media were supplemented with 10% fetal bovine serum (FBS), 1% L-glutamine, penicillin (83 U/mL), and streptomycin (83 µg/mL) (Gibco). Cells were regularly checked for mycoplasma contamination using a luminescence-based kit (Lonza).

### Primary AML cells

AML patient samples were obtained from Scripps MD Anderson, La Jolla, CA (IRB-approved protocol 13-6180) and written informed consent was obtained from each participant, and Rambam Health Campus Center, Haifa, Israel (IRB-approved protocol 0372-17). Fresh blood samples were obtained by peripheral blood draw, PICC line, or central catheter. Filgrastim-mobilized peripheral blood cells were collected from healthy donors and cryopreserved with DMSO. PBMCs were isolated by centrifugation through Ficoll-Paque^TM^ PLUS (17-1440-02, GE Healthcare). Residual red blood cells were removed using RBC Lysis Buffer for Human (Alfa Aesar, cat. # J62990) according to the manufacturer’s instructions. The final PBMC pellets were resuspended in Bambanker serum-free freezing medium (Wako Pure Chemical Industries, Ltd.) and stored under liquid N_2_.

### Antibodies and reagents

The RNF5 antibody was described previously ^7, 41^. Other antibodies were obtained as follows: rabbit anti-cleaved caspase 3 (#9661, Cell Signaling Technology), rabbit anti-PARP (#9532, Cell Signaling Technology), mouse anti-RBBP4 (NBP1-41201, Novus Biologicals), mouse anti-glyceraldehyde 3-phosphate dehydrogenase (GAPDH; ab8245, Abcam), mouse anti-Tubulin (T9026, Sigma), mouse anti-Flag (F1804, Sigma), mouse anti-Myc-Tag (#2276, Cell Signaling Technology), mouse anti-HA (901501, Biolegend), rabbit anti-HDAC1 (#2062, Cell Signaling Technology), rabbit anti-HDAC2 (57156, Cell Signaling Technology), rabbit anti-Ezh2 (5246, Cell Signaling Technology), mouse anti-HSP90 (sc-13119, Santa Cruz Biotechnology), rabbit anti-p27, (#3688, Cell Signaling Technology), rabbit anti-p21 (#2947, Cell Signaling Technology), mouse anti-Ubiquitin (#3939, Cell Signaling Technology), rabbit anti-K63-linkage Specific Polyubiquitin (#5621, Cell Signaling Technology), rabbit anti-Actin (#4970, Cell Signaling Technology), rabbit anti-Histone H3 (#9717, Cell Signaling Technology), mouse anti-Caspase 3 (sc-56053, Santa Cruz Biotechnology), and mouse anti-Calregulin (sc-166837, Santa Cruz Biotechnology). Romidepsin, and *N*-acetyldinaline were purchased from Cayman Chemicals. Thapsigargin and tunicamycin were purchased from Sigma-Aldrich. MG132 was obtained from Selleckchem. Puromycin was purchased from Merck. Annexin V-FITC and propidium iodide were from BioLegend.

### Plasmids and constructs

Plasmids expressing Flag-RNF5-WT, Flag-RNF5-RM, and Flag-RNF5-ΔCT were described previously ^5, 7^. To generate doxycycline-inducible RNF5-WT, RNF5-RM, and RNF5-ΔCT overexpression vectors, coding sequences were amplified from pCDNA3.1-RNF5-WT, pCDNA3.1-RNF5-RM, and pCDNA3.1-RNF5-ΔCT, respectively. The amplified PCR product was inserted into EcoRI-linearized pLVX TetOne-puro plasmid (Clontech) using the NEBuilder HiFi Assembly kit (New England BioLabs). RBBP4 expression vector was obtained from Addgene (Plasmid #20715).

### Gene silencing

Lentiviral pLKO.1 vectors expressing RNF5 or RBBP4-specific shRNAs were obtained from the La Jolla Institute for Allergy and Immunology RNAi Center (La Jolla, CA, USA). Sequences were: shRNF5 #1 (TRCN0000004785) GAGTGTCCAGTATGTAAAGCT, shRNF5 #2 (TRCN0000004788) CGGCAAGAGTGTCCAGTATGT, shRNF5 #3 GAGGATGGATTGAGAGAAT, inducible shRNF5 has same sequence as shRNF5#1. shRBBP4#1 (TRCN0000286103) GCCTTTCTTTCAATCCTTATA, shRBBP4#2 (TRCN0000293556) TGGTCATACTGCCAAGATATC, shRBBP4#3 (TRCN0000293554) ATGCGTCACACTACGACAGTG.

### Transfections and infections

Transient transfections were carried out using CalFectin (SignaGen) according to the manufacturer’s recommendations. Lentiviral particles were prepared using standard protocols. In brief, HEK293T cells were transfected with the relevant vectors and the second-generation packaging plasmids ΔR8.2 and Vsv-G (Addgene). Virus-containing supernatants were collected 48 h later and added with Polybrene to AML cells pre-seeded at ∼5 × 10^5^/well in 24-well plates (Sigma-Aldrich). After 8 h, cells were transferred to 10-cm culture dishes for an additional 24 h prior to experiments.

### Western blotting

Cells were washed twice with cold PBS and lysed by addition of Tris-buffered saline (TBS)-lysis buffer (TBS [50 mM Tris-HCl pH 7.5, 150 mM NaCl], 0.5% Nonidet P-40, 1× protease inhibitor cocktail [Merck], and 1× phosphatase inhibitor cocktail ^42^) followed by incubation on ice for 20 min. Cells from healthy control subjects and AML patients were lysed using hot lysis buffer [100 mM Tris-HCl pH 7.5, 5% sodium dodecyl sulfate (SDS)] followed by incubation 5 min at 95°C and sonication. Some samples were subjected to fractionation using a subcellular protein fractionation kit (Thermo Scientific Pierce), as indicated. Samples were resolved by SDS-PAGE and transferred to nitrocellulose membranes. Membranes were incubated for 1 h at room temperature with blocking solution (0.1% Tween 20/5% non-fat milk in TBS) and then overnight at 4°C with the primary antibodies. Membranes were washed with TBS and incubated 1 h at room temperature with appropriate secondary antibodies (Jackson ImmunoResearch). Finally, proteins were visualized using a chemiluminescence method (Image-Quant LAS400, GE Healthcare).

### Immunoprecipitation

Cells were lysed in TBS-lysis buffer as described above, centrifuged for 10 min at 17,000*g*, and incubated overnight at 4°C with appropriate antibodies. Protein A/G agarose beads (Santa Cruz Biotechnology) were then added for 2 h at 4°C. Beads were pelleted by centrifugation, washed five times with TBS-lysis buffer, and boiled in Laemmli buffer to elute proteins. Finally, proteins were resolved by SDS-PAGE and subjected to Western blotting as described above.

### LC-MS/MS

MOLM-13 cells were infected with doxycycline-inducible Flag-tagged RNF5-encoding or empty plasmids and expression was induced by addition of doxycycline (1 µg/mL) for 48 h. The proteasome inhibitor MG132 (10 μM) was added for 4 h prior to harvest. Cells were lysed in TBS-lysis buffer, and lysates were incubated with anti-Flag-M2-agarose beads (Sigma-Aldrich) overnight at 4°C. Beads were washed with TBS-lysis buffer and proteins were eluted from beads by addition of 3×Flag peptides (150 μg/mL, Sigma) for 1 h at 4°C and then subjected to tryptic digestion followed by LC-MS/MS, as described ^43^.

Raw data were analyzed using MaxQuant (v1.5.5.0) ^44^ with default settings. Protein intensities were normalized using the median centering method. Fold changes were calculated by dividing protein intensity of the Flag immunoprecipitates from RNF5-overexpressing cells by the protein intensity of the Flag immunoprecipitates from control cells. Thresholds were set at two for fold-change and 0.05 for p value obtained using Student’s *t*-test. Proteins identified in all RNF5 immunoprecipitation replicates but in one or no control IP replicates were considered potential interactors with RNF5 if their corresponding fold-change was at least two. Data from the Crapome ^42^ repository were downloaded to filter potential contaminants. Cytoscape ^45^ was used to generate the RNF5 interaction network and the pathway enrichment analysis. Raw MS data were deposited in the MassIVE repository with the reference number MSV000083160.

### Immunofluorescence microscopy

Cells were placed on coverslips on glass slides using a StatSpin cytofuge and fixed with 4% paraformaldehyde for 20 min at room temperature. Slides were then rinsed three times in PBS, and the cells were permeabilized in 0.2% Triton X-100 in PBS for 5 min and blocked with 0.2% TX-100/10% FBS in PBS for 30 min. Primary antibodies were diluted in staining buffer (0.2% Triton X-100/2% FBS in PBS), added to the cells, and the slides were incubated overnight at 4°C in a humidified chamber. Slides were then washed three times in staining buffer, and secondary antibodies (Life Technologies) were diluted in staining buffer and added to slides for 1 h at room temperature in a humidified chamber shielded from light. Finally, slides were washed three times in staining buffer and mounted with Fluoroshield Mounting Medium containing 4′, 6-diamidino-2-phenylindole (DAPI; Sigma-Aldrich). Cells were analyzed using a fluorescence microscope (DMi8; Leica) with a 60× oil immersion objective. Images were processed using the 3D deconvolution tool from LASX software (Leica), and the same parameters were used to analyze all images.

### Cell viability assay

Cell viability and growth were assayed using CellTiter Glo kit (Promega) according to the manufacturer’s recommendations. Cell lines were plated in white 96-well clear-bottomed plates (Corning) at a density of 7 × 10^3^ cells/well and growth was monitored every 24 h using CellTiter Glo reagent. Viability was quantified by measuring luminescence intensity with an Infinite 2000 Pro reader (Tecan).

### Cell cycle analysis

Distribution of cells in each phase of the cell cycle was analyzed by staining with propidium iodide (Merck). Briefly, 1 × 10^6^ cells were washed twice with cold PBS and fixed in 70% ethanol in PBS at 4°C overnight. Cells were washed, pelleted by centrifugation, and treated with RNase A (100 μg/mL) and propidium iodide (40 μg/mL) at room temperature for 30 min. Cell cycle distribution was assessed by flow cytometry (BD LSRFortessa™, BD Biosciences), and data was analyzed using FlowJo software.

### Annexin V and propidium iodide staining

Cells were collected in FACS tubes, washed twice with ice-cold PBS, and resuspended in 100 µL PBS. Annexin V-APC (1.4 µg/mL) was added for 15 min at room temperature in the dark. Then, cells were washed in PBS and resuspended in 200 µL PBS and propidium iodide (50 µg/mL) was added. Samples were then analyzed by flow cytometry (BD LSRFortessa™, BD Biosciences).

### Colony-forming assays

For the soft agar assay, a base layer was formed by mixing 1.5% soft agar (low-melting point agarose, Bio-Rad) and culture medium at a 1:1 ratio and placing the mixture in 6-well plates. Cells were resuspended in medium containing 0.3% soft agar and added to the base layer at 1 × 10^4^ (MOLM-13) or 5 × 10^3^ (U937) cells/well. Agar was solidified by incubation at 4°C for 10 mins before incubation at 37°C. Plates were incubated at 37°C in a humidified atmosphere for 12–18 days. The cells were then fixed overnight with 4% paraformaldehyde, washed with PBS, and stained with 0.05% crystal violet (Merck) for 20 min at room temperature and washed again with PBS. Plates were photographed and colonies were counted on the captured images.

For the methylcellulose assay, WT or *Rnf5^−/–^* Lin^−^Sca1^+^c-Kit^+^ cells transformed with GFP-MLL-AF9 were resuspended in methylcellulose M3234 (Stem Cell Technologies) supplemented with 6 ng/mL IL-3, 10 ng/mL IL-6, and 20 ng/mL stem cell factor (PeproTech). Cells were then added to 35-mm dishes at 10^3^ cells/well and incubated for 6 – 7 days. Colonies were classified as compact and hypercellular (blast-like) or small and diffuse (differentiated). Virtually all colonies fell into one of these two categories.

### RT-qPCR analysis

RNA was extracted using a GenElute Mammalian Total RNA Purification Kit (Sigma) according to standard protocols. RNA concentration was measured using a NanoDrop spectrophotometer (ThermoFisher). cDNA was synthesized from aliquots of 1 μg of total RNA using a qScript cDNA Synthesis Kit (Quanta). Quantitative PCR was performed with SYBR Green I dye master mix (Roche) and a CFX connect Real-Time PCR System (Bio-Rad). Primer sequences are listed in Supplemental Table 4. Primer efficiency was measured in preliminary experiments, and amplification specificity was confirmed by dissociation curve analysis.

### Gene targeting using CRISPR/Cas9

RNF5 sgRNAs were cloned into the pKLV2-U6gRNA-(BbsI)-PGKpuro2ABFP-W lentiviral expression vector and transduced into Cas9-expressing cell lines. All gRNAs were cloned into the BbsI site of the gRNA expression vector as previously described ^46^. Briefly, HEK293T cells were co-transfected with pKLV2-U6gRNA-(BbsI)-PGKpuro2ABFP-W and ectopic packaging plasmids using CalFectin transfection reagent (SignaGen). Virus-containing supernatants were collected 48 h later. MOLM-13 cells were infected by addition of supernatants for 48 h. Cells were then selected with puromycin (0.5 µg/mL) for 48 h and viability was measured. The RNF5-targeting sgRNA sequences were: sgRNF5 #3 F-GCACCTGTACCCCGGCGGAA, R-TTCCGCCGGGGTACAGGTGC. sgRNF5 #4 F-GTTCCGCCGGGGTACAGGTG, R-CACCTGTACCCCGGCGGAAC.

### RNA-seq analysis

PolyA RNA was isolated using the NEBNext Poly(A) mRNA Magnetic Isolation Module, and bar-coded libraries were constructed using the NEBNext Ultra™ Directional RNA Library Prep Kit for Illumina (NEB, Ipswich, MA). Libraries were pooled and single end-sequenced (1×75) on the Illumina NextSeq 500 using the High output V2 kit (Illumina, San Diego, CA). Quality control was performed using Fastqc (v0.11.5, Andrews S. 2010), reads were trimmed for adapters, low quality 5′ bases, and minimum length of 20 using CUTADAPT (v1.1). The number of reads per sample and the number of aligned reads suggested that read quality and number were good and that the data were valid for analysis. High-quality data were then mapped to human reference genome (hg19) using STAR mapping algorithm (version 2.5.2a) ^47^. FeatureCounts implemented in Subread (v1.50) ^48^ was used to count the sequencing reads from mapped BAM files. Analyses of differentially expressed genes was subsequently performed using a negative binomial test method (edgeR) ^49^ implemented using SARTools R Package ^50^. A list of the differentially expressed genes was exported into Excel (Supplemental Table S1) and pathway analysis was performed by uploading the lists of differentially expressed genes to IPA (http://www.ingenuity.com) using the following criteria: |log2(fold change)| >0.4 and p value < 0.05, and compared to public data sets processed by IPA using Analysis Match function. The top matched experiments in LINCS ^51^ were selected by ranking the overall z-scores.

### Chromatin immunoprecipitation (ChIP)

ChIP assay was performed using ChIP Assay Kit (Millipore) following the manufacturer’s instructions. Cells were fixed in 1% formaldehyde in PBS for 10 minutes at 37°C. Briefly, 1×10^6^ cells were used for each reaction. Cells were fixed in 1% formaldehyde at 37°C for10 minutes, and nuclei were isolated with nuclear lysis buffer (Millipore) supplemented with protease inhibitor cocktail (Millipore). Chromatin DNA was sonicated and sheared to a length between 200 bp and 1000 bp. The sheared chromatin was immunoprecipitated at 4°C overnight with anti-H3K9ac (9649, Cell Signaling Technology), anti-H3K27ac (ab3594, Abcam), anti-H3K27me3 (9733, Cell Signaling Technology), anti-RBBP4 (NBP1-41201, Novus). IgG was used as a negative control and anti-RNA polII (Millipore) was used as a positive control antibody. Protein A/G bead-antibody/chromatin complexes were washed with low salt buffer, high salt buffer, LiCl buffer, and TE buffer to remove nonspecific binding. Protein/DNA complex was reverse cross-linked, and DNA was purified using NucleoSpin®. Levels of ChIP-purified DNA were determined with qPCR (see Supplemental Table 5 for primer sequences). The relative enrichments of the indicated DNA regions were calculated using the Percent Input Method according to the manufacturer’s instructions and are presented as % input.

### Small molecule epigenetic regulators screen

Aliquots of compounds (10 mM in DMSO) from a library of 261 epigenetic regulators were dispensed at final concentrations of 0.5 µM or 5 µM into the wells of a Greiner (Monroe, NC, Cat #781080) 384-well TC-treated black plate using a Labcyte Echo 555 acoustic pipetter (Labcyte, San Jose, CA). U937 cells expressing an inducible shRNF5 vector were induced with doxycycline for 72 h and dispensed into the prepared plates at a density of 5 × 10^2^/well in 50 µL RPMI-based culture medium (described above) using a Multidrop Combi (Thermo Fisher Scientific, Pittsburgh, PA). The plates were briefly centrifuged at 1000 rpm and incubated at 37°C with 5% CO_2_ for an additional 6 days using MicroClime Environmental lids (Labcyte, San Jose, CA). Plates were placed at room temperature for 30 min to equilibrate, 20 µL/well CellTiter-Glo Luminescent Cell Viability Assay reagent (Promega, Madison, WI) was added using a Multidrop Combi, and the plates were analyzed with an EnVision multimode plate reader (PerkinElmer, Waltham, MA).

For the analysis, the intensity of induced shRNF5-expressing cells was divided by the intensity of uninduced cells. Ratios were log_2_ transformed and the thresholds were calculated based on the distribution of the log_2_ ratios. The upper threshold was calculated as the Q3 + 1.5xQ, where Q3 is the third quartile and IQ is the interquartile. The lower threshold was calculated as the Q1 − 1.5xIQ, where Q1 is the first quartile. Ratios outside these thresholds were considered outliers from the global ratio distribution and thus were potential candidates for having a differential effect on RNF5-KD or control cells.

### MLL-AF9–mediated transformation of bone marrow cells and generation of MLL-AF9– leukemic mice

HEK293T cells were co-transfected with Murine Stem Cell Virus (MSCV)-based MLL-AF9 IRES-GFP ^22^ and ectopic packaging plasmids. Viral supernatants were collected 48 h later and added to Lin^−^Sca-1^+^c-Kit^+^ cells isolated from the bone marrow of WT or *Rnf5^−/–^* C57BL/6 mice. Transduced cells were maintained in DMEM supplemented with 15% FBS, 6 ng/mL IL-3, 10 ng/mL IL-6, and 20 ng/mL stem cell factor, and transformed cells were selected by sorting for GFP^+^ cells. To generate “primary AML mice,” GFP-MLL-AF9–transduced cells were resuspended in PBS at 1 × 10^6^ cells/200 µL and injected intravenously into sublethally irradiated (650 Rad) 6- to 8-week-old C57BL/6 female mice.

### Statistical analysis

Differences between two groups were assessed using two-tailed unpaired or paired *t*-tests or Wilcoxon rank-sum test, and differences between group means were evaluated using *t*-tests or ANOVA. Two-way ANOVA with Tukey’s multiple comparison test was used to evaluate experiments involving multiple groups. Survival was analyzed by the Kaplan–Meier method and evaluated with a log-rank test. All analyses were performed using Prism software (GraphPad, La Jolla, CA, USA). *P* < 0.05 was considered significant.

## ACKNOWLEDGEMENTS

We thank Drs. Yuval Shaked, Tsila Zuckerman, and Netanel Horowitz (Faculty of Medicine, Technion) for providing leukemic cell lines, and members of the Deshpande and Ronai labs for technical support and discussions. We thank SBP and Technion Core facilities for help along different phases of this study. We thank Nancy R. Gough (BioSerendipity, LLC) for editorial assistance. Z.A.R. gratefully acknowledges support from the National Cancer Institute grant (R35CA197465) and the Technion. A.K. was supported by a Faculty of Medicine fellowship at the Technion. Sanford Burnham Prebys Shared Resources are supported by the NCI Cancer Center Support Grant (P30 CA030199).

## DATA AVAILABILITY

MS data is available on MSV online (MSV000083160). RNAseq data is available at the Gene Expression Omnibus under accession code GSE155929. Source files for original data (qPCR – excel files; Westerns raw data in PDF format are available online.

## AUTHOR CONTRIBUTIONS

A.K. and Z.A.R. designed the experiments; A.K., A.D., Yo.F., I.L., Y.L., and I.L., performed the experiments; A.K., J.S.L., B.F., D.F., Y.F., T.Z., J.Y., E.R., A.J.D and Z.A.R. analyzed the data; I.P., and M.J., performed the epigenetic screen; D.F., R.A., N.H., Y.O., C.B., J.R.M., and K.V. provided access to patient samples and data; and A.K. and Z.A.R. wrote the manuscript.

## COMPETING INTERESTS STATEMENT

ZAR is co-founder and serves as scientific advisor to Pangea Therapeutics. All other authors declare no competing interests.

TABLES – This manuscript contains 5 supplemental tables

FIGURES – This manuscript contains 7 figures and 7 extended data figures

## EXTENEDED DATA FIGURE LEGENDS

**Extended Data Fig. 1:**
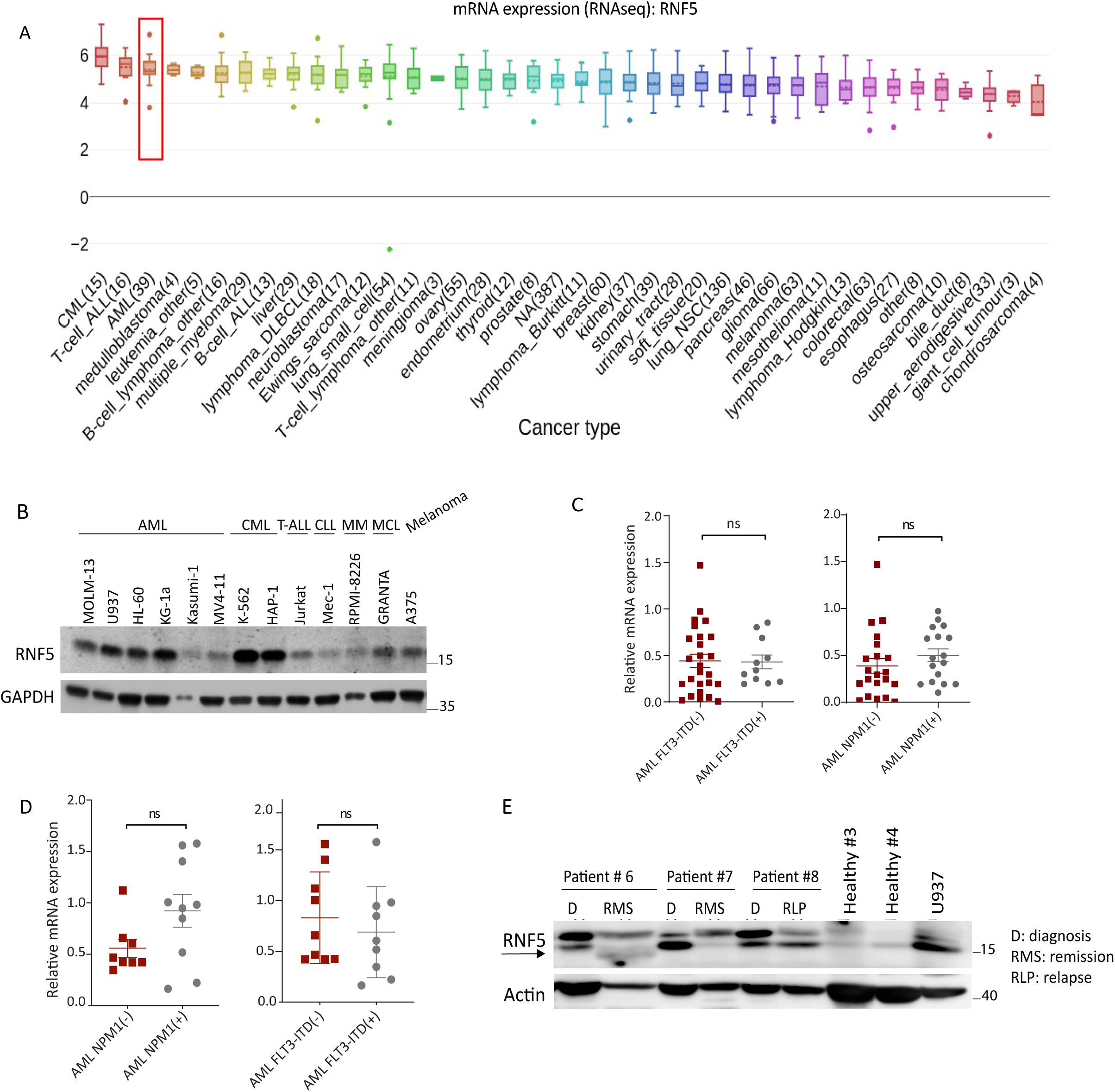
Level of RNF5 protein and transcripts inversely correlates with AML patient outcome. **(A)** *RNF5* expression data obtained from RNA-seq datasets in the Cancer Cell Line Encyclopedia. Transcript per million (TPM) gene expression data for protein-coding genes using RSEM. Log2 transformed, using a pseudo-count of 1w. The box plot is sorted and colored by the average distribution of *RN5* expression in a lineage. Lineages are composed of a number of cell lines from the same area or system of the body. The number next to the lineage name indicates how many cell lines are in the lineage. The highest average distribution is on the left and is colored red. The dashed line within a box is the mean. **(B)** Western blot analysis of RNF5 in lysates of a panel of cancer cell lines: AML, acute myeloid leukemia; CML, chronic myeloid leukemia; ALL, acute lymphoblastic leukemia; CLL, chronic lymphoblastic leukemia; MM, multiple myeloma; MCL, mantle cell lymphoma; and melanoma. **(C, D)** Relative *RNF5* expression in AML samples positive or negative for NPM1 or FLT3 mutations. Samples were obtained from Scripps Health Center (C) or Rambam Health Care Campus Center (D). **(E)** Western blot analysis of RNF5 abundance in PBMCs from healthy donors and AML patients in the Rambam Health Campus Center cohort. The position of RNF5 is indicated. The upper band is unspecific.

**Extended Data Fig. 2:**
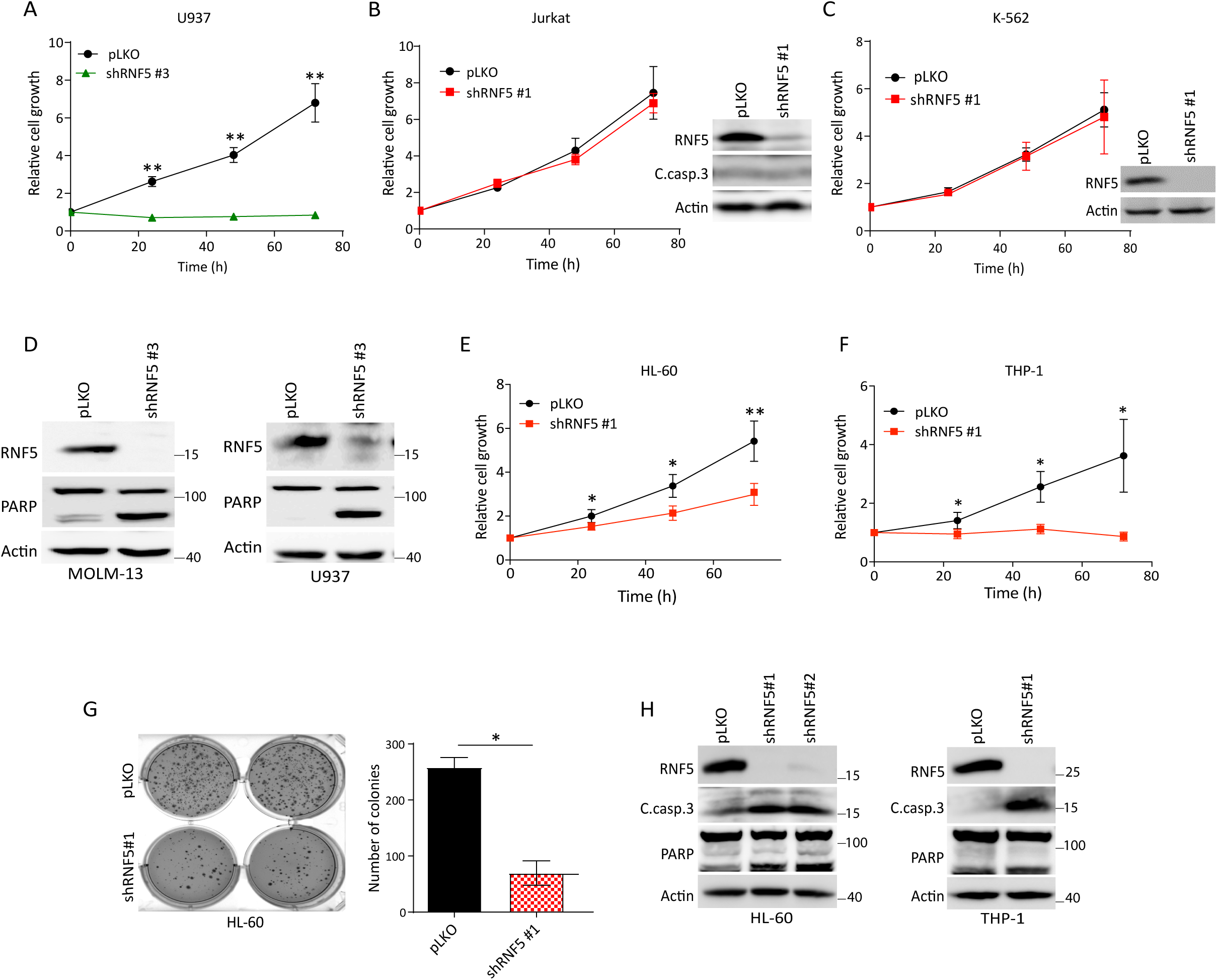
RNF5 is required for AML cell growth. **(A)** Luminescence growth assay of U937 cells expressing empty vector (pLKO) or shRNF5 #3. **(B)** Luminescence growth assay of Jurkat cells expressing pLKO or shRNF5 #1. Western blot shows knockdown efficiency. (C. casp.3 = cleaved caspase 3). **(C)** Luminescence growth assay of K562 cells expressing pLKO or shRNF5 #1. Western blot shows knockdown efficiency. **(D)** Western blot analysis of MOLM-13 and U937 cells 5 days after transduction with pLKO or the shRNF5 #3. Data are representative of three experiments. **(E, F)** Luminescence growth assays of HL-60 or THP-1 cells transduced with pLKO or shRNF5 #1. **(G)** Plate images (left) and quantification (right) of HL-60 colonies in soft agar. Colonies were assessed after 14 days in culture. **(H)** Western blot analysis of the indicated proteins in HL-60 and THP-1 cells 5 days after transduction with pLKO, shRNF5 #1, or shRNF5 #2. Data are representative of three experiments. Quantified data are presented as the mean ± SD and are representative of three independent experiments. \**P* < 0.05 and \*\**P* < 0.01 by two-tailed *t*-test.

**Extended Data Fig. 3:**
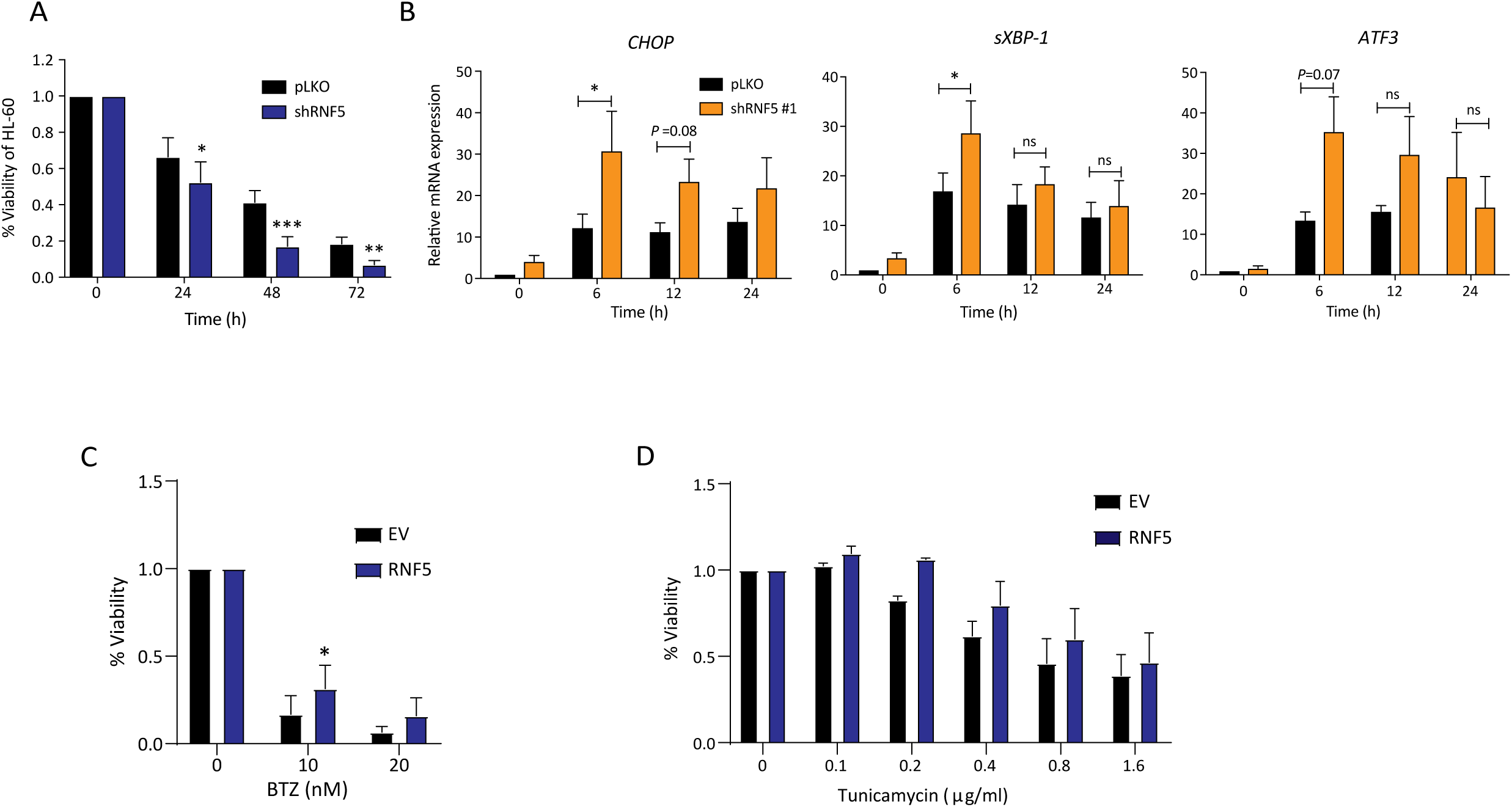
ER Stress synergizes with RNF5 inhibition in AML cells. (**A**) Luminescence growth assay of HL-60 cells expressing pLKO or shRNF5 after treatment with tunicamycin (2 µg/mL) for the indicated times. **(B)** RT-qPCR analysis of UPR-related genes in HL-60 cells treated with thapsigargin (1 µM) for the indicated times. **(C)** Luminescence growth assay of MOLM-13 cells induced to express Flag-tagged RNF5 for 2 days with doxycycline 1 µg/mL and then treated with the indicated concentrations of bortezomib (BTZ) for 24 h. **(D)** Luminescence growth assay of MV-4-11 cells induced to express Flag-tagged RNF5 for 2 days with doxycycline 1 µg/mL and then treated with the indicated concentrations of tunicamycin for 24 h. Data are presented from two independent experiments. \**P* < 0.05, \*\**P* < 0.01, and \*\*\**P* < 0.001 by two-tailed *t*-test. ns: not significant.

**Extended Data Fig. 4:**
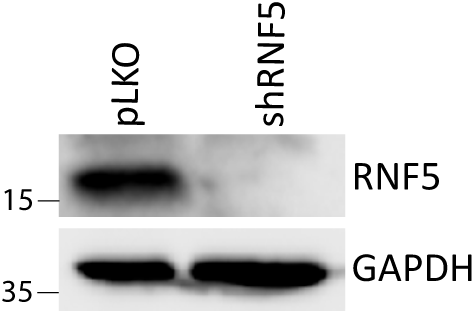
Validation of inducible RNF5 knockdown in U937 cells used for *in vivo* leukemia evaluation. U937-pGFL cells expressing pLKO or inducible shRNF5 were treated with Dox (1 µg/mL) for three days. Western blot analysis was performed to detect RNF5. GADPH served as a loading control.

**Extended Data Fig. 5:**
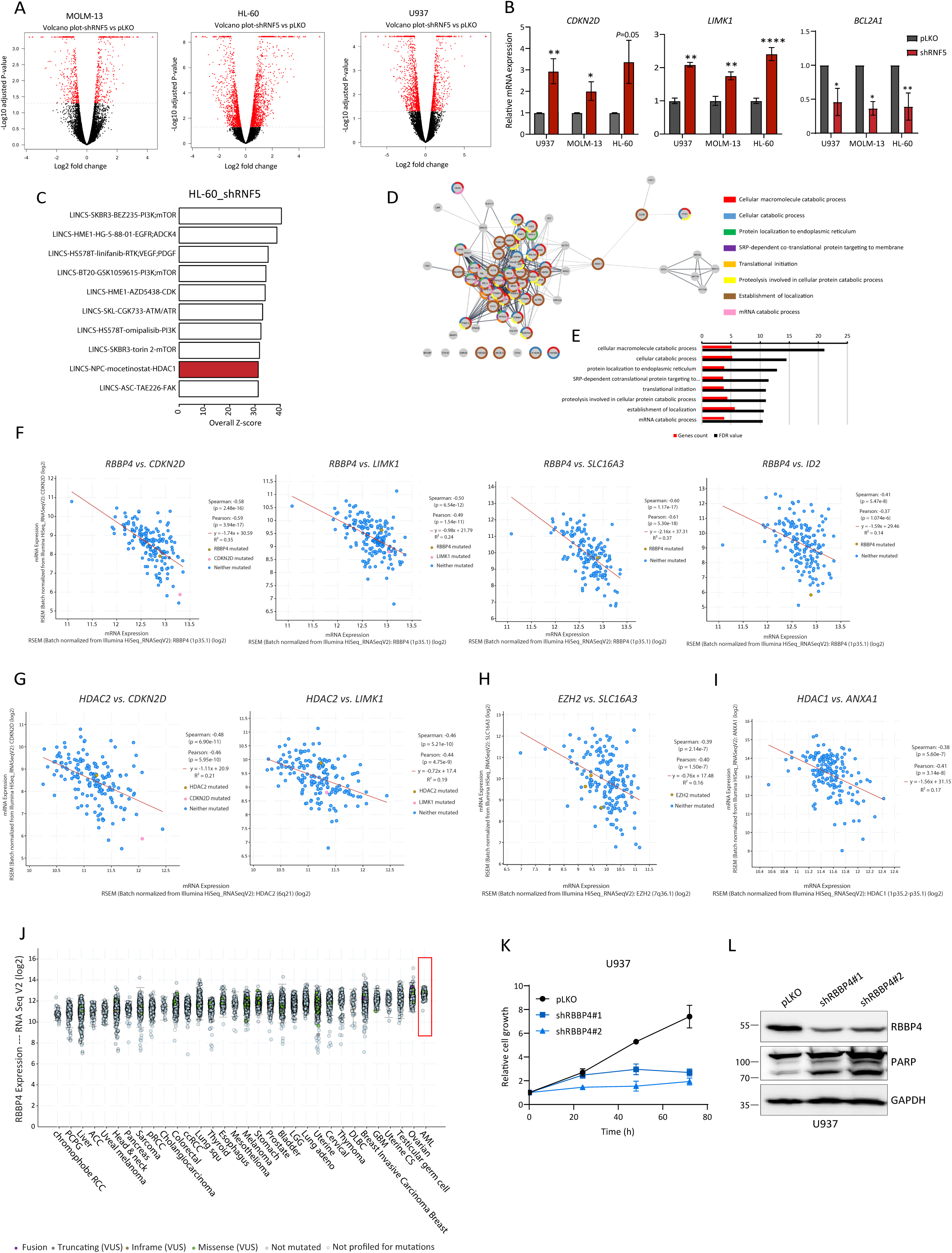
RNF5 affects gene transcription in AML cells. **(A)** Volcano plots highlight differentially expressed genes in WT compared with RNF5 KD AML cells. Each dot represents a gene. Genes with Benjamini-Hochberg (BH) corrected p value< 0.05 and Log2 transformed fold change >= 0.4 or <= −0.4 were selected as significantly differentially expressed genes and are highlighted in red. A list of differentially expressed genes is available in Supplementary Table 1. **(B)** qPCR validation of a select subset of genes identified as deregulated upon RNF5-KD by RNA-seq analysis. \**P* < 0.05, \*\**P* < 0.01, and \*\*\*\**P* < 0.0001 by two-tailed *t*-test. **(C)** Top ten drug screening results from LINCS database matched by shRNF5 in HL-60 cell line. Values are overall z-scores from IPA Analysis Match database. HDAC1 inhibitor results are shown in red. **(D)** RNF5 interaction network generated from the immunoprecipitation data and Cytoscape. The colors associated with the different pathways are displayed. **(E)** Pathway enrichment analysis displaying the genes count (log2 transformed) and the corresponding false discovery rate (-log10 transformed) for each pathway**. (F-I)** Co-expression of *RBBP4* (F), *HDAC2* (G), *EZH2* (H) or *HDAC1* (I) RNA and the indicated RNF5 target genes in AML analyzed in cBioPortal using data from TCGA. **(J)** Analysis of RBBP4 expression in different human cancers from the cBioPortal using data from TCGA. **(K)** Growth assay of U937 cells after transduction with empty vector (pLKO) or the indicated shRBBP4 constructs. Data are presented as the mean ± SD of two independent experiments. **(L)** Western blot analysis of the indicated proteins in U937 cells expressing empty vector (pLKO) or two different shRBBP4 constructs. Quantified data are presented as the mean ± SD and are representative of three independent experiments. \**P* < 0.05, \*\**P* < 0.01, \*\*\*\**P* < 0.0001 by two-tailed *t*-test.

**Extended Data Fig. 6:**
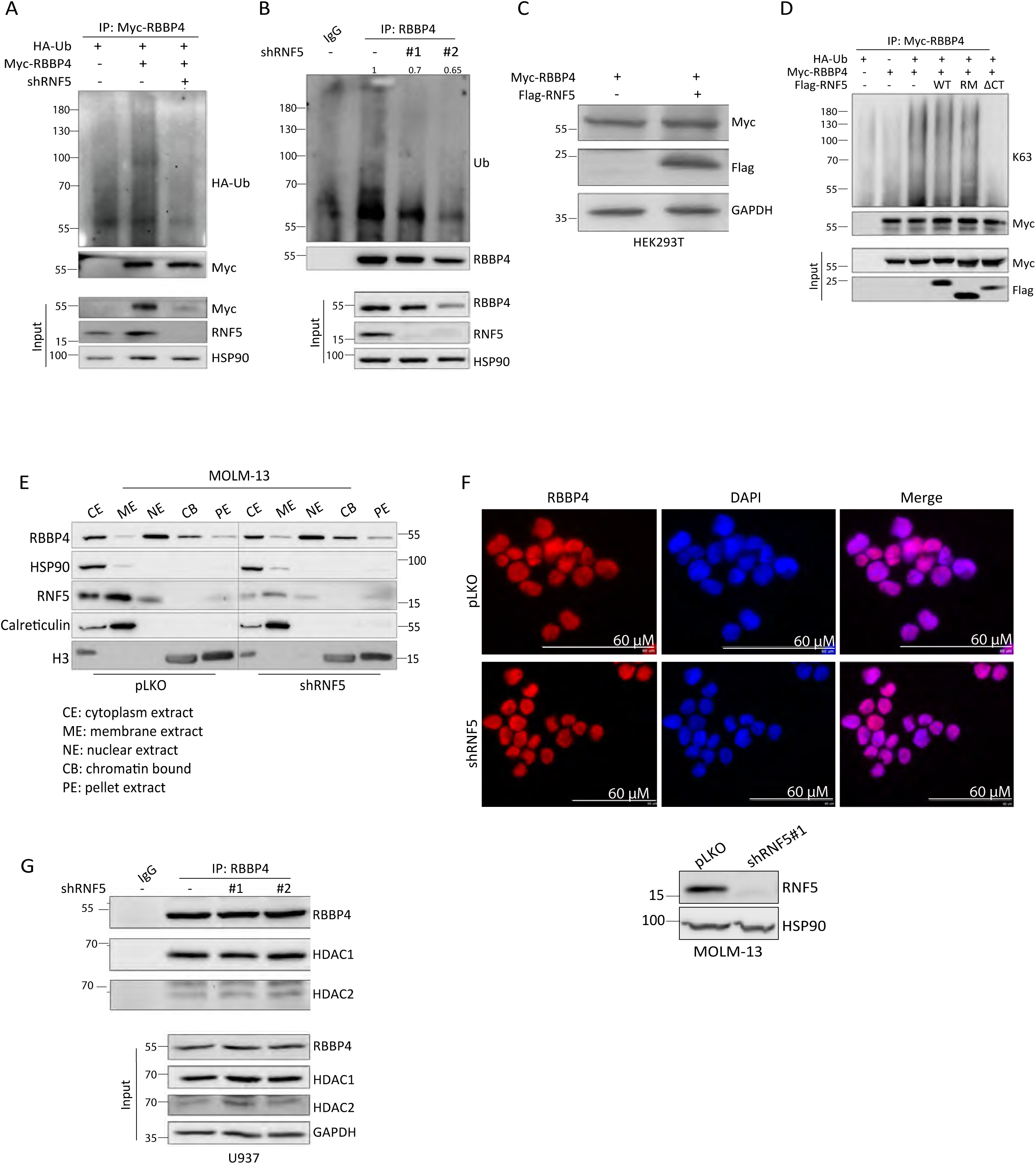
RNF5 interacts with- and ubiquitinates RBBP4. **(A)** Western blot analysis of anti-Myc immunoprecipitates and lysates of HEK293T cells co-expressing Myc-RBBP4, HA-Ub, and shRNF5. Cells were treated with MG132 (10 µm 4 h) before lysis. **(B)** Western blot analysis of anti-RBBP4 immunoprecipitates and lysates of MOLM-13 cells expressing the indicated shRNF5 constructs. Cells were treated with MG132 (10 µm 4 h) before lysis. Quantification of the ubiquitination smear relative to the amount of RBBP4 pull down is shown. **(C)** Western blot analysis of anti-Myc immunoprecipitates and lysates of HEK293T cells co-expressing Myc-RBBP4, HA-Ub, and the indicated Flag-tagged RNF5 constructs. Cells were treated with MG132 (10 µm 4 h) before lysis. **(D)** Western blot analysis of the indicated proteins in HEK293T cells transfected with Myc tagged RBBP4 (Myc-RBBP4) and Flag-tagged full-length RNF5 (Flag-RNF5). **(E)** Western blot analysis of RBBP4 and RNF5 in indicated cellular fractions in MOLM-13 cells expressing empty vector (pLKO) or shRNF5 #1. Histone H3, HSP90, and calreticulin serve as controls for chromatin, cytosol, and membrane fractions, respectively. **(F)** Immunofluorescence staining of RBBP4 (red) in control or shRNF5-expressing MOLM-13 cells. Nuclei were stained with DAPI (blue). Scale bar 60µM. Western blot shows RNF5-KD efficiency. **(G)** Immunoprecipitation and Western blot analysis of the interaction of RBBP4 with HDAC1, HDAC2, or EZH2 in U937 cells expressing empty vector or the indicated shRNF5 construct. Cells were treated with MG132 (10 µm 4 h) before lysis.

**Extended Data Fig. 7:**
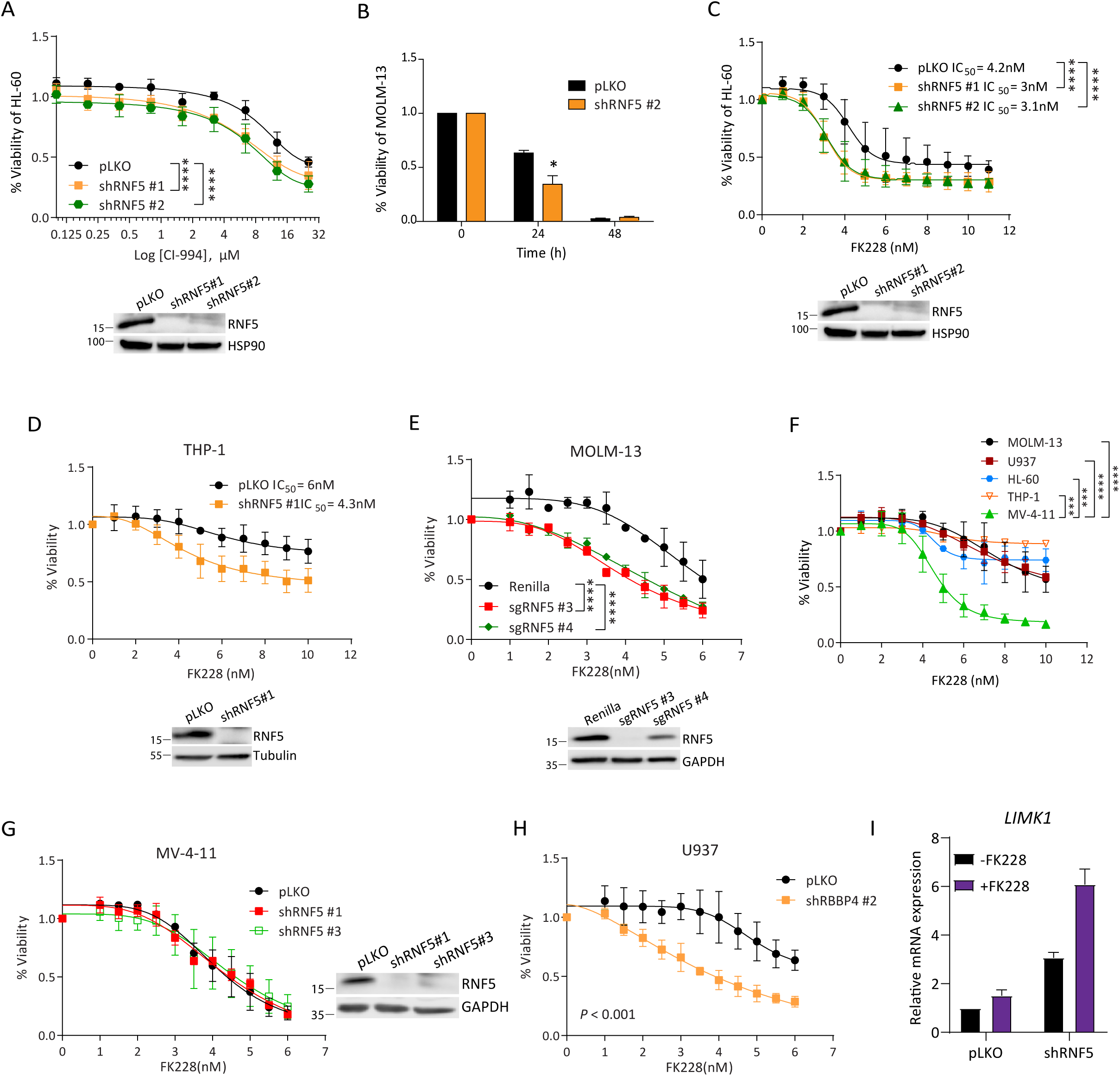
RNF5 KD sensitized AML cells to HDAC inhibition. **(A)** Viability of HL-60 cells expressing pLKO or two shRNF5 constructs after treatment for 24 h with the indicated concentrations of CI-994. Western blots show confirmation of RNF5-KD **(B)** Viability of MOLM-13 cells after treatment for 24 h with 3.5 nM FK228. **(C, D)** Viability of HL-60 (C) or THP-1(D) cells expressing pLKO or shRNF5 constructs after treatment for 24 h with the indicated concentrations of FK228. Western blots show confirmation of RNF5-KD. **(E)** Viability of MOLM-13 cells stably expressing Cas9 and transduced with control Renilla-targeting sgRNA or RNF5-targeting sgRNAs and treated for 24 h with indicated concentrations of FK228. Western blot shows reduction in RNF5. **(F)** Viability of MOLM-13, U937, MV-4-11, THP-1, and HL-60 cells after treatment for 24 h with the indicated concentrations of FK228. **(G)** Viability of MV-4-11 cells expressing pLKO or shRNF5 constructs and treated for 24 h with the indicated concentrations of FK228. Data are presented as the mean ± SD of two independent experiments. **(H)** Viability of U937 cells expressing pLKO or shRBBP4 and treated for 24 h with the indicated concentrations of FK228. **(I)** RT-qPCR analysis of *LIMK1* mRNA in MOLM-13 cells expressing empty vector (pLKO) or shRNF5 #1 and treated with 4nM FK228 for 15 h. Data are presented as the mean ± SD of two independent experiments. \**P* < 0.05, \*\*\**P* < 0.001, and \*\*\*\**P* < 0.0001 by two-tailed *t*-test (B) or by two-way ANOVA **(H)** followed by Tukey’s multiple comparison test (A, C, E and F).

## SUPPLEMENTAL TABLES

**Supplemental Table S1.** Genes commonly deregulated upon RNF5 knockdown in AML cell lines (HL-60, MOLM-13, and U937)

| Upregulated genes |  |  |  |  |  |  |  |
| --- | --- | --- | --- | --- | --- | --- | --- |
|  |  | Cell line |  |  |  |  |  |
|  |  | MOLM-13 |  | U937 |  | HL-60 |  |
| ID | Name | log2Fold Change | p.adj | log2Fold Change | p.adj | log2Fold Change | p.adj |
| ENSG00000102996.4 | MMP15 | 2.516 | 2.35E-04 | 1.201 | 4.25E-02 | 0.912 | 1.17E-02 |
| ENSG00000165178.8 | NCF1C | 1.729 | 3.79E-04 | 2.599 | 4.21E-04 | 1.901 | 3.08E-04 |
| ENSG00000196878.8 | LAMB3 | 1.676 | 1.08E-04 | 1.687 | 2.02E-02 | 2.877 | 1.09E-13 |
| ENSG00000078246.11 | TULP3 | 1.613 | 3.51E-05 | 1.073 | 6.31E-02 | 0.87 | 4.92E-03 |
| ENSG00000159388.5 | BTG2 | 1.473 | 1.28E-05 | 1.257 | 2.45E-02 | 1.842 | 2.35E-06 |
| ENSG00000185585.15 | OLFML2A | 1.414 | 1.26E-03 | 2.136 | 6.75E-05 | 3.116 | 1.37E-05 |
| ENSG00000182487.8 | NCF1B | 1.401 | 1.63E-03 | 3.021 | 1.18E-02 | 1.679 | 8.88E-04 |
| ENSG00000124762.9 | CDKN1A | 1.297 | 4.63E-06 | 2.513 | 3.88E-08 | 1.892 | 1.74E-04 |
| ENSG00000158517.9 | NCF1 | 1.286 | 2.74E-02 | 2.636 | 6.99E-05 | 1.74 | 9.42E-07 |
| ENSG00000135046.9 | ANXA1 | 1.128 | 1.61E-05 | 2.133 | 2.16E-08 | 0.406 | 3.69E-02 |
| ENSG00000129355.6 | CDKN2D | 1.103 | 4.47E-03 | 0.93 | 3.13E-02 | 1.849 | 1.86E-10 |
| ENSG00000009335.13 | UBE3C | 1.067 | 3.15E-06 | 1.619 | 1.25E-05 | 0.515 | 1.86E-02 |
| ENSG00000174007.7 | CEP19 | 1.055 | 8.12E-02 | 1.542 | 3.49E-02 | 1.24 | 7.78E-05 |
| ENSG00000107957.12 | SH3PXD2A | 0.955 | 2.82E-05 | 1.306 | 1.00E-02 | 0.478 | 1.29E-02 |
| ENSG00000151239.9 | TWF1 | 0.926 | 1.08E-04 | 1.248 | 1.22E-02 | 1.01 | 1.34E-04 |
| ENSG00000084112.10 | SSH1 | 0.838 | 2.01E-03 | 1.173 | 7.76E-03 | 0.561 | 3.69E-03 |
| ENSG00000115216.9 | NRBP1 | 0.823 | 2.87E-04 | 0.807 | 4.57E-02 | 1.108 | 4.35E-07 |
| ENSG00000139278.5 | GLIPR1 | 0.822 | 7.04E-03 | 3.01 | 3.25E-09 | 0.722 | 4.74E-04 |
| ENSG00000106683.10 | LIMK1 | 0.806 | 7.55E-03 | 1.069 | 3.77E-03 | 1.271 | 9.14E-09 |
| ENSG00000085491.11 | SLC25A24 | 0.791 | 3.57E-03 | 1.029 | 7.05E-03 | 1.35 | 8.75E-11 |
| ENSG00000167996.11 | FTH1 | 0.783 | 5.62E-02 | 1.343 | 2.18E-04 | 1.848 | 8.53E-15 |
| ENSG00000115738.5 | ID2 | 0.769 | 3.09E-02 | 1.495 | 3.07E-05 | 0.949 | 4.69E-05 |
| ENSG00000161638.6 | ITGA5 | 0.735 | 6.05E-03 | 1.002 | 8.49E-03 | 0.686 | 3.61E-04 |
| ENSG00000163297.12 | ANTXR2 | 0.721 | 3.68E-02 | 0.898 | 1.56E-02 | 1.014 | 1.93E-05 |
| ENSG00000100345.16 | MYH9 | 0.719 | 3.67E-03 | 0.812 | 5.04E-02 | 0.458 | 1.56E-02 |
| ENSG00000150093.14 | ITGB1 | 0.712 | 8.89E-03 | 1.189 | 9.87E-03 | 0.501 | 2.24E-02 |
| ENSG00000141526.10 | SLC16A3 | 0.696 | 3.86E-03 | 1.426 | 3.67E-05 | 0.535 | 1.66E-02 |
| ENSG00000141526.10 | SLC16A3 | 0.696 | 3.86E-03 | 1.426 | 3.67E-05 | 0.535 | 1.66E-02 |
| ENSG00000124145.5 | SDC4 | 0.688 | 9.72E-02 | 1.3 | 2.26E-03 | 1.356 | 7.10E-04 |
| ENSG00000127507.13 | EMR2 | 0.656 | 6.69E-03 | 1.933 | 5.13E-07 | 0.898 | 1.49E-07 |
| ENSG00000165410.10 | CFL2 | 0.61 | 7.53E-02 | 0.952 | 5.86E-02 | 1.18 | 6.67E-06 |
| ENSG00000069956.7 | MAPK6 | 0.605 | 1.59E-02 | 1.225 | 3.89E-04 | 0.544 | 6.08E-03 |
| ENSG00000172830.8 | SSH3 | 0.584 | 4.82E-02 | 1.869 | 1.50E-02 | 1.243 | 3.42E-04 |
| ENSG00000102007.6 | PLP2 | 0.576 | 4.33E-02 | 1.137 | 1.81E-03 | 0.587 | 5.57E-03 |
| ENSG00000117500.8 | TMED5 | 0.539 | 6.79E-02 | 0.87 | 2.80E-02 | 0.705 | 2.01E-04 |
| ENSG00000152104.7 | PTPN14 | 0.524 | 8.03E-02 | 1.818 | 6.64E-05 | 0.78 | 4.93E-03 |

| Downregulated genes |  |  |  |  |  |  |  |
| --- | --- | --- | --- | --- | --- | --- | --- |
|  |  | Cell line |  |  |  |  |  |
|  |  | MOLM-13 |  | U937 |  | HL-60 |  |
| ID | Name | log2Fold Change | p.adj | log2Fold Change | p.adj | log2Fold Change | p.adj |
| ENSG00000146433.8 | TMEM181 | -0.481 | 8.74E-02 | -0.934 | 1.47E-02 | -0.973 | 9.99E-05 |
| ENSG00000257093.2 | KIAA1147 | -0.625 | 3.68E-02 | -1.362 | 1.44E-04 | -1.105 | 3.96E-07 |
| ENSG00000124766.4 | SOX4 | -0.639 | 1.36E-02 | -1.063 | 3.92E-03 | -0.468 | 3.20E-02 |
| ENSG00000123562.12 | MORF4L2 | -0.655 | 8.10E-03 | -0.982 | 1.10E-02 | -0.431 | 3.91E-02 |
| ENSG00000109929.5 | SC5D | -0.675 | 1.83E-02 | -0.805 | 5.38E-02 | -0.609 | 2.35E-03 |
| ENSG00000112972.10 | HMGCS1 | -0.695 | 1.54E-02 | -2.068 | 3.59E-08 | -1.023 | 9.78E-08 |
| ENSG00000198689.5 | SLC9A6 | -0.801 | 6.61E-03 | -0.894 | 4.04E-02 | -0.564 | 2.02E-02 |
| ENSG00000180354.11 | MTURN | -0.863 | 3.27E-02 | -0.903 | 3.53E-02 | -1.716 | 1.82E-13 |
| ENSG00000160767.16 | FAM189B | -0.894 | 4.38E-03 | -1.461 | 8.54E-03 | -1.246 | 1.24E-04 |
| ENSG00000176390.10 | CRLF3 | -0.971 | 5.12E-03 | -1.213 | 5.31E-03 | -0.642 | 1.54E-02 |
| ENSG00000260196.1 | RP1-239B22.5 | -1.045 | 7.34E-03 | -1.159 | 1.10E-02 | -0.627 | 6.19E-03 |
| ENSG00000165389.6 | SPTSSA | -1.098 | 3.02E-06 | -1.062 | 1.17E-02 | -0.967 | 2.47E-06 |
| ENSG00000150459.8 | SAP18 | -1.239 | 4.98E-09 | -1.02 | 6.39E-03 | -0.917 | 2.72E-05 |
| ENSG00000171867.12 | PRNP | -1.255 | 2.93E-07 | -0.902 | 1.35E-02 | -1.254 | 1.72E-09 |
| ENSG00000182557.3 | SPNS3 | -1.296 | 5.85E-06 | -1.577 | 3.18E-03 | -1.085 | 5.98E-05 |
| ENSG00000070669.12 | ASNS | -1.325 | 1.45E-02 | -1.161 | 1.14E-02 | -0.644 | 4.77E-04 |
| ENSG00000121848.9 | RNF115 | -1.388 | 3.81E-09 | -1.644 | 1.51E-05 | -1.341 | 6.49E-11 |
| ENSG00000119414.7 | PPP6C | -1.392 | 8.66E-10 | -0.963 | 1.03E-02 | -1.164 | 1.70E-06 |
| ENSG00000113048.12 | MRPS27 | -1.398 | 2.07E-09 | -1.003 | 9.40E-03 | -1.64 | 5.78E-15 |
| ENSG00000140379.7 | BCL2A1 | -1.45 | 2.27E-02 | -1.124 | 2.71E-02 | -1.351 | 5.10E-03 |
| ENSG00000197223.7 | C1D | -1.515 | 1.27E-05 | -1.133 | 1.73E-02 | -0.545 | 3.05E-02 |
| ENSG00000134049.3 | IER3IP1 | -1.517 | 6.06E-08 | -0.848 | 3.83E-02 | -1.068 | 5.21E-09 |
| ENSG00000011028.9 | MRC2 | -1.551 | 6.19E-13 | -1.693 | 1.30E-03 | -1.135 | 2.53E-04 |
| ENSG00000204308.6 | RNF5 | -3.798 | 5.26E-45 | -3.647 | 1.51E-15 | -3.04 | 6.51E-14 |

**Supplemental Table S2.** List of RNF5 interacting proteins identified by LC-MS/MS.

| Protein name | Gene name | Number of Peptides | ratio IP RNF5 / IP Ctrl | p-value |
| --- | --- | --- | --- | --- |
| E3 ubiquitin-protein ligase RNF5 | RNF5 | 5 | Found only in RNF5 IP |  |
| BAG family molecular chaperone regulator 2 | BAG2 | 11 | Found only in RNF5 IP |  |
| Transitional endoplasmic reticulum ATPase | VCP | 20 | Found only in RNF5 IP |  |
| Large proline-rich protein BAG6 | BAG6 | 9 | Found only in RNF5 IP |  |
| Erlin-2 | ERLIN2 | 12 | Found only in RNF5 IP |  |
| FAS-associated factor 2 | FAF2 | 9 | Found only in RNF5 IP |  |
| 26S protease regulatory subunit 7 | PSMC2 | 4 | Found only in RNF5 IP |  |
| 26S protease regulatory subunit 6A | PSMC3 | 4 | Found only in RNF5 IP |  |
| 26S protease regulatory subunit 10B | PSMC6 | 4 | Found only in RNF5 IP |  |
| 26S proteasome non-ATPase regulatory subunit 11 | PSMD11 | 3 | Found only in RNF5 IP |  |
| Erlin-1 | ERLIN1 | 19 | 189.9 | 0.005 |
| Proteasome subunit beta type-6 | PSMB6 | 2 | 8.7 | 0.023 |
| Heat shock 70 kDa protein 1A | HSPA1A | 17 | 8.0 | 0.009 |
| Heat shock cognate 71 kDa protein | HSPA8 | 36 | 6.1 | 0.013 |
| Proteasome subunit alpha type-7 | PSMA7 | 6 | 4.6 | 0.038 |
| Proteasome subunit alpha type | PSMA6 | 8 | 4.3 | 0.004 |
| Proteasome subunit alpha type | PSMA1 | 4 | 4.1 | 0.014 |
| Proteasome subunit alpha type-2 | PSMA2 | 4 | 3.1 | 0.016 |
| Proteasome subunit alpha type-5 | PSMA5 | 3 | 2.8 | 0.039 |
| Proteasome subunit beta type-1 | PSMB1 | 4 | 2.8 | 0.006 |
| Heat shock protein HSP 90-beta | HSP90AB1 | 11 | 2.5 | 0.002 |
| Stress-70 protein, mitochondrial | HSPA9 | 7 | 2.2 | 0.014 |
| 78 kDa glucose-regulated protein | HSPA5 | 21 | 2.2 | 0.008 |
| Heat shock protein beta-1 | HSPB1 | 9 | 3.1 | 0.016 |
| T-complex protein 1 subunit theta | CCT8 | 7 | 2.7 | 0.024 |
| T-complex protein 1 subunit gamma | CCT3 | 6 | 2.1 | 0.002 |
| Actin-related protein 2 | ACTR2 | 4 | Found only in RNF5 IP |  |
| Tropomodulin-3 | TMOD3 | 4 | Found only in RNF5 IP |  |
| Transmembrane protein 33 | TMEM33 | 4 | Found only in RNF5 IP |  |
| Ubiquitin-associated domain-containing protein 2 | UBAC2 | 6 | Found only in RNF5 IP |  |
| Sequestosome-1 (p62) | SQSTM1 | 5 | 29.1 |  |
| Ubiquitin-40S ribosomal protein S27a | UBB | 11 | 12.2 | 0.008 |
| BRI3-binding protein | BRI3BP | 3 | 8.4 | 0.000 |
| ADP/ATP translocase 3 | SLC25A6 | 9 | 8.1 | 0.002 |
| ADP/ATP translocase 2 | SLC25A5 | 11 | 5.9 | 0.001 |
| Voltage-dependent anion-selective channel protein 2 | VDAC2 | 2 | 4.7 | 0.005 |
| Splicing factor 3B subunit 2 | SF3B2 | 3 | 4.4 |  |
| Gelsolin | GSN | 5 | 4.4 | 0.038 |
| Tubulin alpha-1B chain | TUBA1B | 17 | 4.0 | 0.003 |
| 60S acidic ribosomal protein P1 | RPLP1 | 4 | 3.7 | 0.049 |
| Eukaryotic translation initiation factor 3 subunit C | EIF3C | 3 | 3.7 | 0.049 |
| Annexin A1 | ANXA1 | 14 | 3.5 | 0.048 |
| 40S ribosomal protein S15a | RPS15A | 4 | 3.4 | 0.026 |
| Keratin, type I cytoskeletal 13 | KRT13 | 30 | 3.3 | 0.017 |
| 40S ribosomal protein S3 | RPS3 | 6 | 3.1 | 0.025 |
| Histone-binding protein RBBP4 | RBBP4 | 5 | 3.1 | 0.043 |
| Carboxypeptidase A4 | CPA4 | 3 | 2.9 | 0.047 |
| Actin, alpha skeletal muscle | ACTA1 | 19 | 2.8 | 0.035 |
| Keratin, type I cuticular Ha3-II | KRT33B | 58 | 2.8 | 0.042 |
| Keratin, type I cytoskeletal 19 | KRT19 | 14 | 2.5 | 0.007 |
| Pre-mRNA-splicing factor ATP-dependent RNA helicase DHX15 | DHX15 | 3 | 2.4 | 0.001 |
| Keratin, type II cuticular Hb4 | KRT84 | 56 | 2.4 |  |
| 60S ribosomal protein L30 | RPL30 | 3 | 2.4 | 0.010 |
| 60S ribosomal protein L4 | RPL4 | 17 | 2.3 | 0.022 |
| Protein S100-A8 | S100A8 | 4 | 2.3 | 0.038 |
| ATP synthase subunit alpha, mitochondrial | ATP5A1 | 7 | 2.2 | 0.020 |
| Serine/threonine-protein kinase 38 | STK38 | 17 | 2.2 | 0.035 |
| Elongation factor 1-delta | EEF1D | 6 | 2.2 | 0.015 |
| 60S ribosomal protein L18a | RPL18A | 8 | 2.1 | 0.040 |
| Tubulin beta chain | TUBB | 13 | 2.1 | 0.037 |
| 40S ribosomal protein S24 | RPS24 | 5 | 2.1 | 0.033 |
| Protein phosphatase 1B | PPM1B | 16 | 2.1 | 0.034 |
| 60S ribosomal protein L10 | RPL10 | 7 | 2.1 | 0.003 |
| 40S ribosomal protein S8 | RPS8 | 11 | 2.0 | 0.021 |

**Supplemental Table S3.**
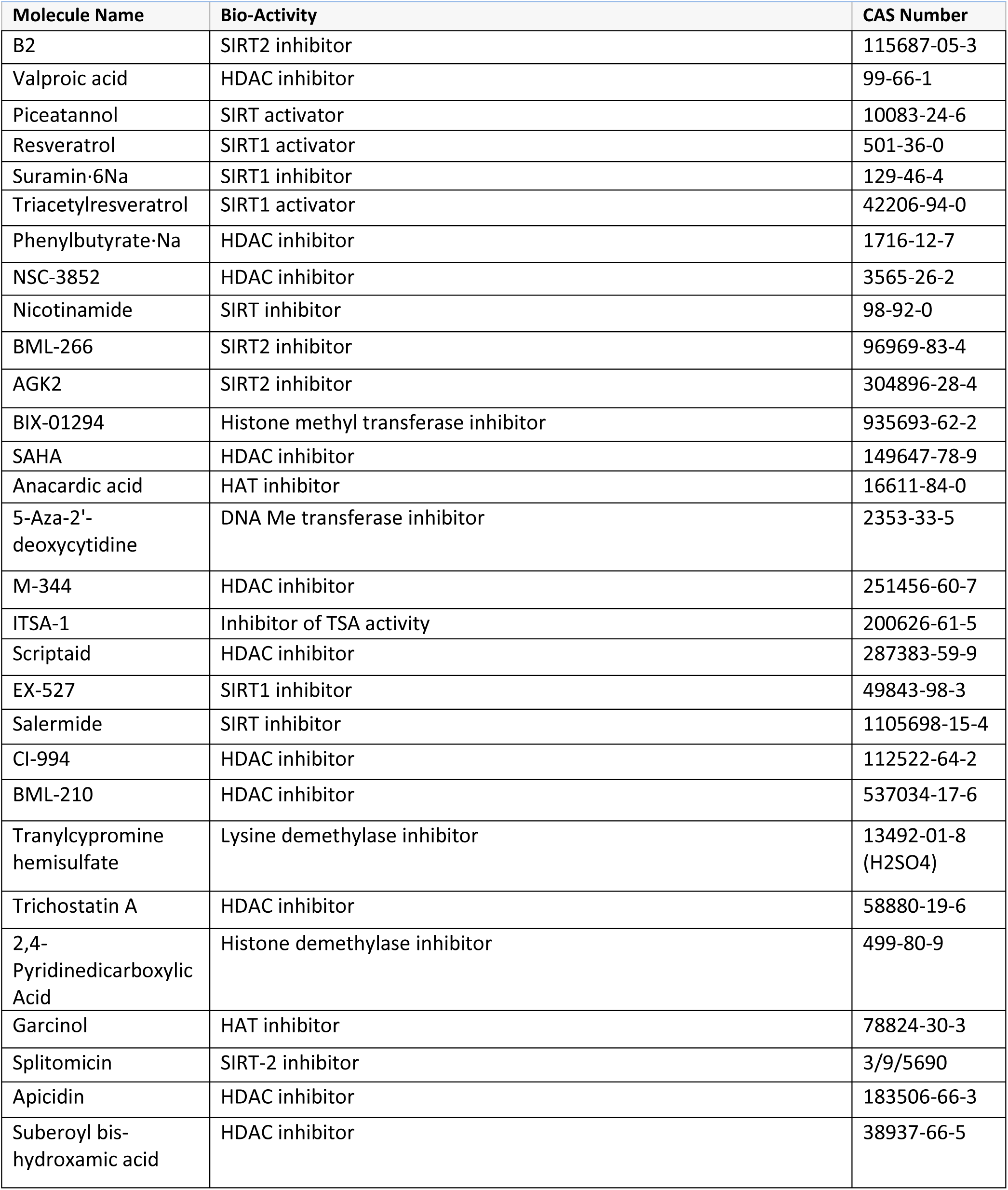
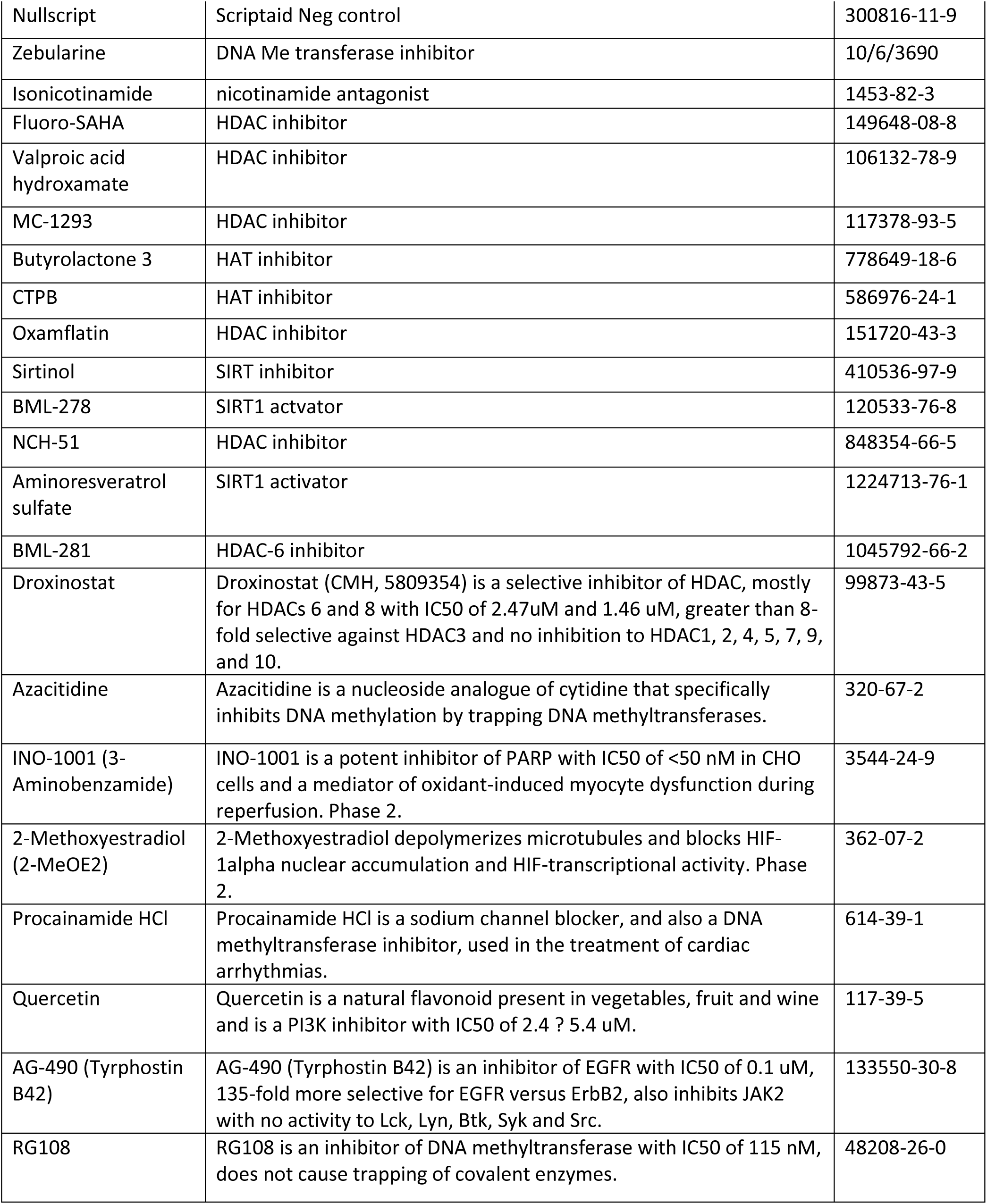
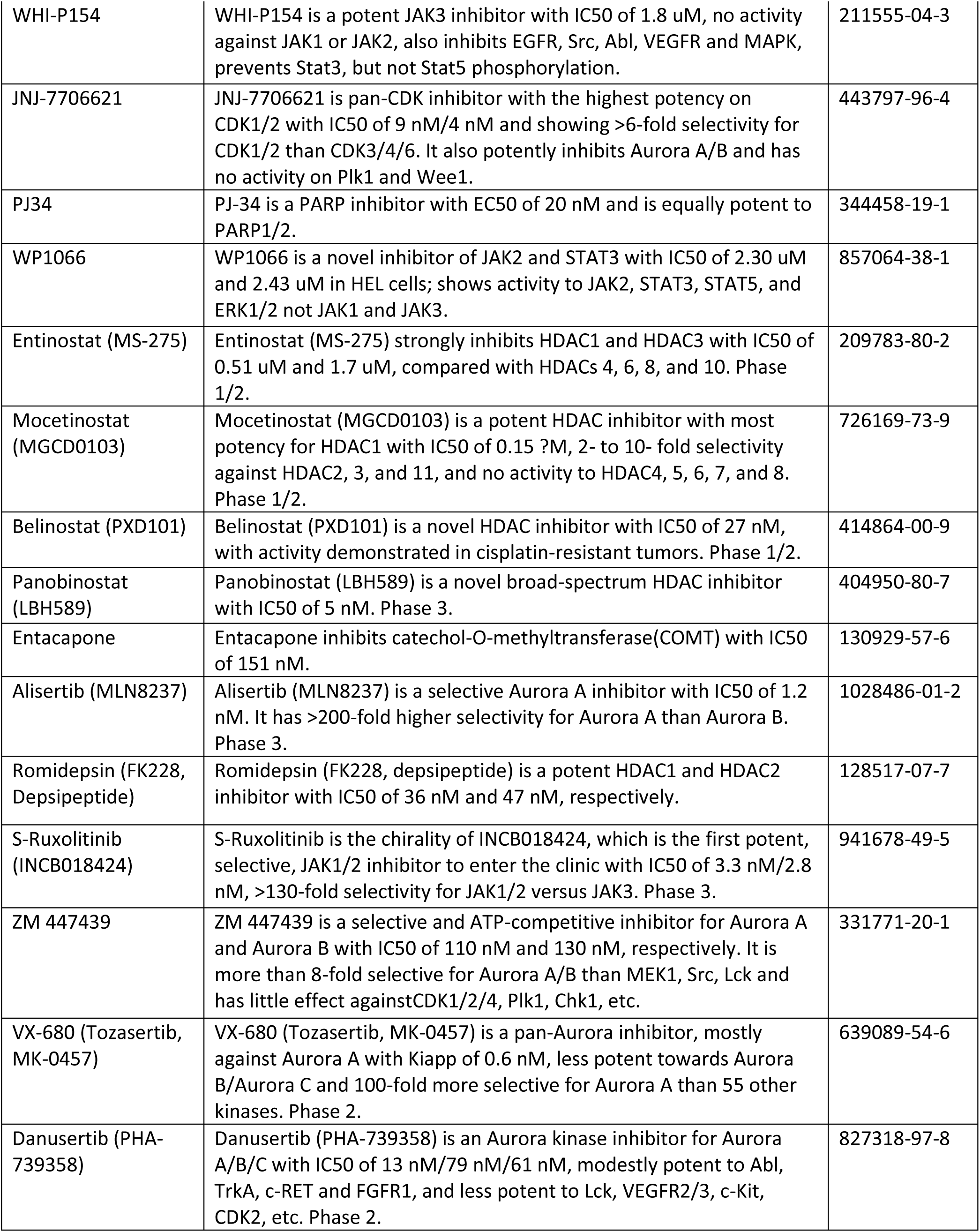
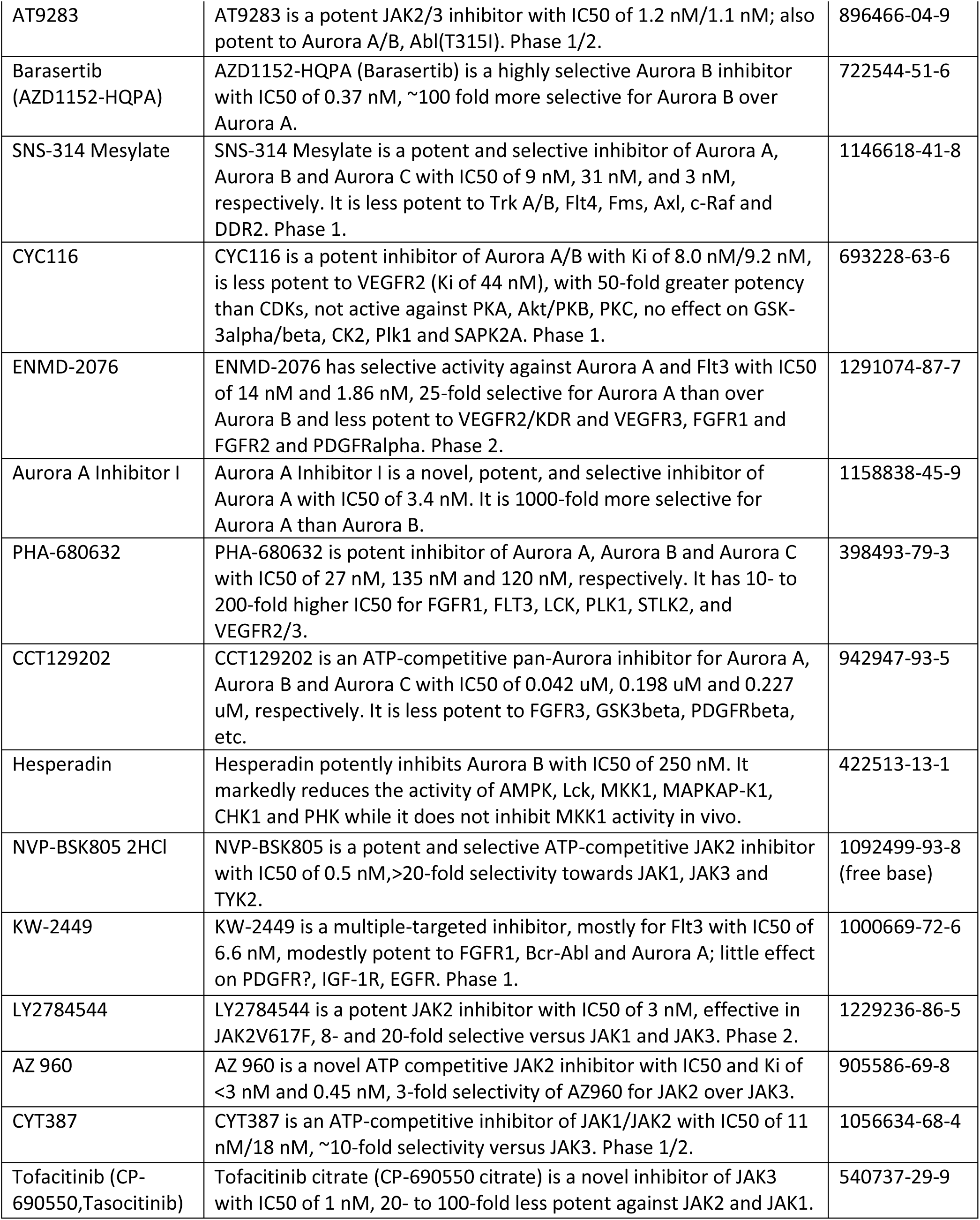
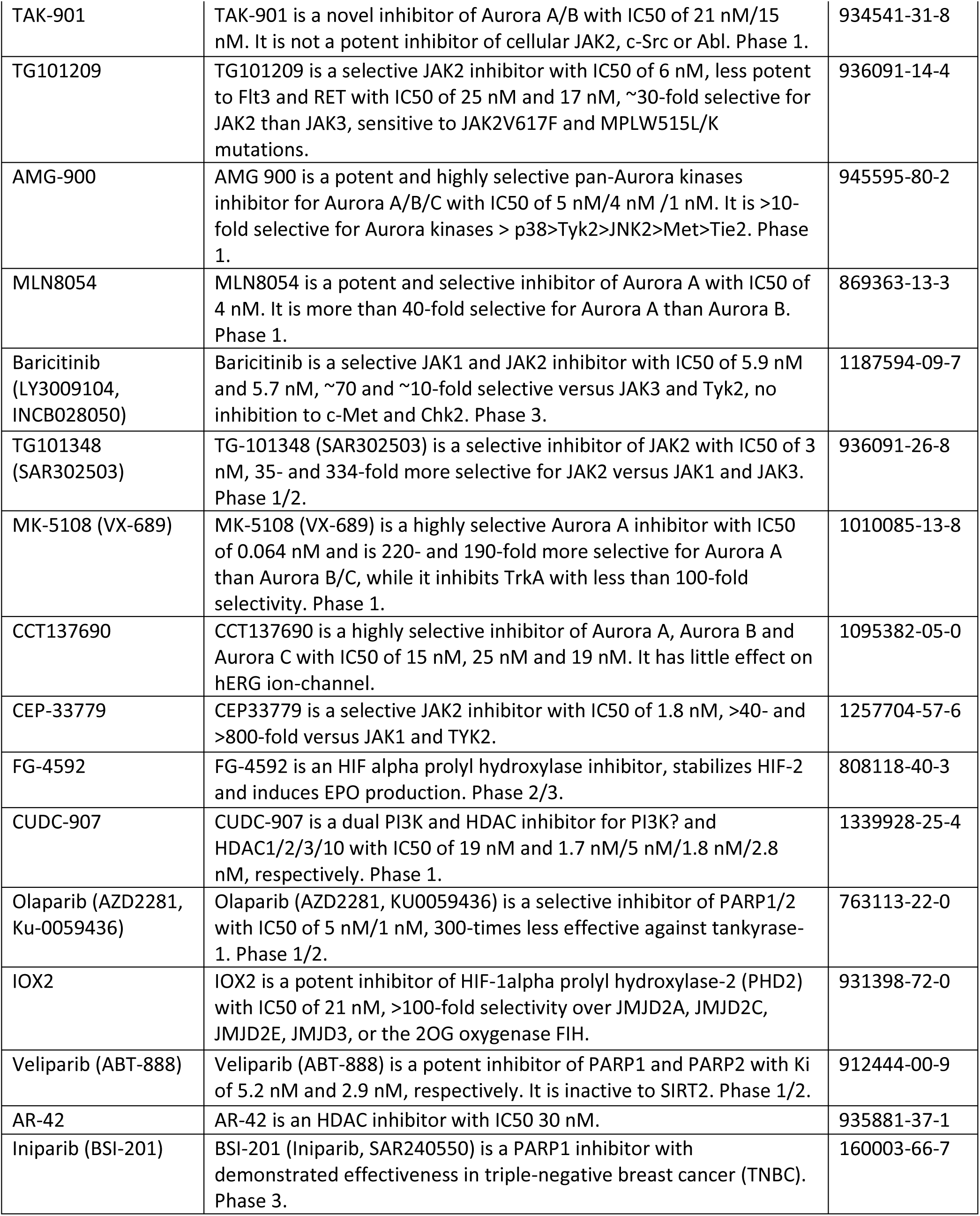
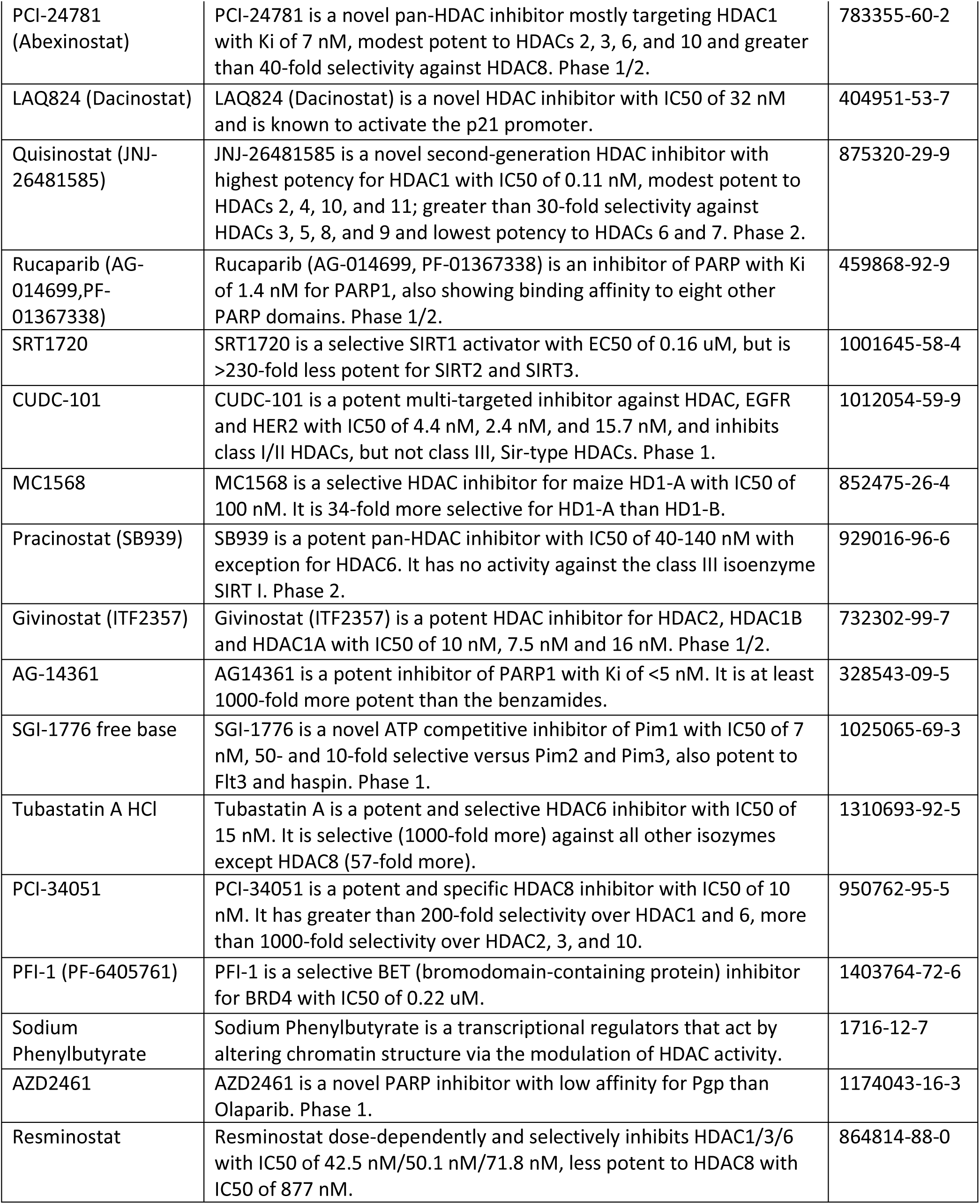
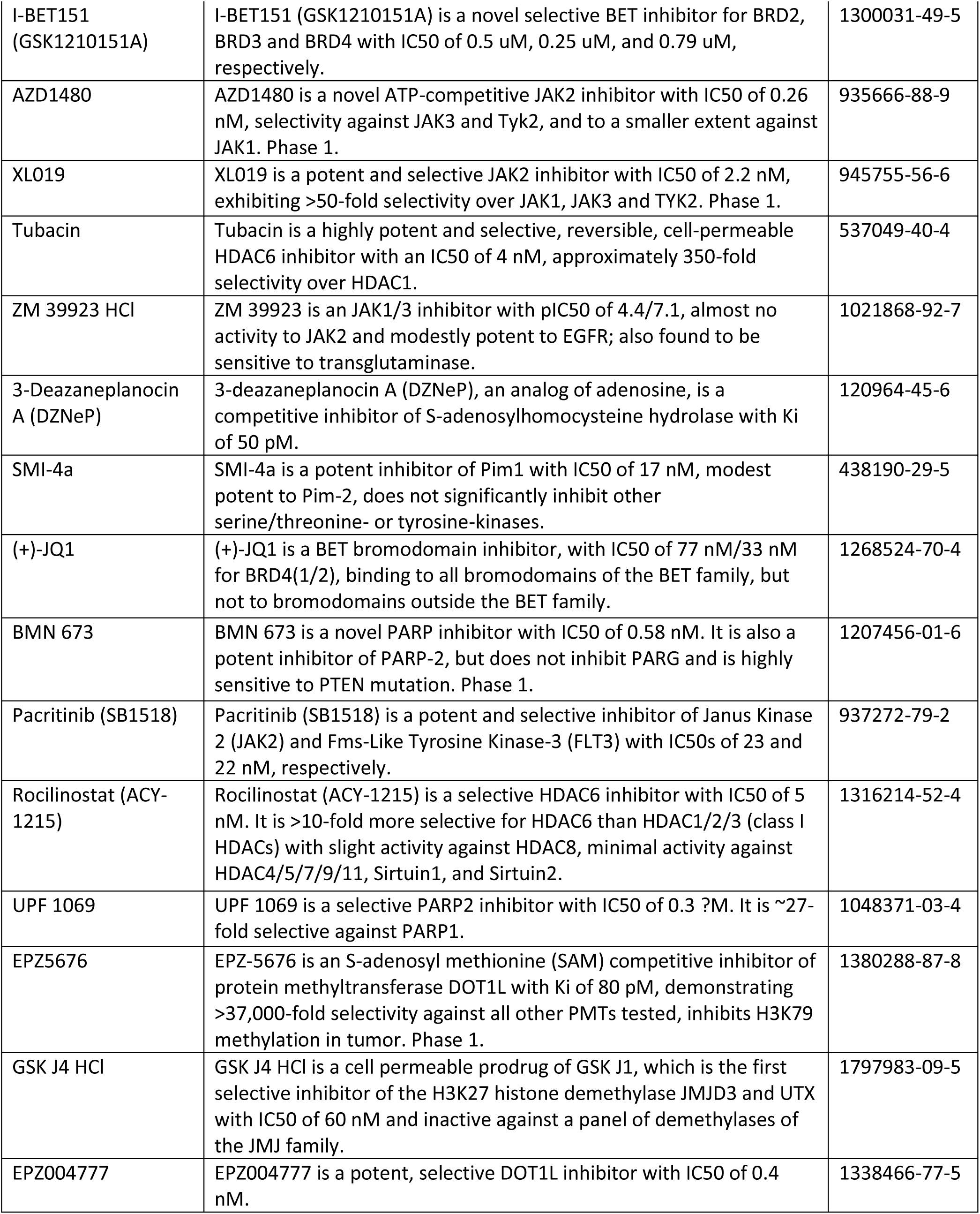
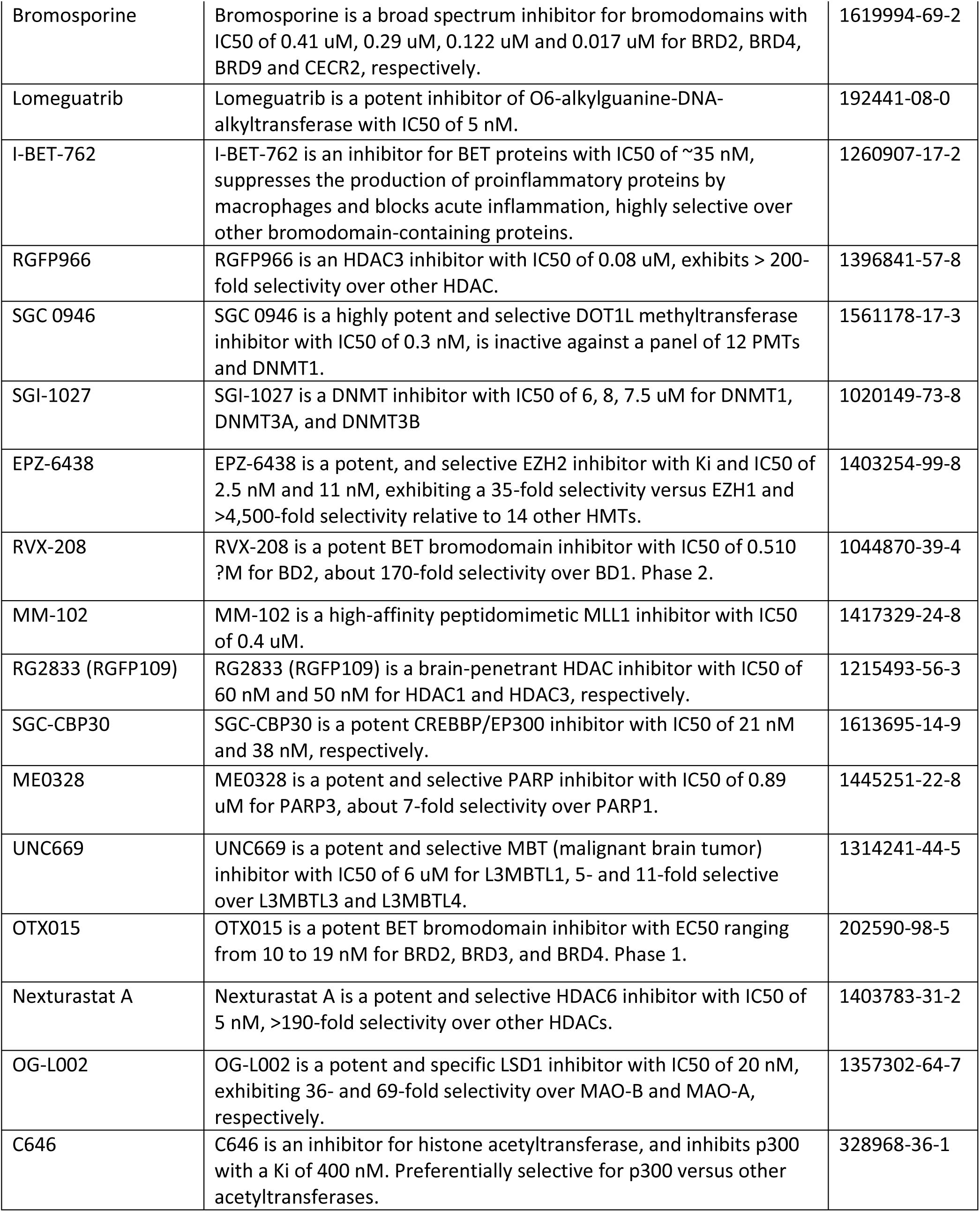
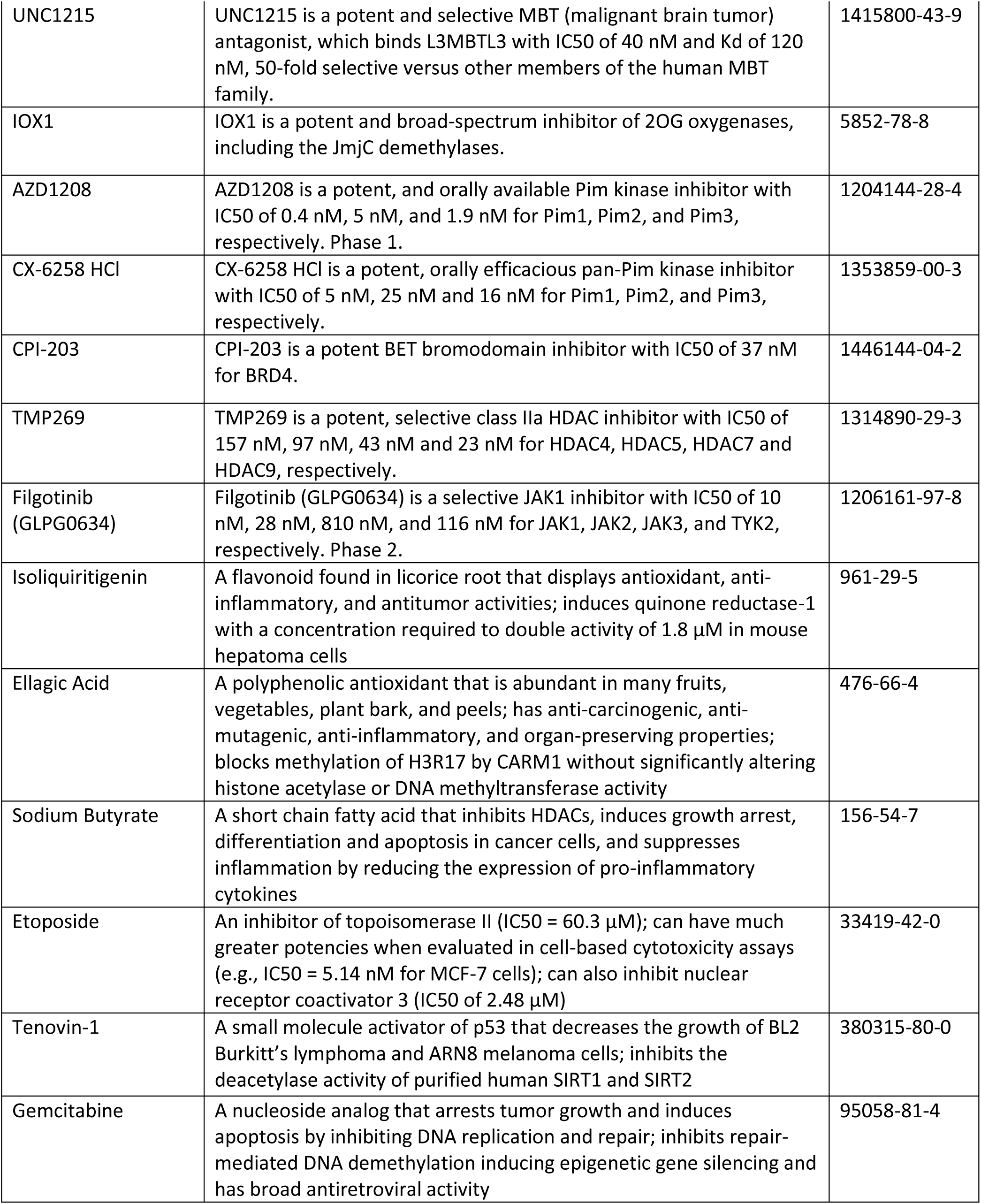
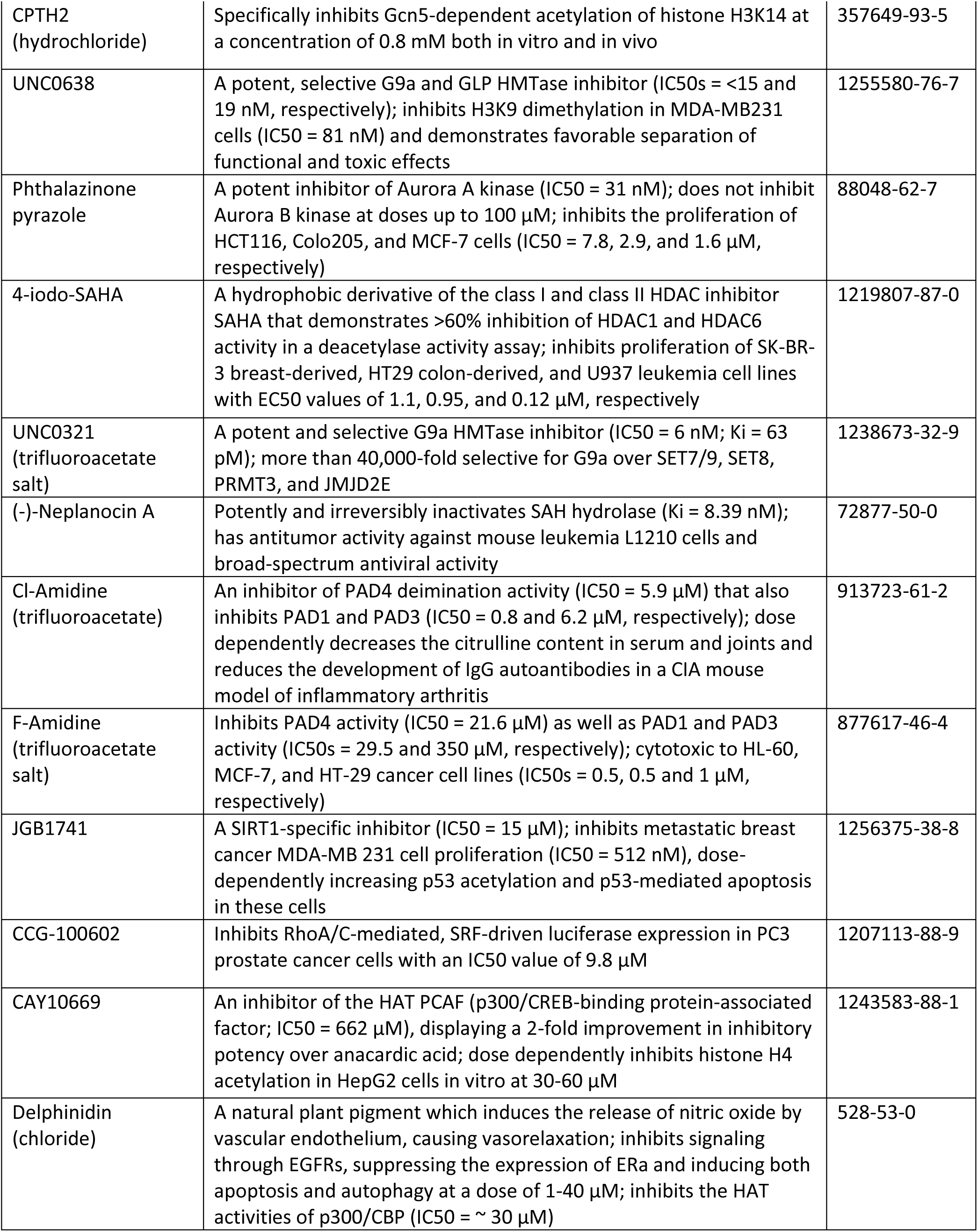
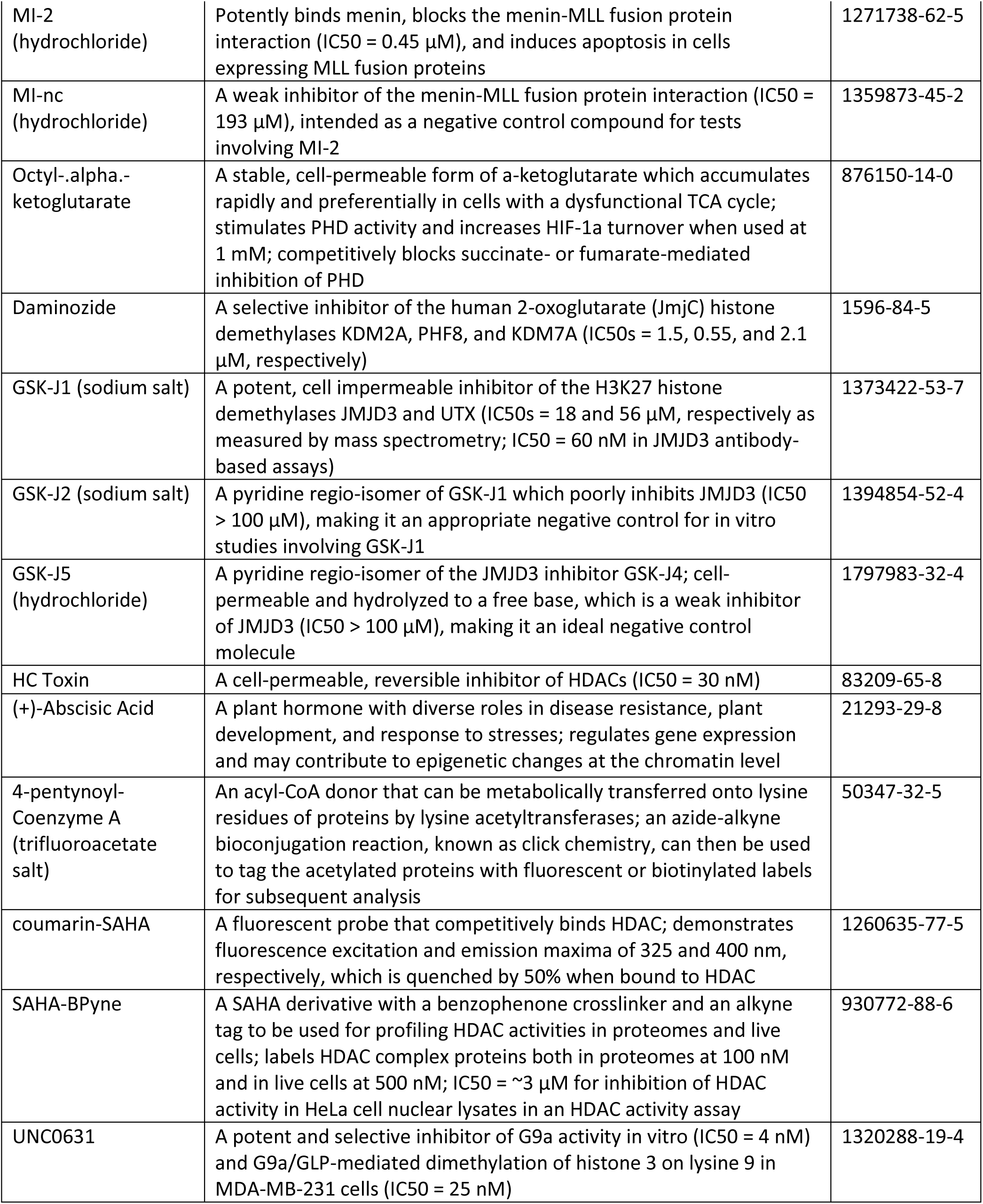
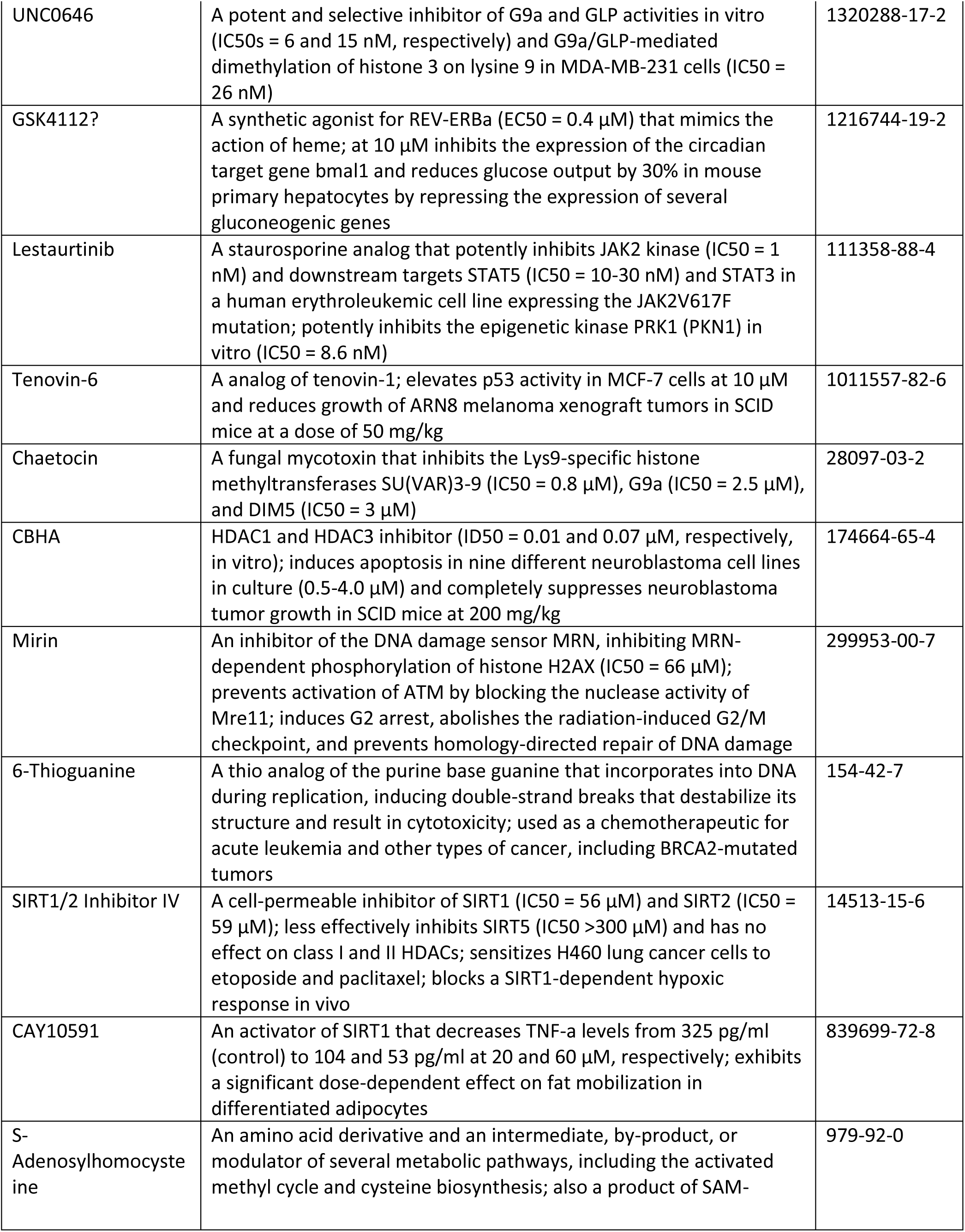
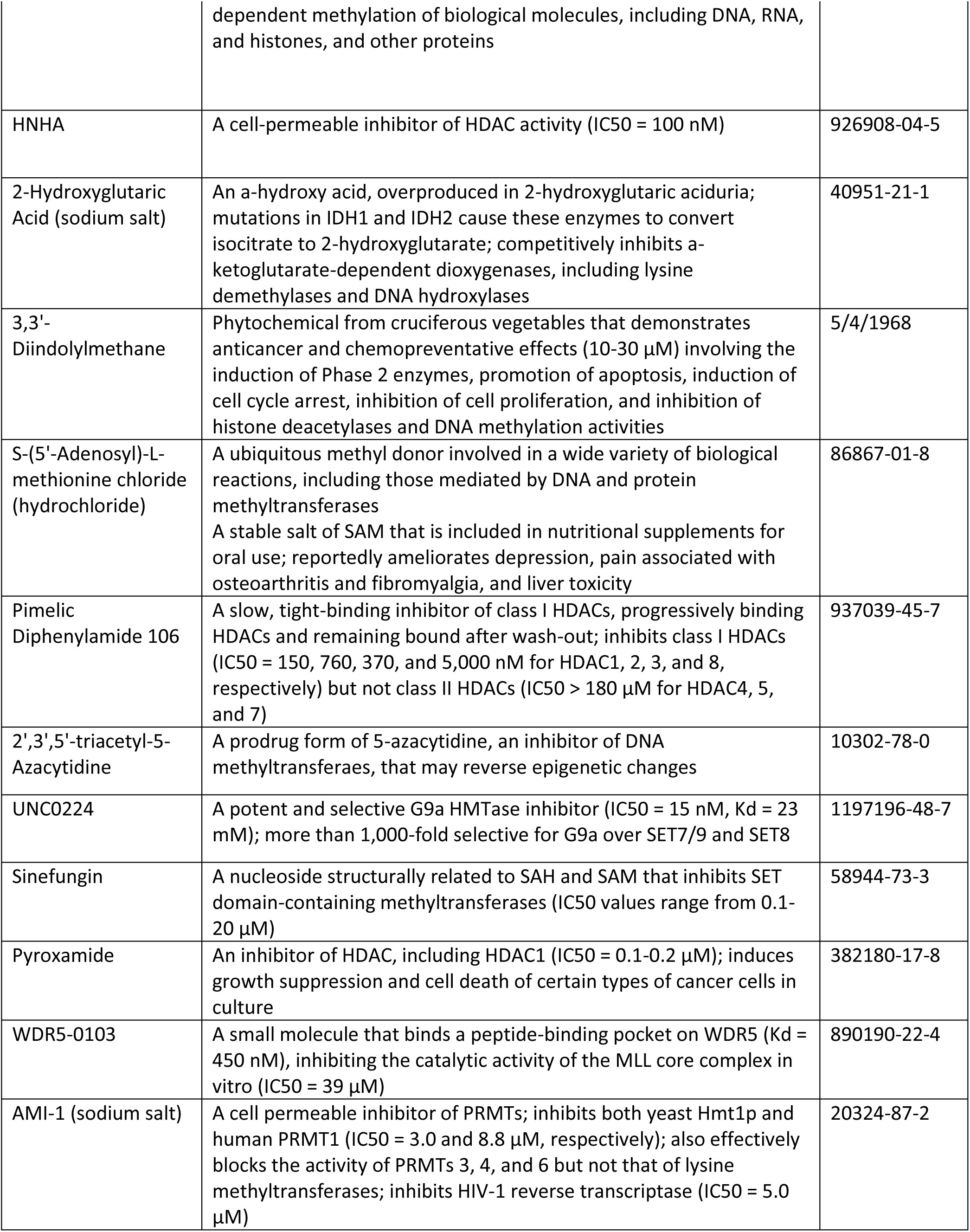
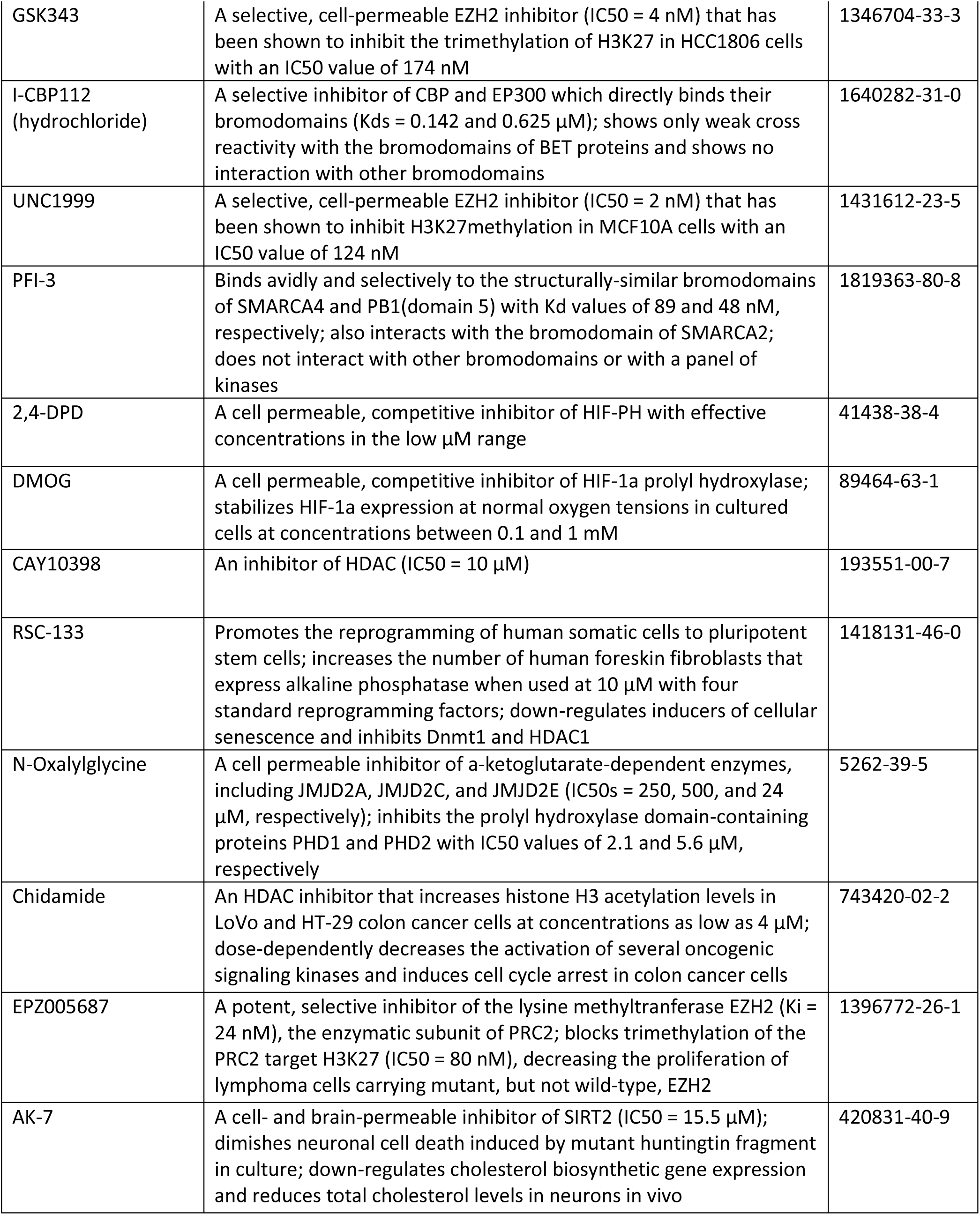
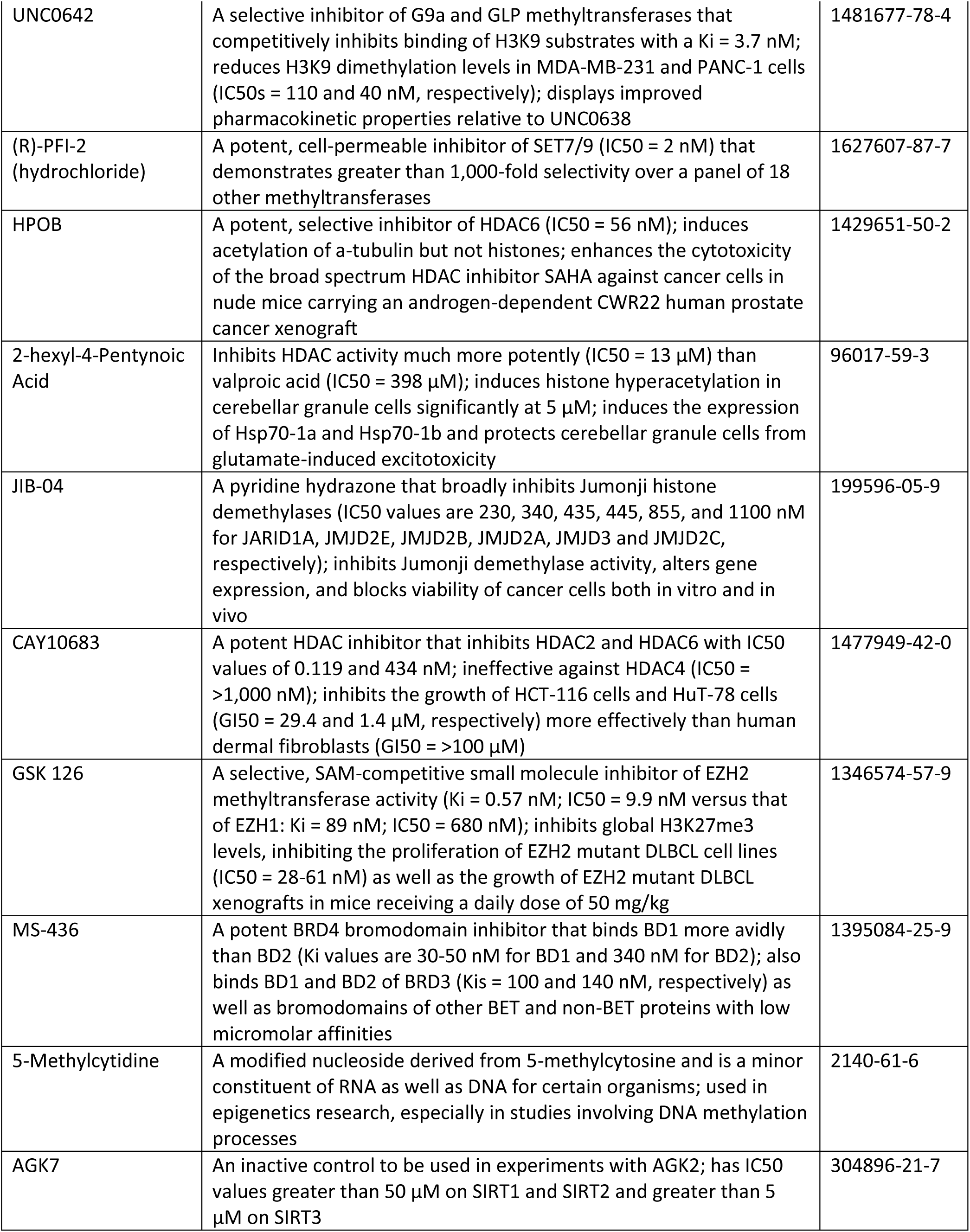
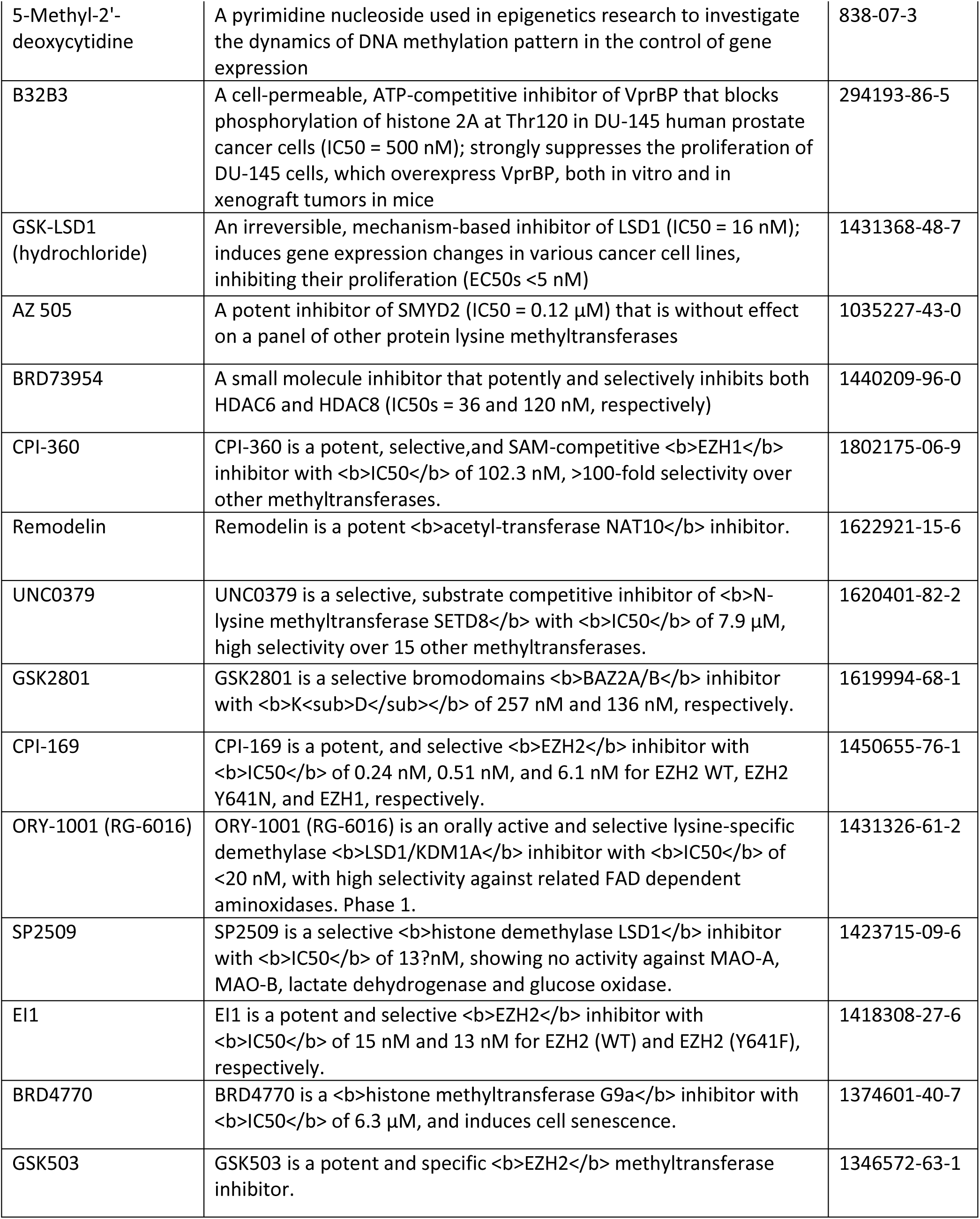
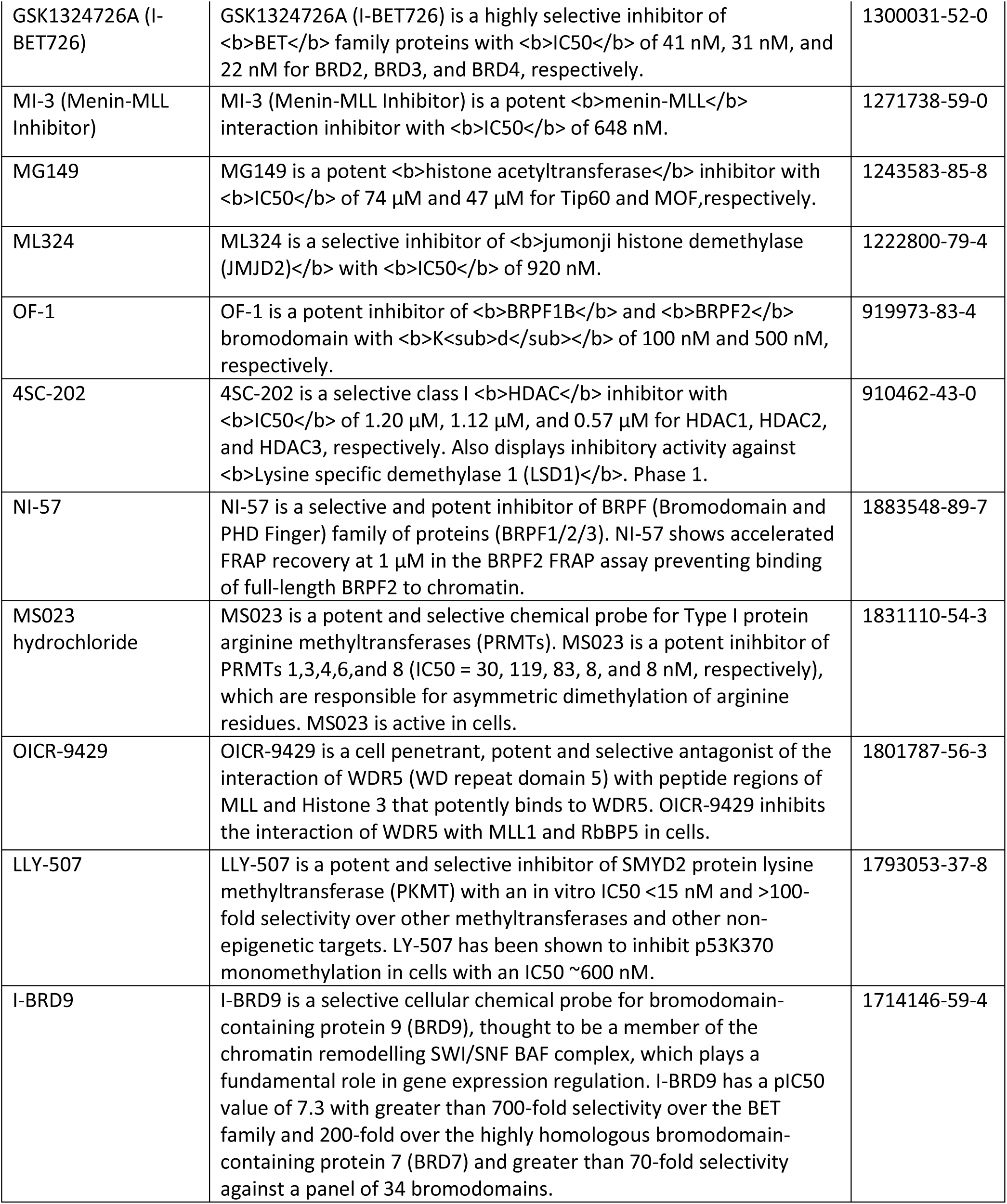
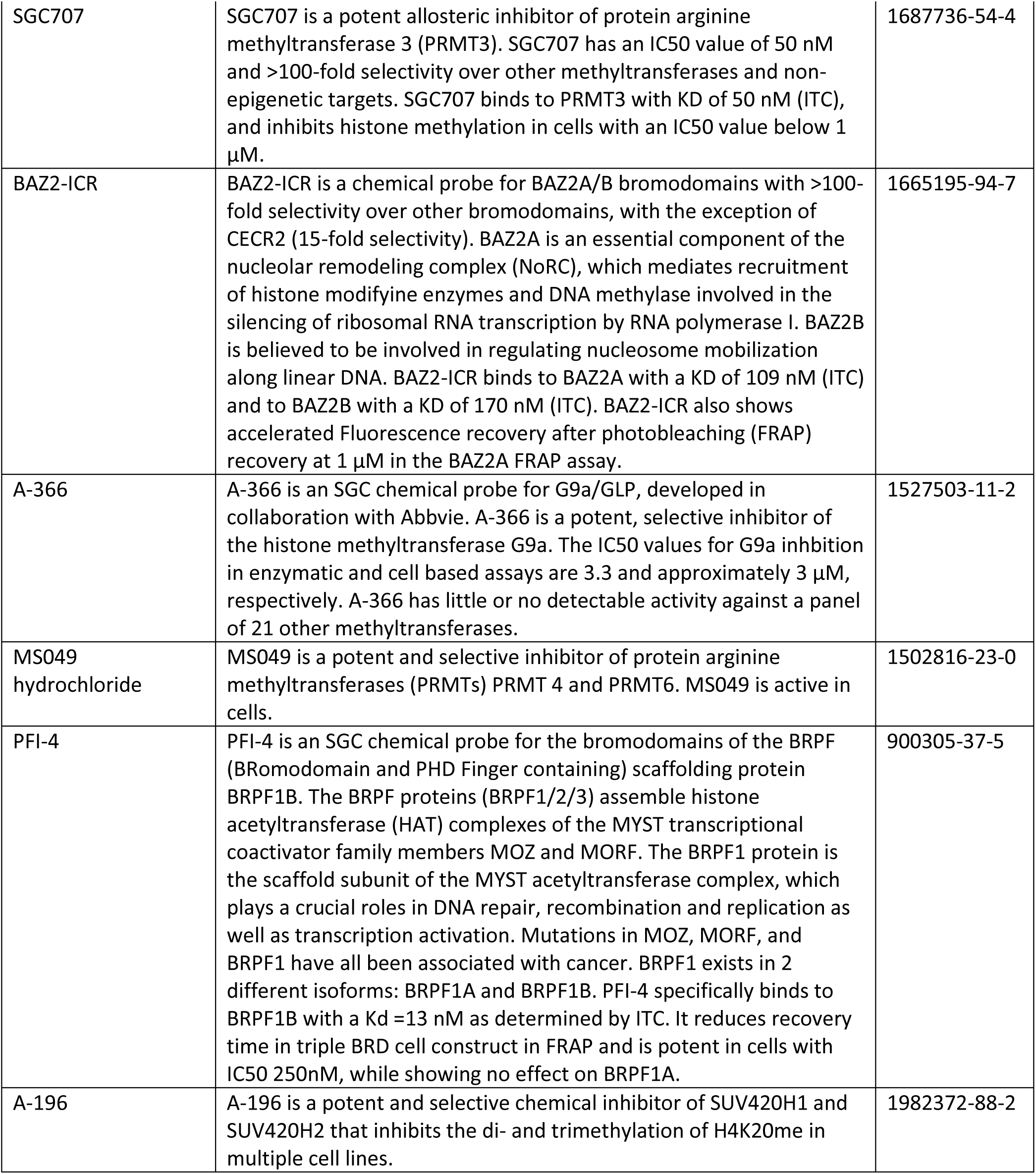
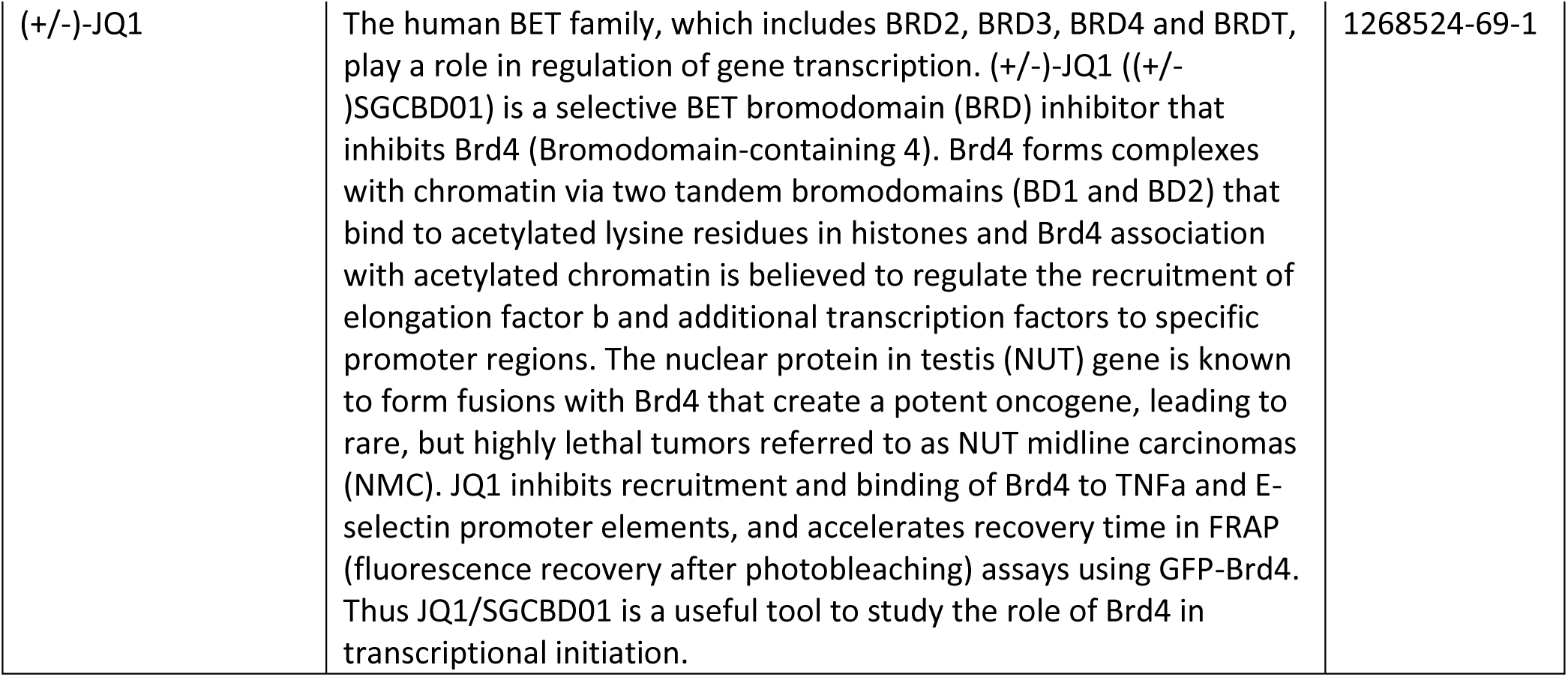
List of small molecule epigenetic modulators used to identify possible synergy with RNF5 knockdown AML cells.

**Supplemental Table S4.**
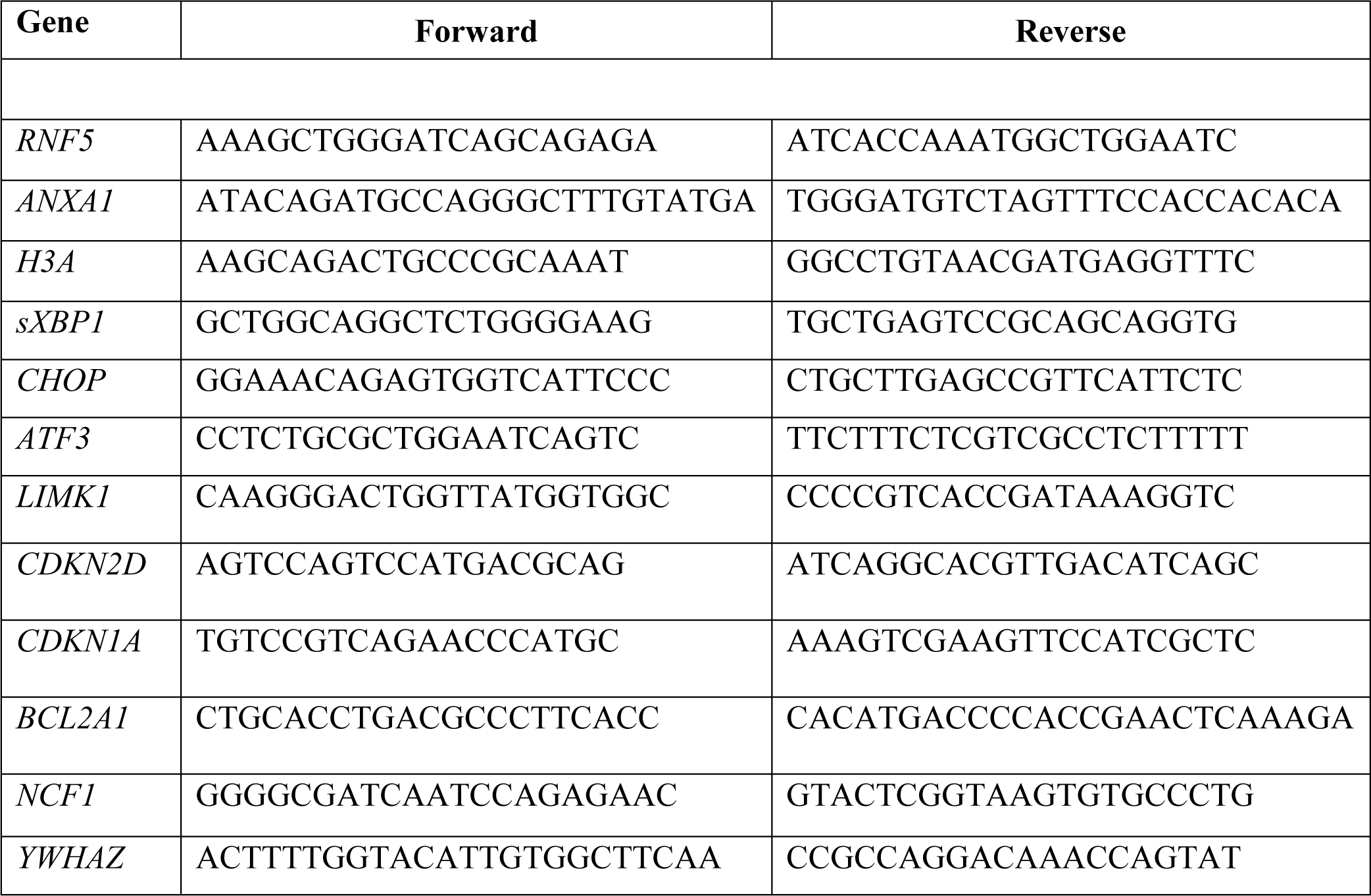
List of primers used for RT-qPCR analysis.

**Supplemental Table S5.**
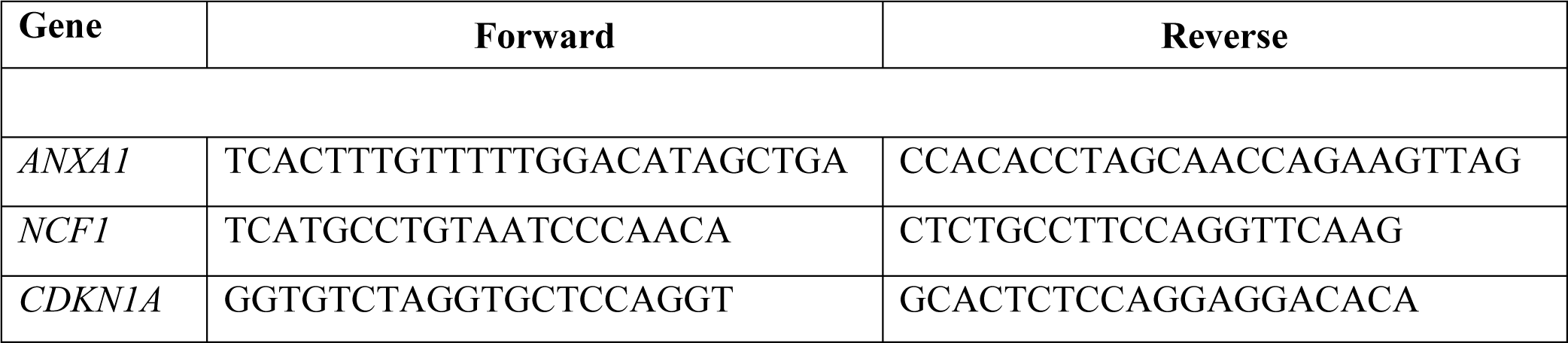
List of primers used for ChIP analysis.

## Notes

### Competing Interest Statement

There are no competing interest that are relevant to the work presented in this manuscript. ZAR is co-founder and serves as scientific advisor to Pangea Therapeutics. All other authors declare no competing interests.

